# Dual ATPase-activated DNA hemimethylation and cleavage by the MANTIS defence system

**DOI:** 10.64898/2026.09.23.753747

**Authors:** Steven J. Shaw, Sam C. Went, Aleksei Agapov, Joseph Westley, Anna Olina, Shirley Graham, Charlotte E. Chong, MultiDefence Consortium, Kate S. Baker, Malcolm F. White, Stineke van Houte, Edze R. Westra, Tim R. Blower, Mark D. Szczelkun

## Abstract

Bacterial defence systems are diverse and form major barriers to horizontal gene transfer and bacteriophage (phage) infection. Though restriction-modification systems were the first such defences discovered and remain the most widespread, the basis of self/non-self discrimination in many newly identified methylation-based systems remains unclear. Here, we define the mechanism of a widely distributed BREX-related defence family, previously termed Gao_RL, which we rename MANTIS (Methyltransferase, ATPases and Nuclease Targeting Inverted-repeat Sequences). Using phage infection and plasmid transformation assays in *Pseudomonas aeruginosa*, biochemical reconstitution, and structural biology, we show that MANTIS establishes self-identity through adenine hemimethylation of short asymmetric DNA sequences. Unexpectedly, a AAA+ ATPase (MtsA) and a co-regulator (MtsB) are both required to recruit and activate the methyltransferase MtsC, revealing a previously unknown ATP-dependent mode of epigenetic modification. DNA recognition by the MtsABC complex then triggers restriction by a dimeric helicase-nuclease effector (MtsD), with activity dependent on target-site architecture: inverted pairs of unmodified sites support robust restriction, whereas direct repeats or single sites are restricted less efficiently. Consistent with this model, virulent phages show strand-biased depletion of MANTIS target sites, indicating evolutionary pressure to evade restriction. Together, these findings reveal how dual ATPases couple self-modification to non-self restriction and provide a framework for understanding the expanding landscape of methylation-based defence systems.

**Graphical Abstract:** 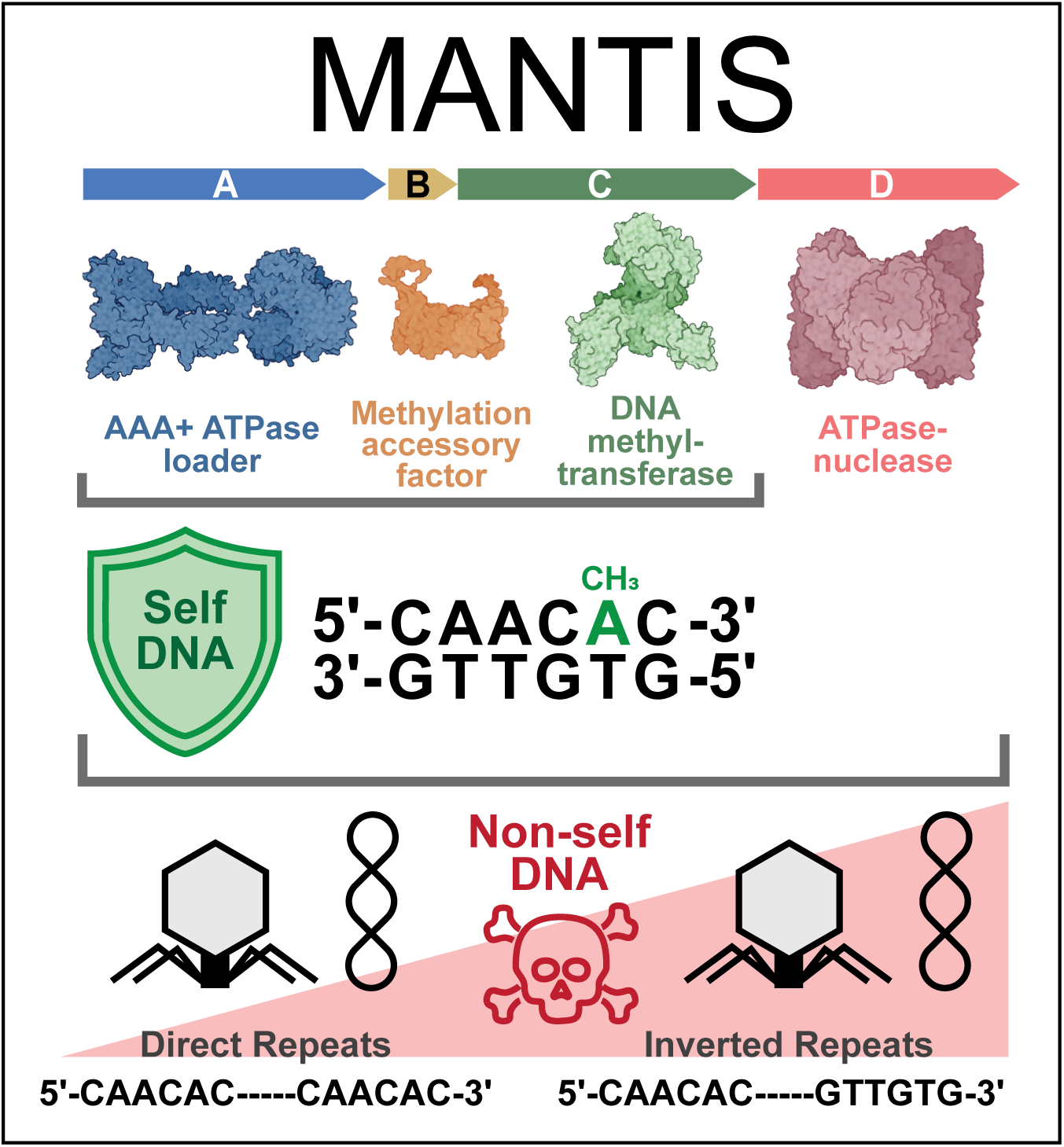

## INTRODUCTION

Bacteria are continually challenged by bacteriophages and other mobile genetic elements (MGEs), and a diverse array of defences have evolved to regulate this process. Recent systematic searches have uncovered a growing list of candidate defence systems^1–5^, but for many, mechanistic understanding remains limited. One example is the “restriction-like” Gao_RL (also termed Gao_29)^6^, which encodes features reminiscent of Bacteriophage Exclusion (BREX)-like ATPases and methylation-based self-marking^7^, but which was only analysed based on restriction activity with a subset of DNA phages. Here, we combine bioinformatics and phenotypic assays with biochemical, biophysical, and structural approaches to define how Gao_RL discriminates self from non-self DNA. Based on our findings, we reveal a new defence strategy and rename the system as MANTIS, for <u>M</u>ethyltransferase, <u>A</u>TPases, and <u>N</u>uclease <u>T</u>argeting of Inverted-repeat <u>S</u>equences, with individual components abbreviated Mts (**Fig. 1A**).

**Figure 1.**
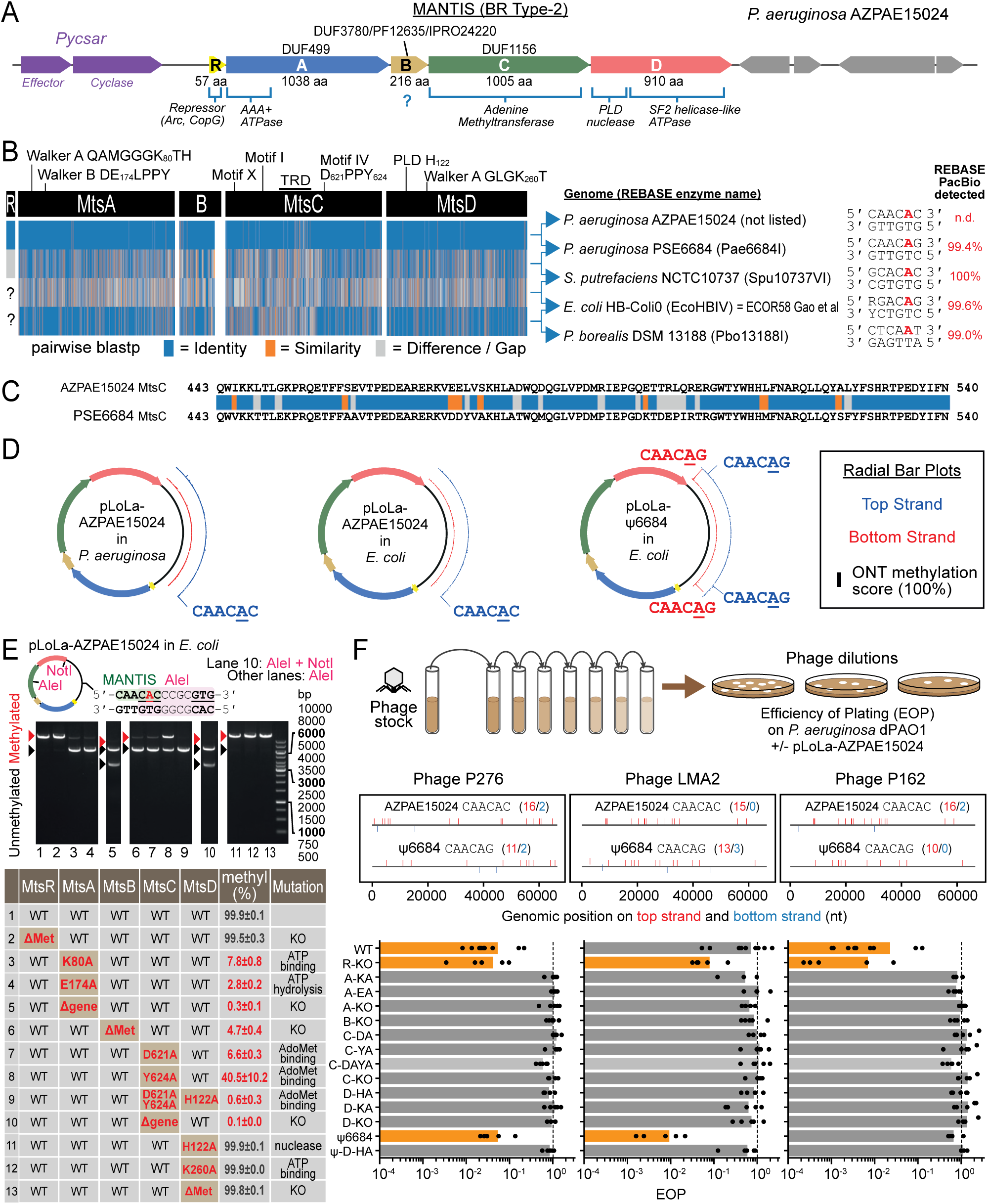
Protein requirements for DNA methylation and phage restriction by MANTIS. (**A**) Genomic region of *P. aeruginosa* AZPAE15024 showing the MANTIS BR-type-2 operon and encoded proteins/domains. (**B**) Pairwise BLASTP comparisons of MANTIS BR-type-2 systems. For Pae6684I, Spu10737VI, EcoHBIV and Pbo13188I, target sequences and methylation sites were validated by PacBio sequencing^17^. (**C**) Pairwise BLASTP comparison of the MtsC target-recognition domain (TRD) from AZPAE15024 and PSE6684. A chimeric variant (u6684) was generated by replacing this region in the AZPAE15024 TRD with that from PSE6684. (**D**) ONT methylation profiles corrected for background pLoLa methylation shown as radial bar charts (black bar = 100% methylation) for the top (blue) and bottom (red) strands of plasmids carrying the AZPAE15024 system grown in *P. aeruginosa* or *E. coli*, and the u6684 chimera grown in *E. coli*. (**E**) Effects of protein deletions and mutations on AZPAE15024 methylation. Plasmids contained a MANTIS site overlapping an AleI site, whose cleavage is blocked by MANTIS hemimethylation. Products from AleI or AleI + NotI digests were quantified following agarose gel electrophoresis. Values are means ± SD (n = 3). (**F**) EOP assay for phage restriction by MANTIS. Top, assay schematic; middle, distribution of MANTIS recognition sites on the top (red) and bottom (blue) strands of phages P276, LMA2 and P162; bottom, EOP on strains expressing wild-type or mutant MANTIS systems. Mutants are indicated by the Mts protein affected, followed by KO or the point mutation (see panel E). Bars show means and dots indicate biological replicates. Orange highlights restriction.

MANTIS can be classified as a Type-2 member of the DUF499-centred BREX-related (BR) systems that are widely distributed across over 3500 prokaryotic taxa^8^, aligning with BREX in their bacterial distribution but being more abundantly represented in archaea. BR-type 2 systems comprise four genes (**Fig. 1A**), encoding: MtsA, a multidomain protein with an N-terminal “ATPases Associated with diverse cellular Activities” (AAA+) domain of the ORC/Cdc6 clade; MtsB, a BR Type-2-specific small DUF3870 protein of unknown function; MtsC, an N6-adenine DNA methyltransferase (MTase); and MtsD, a multidomain protein including a metal-independent HKD/phospholipase D (PLD) nuclease and Superfamily 2 (SF2) helicase-like ATPase. We note that some MANTIS operons, including the *Pseudomonas* system analysed here, additionally encode MtsR, a putative repressor related to Arc/CopG.

DNA methylation is a widely-used epigenetic marker of self in bacterial defences, including Restriction-Modification (RM)^9^, BREX^7^, Methylation-Associated Defence System (MADS)^10^, and Defence Island System Associated with Restriction–Modification (DISARM)^11^. In classical RM, non-self DNA is targeted at specific sequences by a toxic nuclease, while methylation of those sites on the host genome blocks autoimmune cleavage. In MANTIS, MtsC is the putative protective MTase while MtsD is the candidate nuclease effector, with a domain architecture that resembles restriction enzymes that couple PLD nucleases to DNA-translocating helicase-like ATPase motors^12–14^. The second ATPase, MtsA, is a DUF499 protein likely to function as a structural scaffold and regulatory switch, analogous to its BrxC homologue in BREX^15^, which coordinates methylation and restriction activities via subunit and nucleotide interactions. MtsB is more enigmatic, but DUF3870 has been proposed to be a reader of DNA modification, potentially substituting for TUDOR-domains found in other BR systems^8^.

Here, we show that a *P. aeruginosa* MANTIS system coordinates dual ATPases during DNA modification and restriction. Methylation of just one strand of a 6-bp recognition site (“hemimethylation”) requires MtsA to load MtsC at specific sites, MtsB to stimulate methylation, and ATP hydrolysis by MtsA to enable catalytic turnover of methylation. Restriction of diverse phages and plasmids requires all four components (MtsA-D) and ATPase activities of both MtsA and MtsD. For both methylation and restriction, MtsC targets the recognition sequence. From analysis of phage and plasmid restriction, we propose a graded effector mechanism in which inverted pairs of unmodified recognition sites elicit strong restriction, necessary for protection against phages that produce costly infection, while direct repeats support weak restriction, nonetheless sufficient for protection against less virulent phages. Our MANTIS model provides important predictions for other BREX and BR types and more generally for systems that rely on DNA hemimethylation and/or ATPases.

## RESULTS

### MANTIS hemimethylates asymmetric 6 bp sequences

We analysed MANTIS from *P. aeruginosa* AZPAE15024 (**Fig. 1A**). To examine if MANTIS is a standalone defence or interacts with other defences, we considered the distribution of defence systems in 45 MANTIS-containing *P. aeruginosa* genomes (**Supplementary Figs. 1-2**)^16^. Though analysis accounting for phylogenetic structure did not identify any significant associations or dissociations between MANTIS and other defence systems (not shown), pairwise co-occurrence analysis revealed significant association of MANTIS with several defence systems (**Supplementary Fig. 2**). This suggests that MANTIS is a standalone defence system, with observed co-occurrences reflecting shared evolutionary history.

Given the genetic and domain architecture similarities between MtsC/MtsD and RM systems, we then searched REBASE^17^ for related examples that might guide our analysis. We identified closely-related proteins in *P. aeruginosa* PSE6684, annotated as an uncharacterised Type II RM system, Pae6684I (**Fig. 1B**). Though unannotated, the accompanying MtsR, MtsA and MtsB genes were also present, classifying Pae6684I as a putative MANTIS system. The Pae6684I recognition site had been assigned in REBASE using PacBio as 5’-CAACAG-3’, with the third adenine modified to give hemimethylated DNA. A wider REBASE search revealed further putative MANTIS systems, several with PacBio data supporting related hemimethylated targets, including an *Escherichia coli* variant with proteins >99% identical to those in Gao et al (**Fig. 1B**).

Amino acid sequence comparison revealed that the greatest divergence between the AZPAE15024 and Pae6684I proteins lay within amino acids (aa) 444–527 of MtsC, corresponding to part of a putative target recognition domain (TRD) (**Fig. 1B,C**), suggesting close but differing DNA sequence specificities. Using nanopore sequencing of plasmids carrying the full MANTIS operon in *P. aeruginosa* or *E. coli*, we confirmed the AZPAE15024 recognition site as 5’-CAACAC-3’, with hemimethylation at the third adenine and a single-bp change from Pae6684I (**Fig. 1D**). High MtsD conservation between AZPAE15024 and Pae6684I (99% amino acid identity, **Fig. 1B**), then implied that effector target recognition is likely mediated by MtsC, analogous to the MTases of Type III RM that load their helicase-nuclease. To test this, we replaced residues 445– 527 of AZPAE15024 MtsC with the corresponding region from Pae6684I to generate a chimeric system (u6684). Nanopore sequencing of u6684 showed the expected Pae6684I specificity (**Fig. 1D**), while the lack of detectable toxicity indicates that exchange of the MtsC TRD alone is sufficient to switch MtsD specificity; this is explored further below.

### DNA methylation requires MtsABC

Using the plasmid-based AZPAE15024 operon, we quantified DNA methylation in *E. coli* by measuring inhibition of *in vitro* DNA cleavage at an overlapping AleI restriction site (**Fig. 1E**). To assess roles for MANTIS proteins, we generated single-gene knockouts or mutations in key catalytic residues: the Walker A (K80A) or Walker B (E174A) motifs of the MtsA AAA+ ATPase; the DPPY motif of the MtsC Mtase (D621A and/or Y624A); the PLD nuclease motif of MtsD (H122A); or, the Walker A motif of the helicase-like ATPase domain of MtsD (K260A).

Knockout of *mtsC* was not toxic during transformation, despite the absence of protective methylation (**Fig. 1E**). In contrast, transformation with the MtsC double mutant (D621A/Y624A), which produced close to no methylation, was only possible when combined with an MtsD nuclease mutation. This supports a targeting hypothesis, as above: loss of MtsC prevents MtsD DNA recognition and thus toxicity, whereas a non-methylating but targeting-competent MtsC triggers autoimmunity via MtsD. Single MtsC mutants (D621A or Y624A) showed reduced but detectable methylation without evident toxicity. Residual MtsD activity must be toxic when methylation activity is completely lost as with the MtsC double mutant, but partial modification must be sufficient for protection.

Knockout of *mtsA* or *mtsB*, or mutation of the AAA+ ATPase motifs of MtsA significantly reduced DNA methylation (**Fig. 1E**) without detectable toxicity. In contrast, knockout or mutation of *mtsR* or *mtsD* had no effect on plasmid methylation (**Fig. 1E**), indicating these proteins are not required for DNA modification, at least using high copy-number plasmids.

Together, these results indicate that DNA methylation by MtsC requires MtsB and both ATP binding (Walker A) and hydrolysis (Walker B) activities of MtsA. The biochemical characteristics of DNA methylation are examined in detail below.

### Phage restriction requires MtsABCD

Hemimethylation of asymmetric 6-bp MANTIS sites creates a potential autoimmunity problem after replication when unmodified sites arise that could trigger cleavage. Type III and Type ISP RM systems resolve the same issue via “site-orientation preference”, in which ATP-dependent long-range communication between inverted site pairs on the same DNA is necessary for cleavage, leaving single sites and direct repeats resistant^18–21^. We reasoned that MANTIS similarly requires inverted site pairs to trigger restriction and predicted differences in efficiency of plating (EOP) between phages with or without such arrangements. To test this, we performed plaque assays on lawns of *P. aeruginosa* lacking native defences (dPAO1;^22^) using three virulent phages (P276, LMA2, and P162; **Fig. 1F**) with distinct AZPAE15024 and u6684 site configurations. We measured the EOP of the AZPAE15024 mutant panel (above), together with wild-type (wt) and MtsD nuclease-inactive (H122A) u6684 variants, relative to an empty vector control.

Phage P276, with sites for AZPAE15024 and u6684 on both strands and thus multiple inverted site pairs, was restricted by both wt MANTIS systems by 1-2 orders of magnitude (**Fig. 1F**). In contrast, phage without inverted sites were resistant to the cognate system: LMA2 was resistant to AZPAE15024 (all sites on the top strand in direct repeat) but was restricted ∼100-fold by u6684 (sites on both strands with multiple inverted site pairs); Conversely, P162 was restricted ∼10-fold by AZPAE15024 (sites on both strands with multiple inverted site pairs) but resistant to u6684 (all sites on the top strand in direct repeat). These results are consistent with restriction favouring inverted site repeats.

Deletion of MtsD (AZPAE15024), or mutation of its Walker A (AZPAE15024) or PLD nuclease (AZPAE15024 and u6684) motif/domain alleviated restriction of sensitive phages (**Fig. 1F**), consistent with a role for MtsD as an ATPase-dependent nuclease effector. In addition, all deletions and mutations of MtsA, -B and -C also alleviated restriction; this is consistent with the lack of toxicity noted above. We therefore propose that the effector activity of MtsD requires MtsA and its ATPase activity, MtsB, and DNA sequence recognition by MtsC. Restriction is tuned by the methylation activity of MtsC.

Mutation of *mtsR* to prevent its expression (AZPAE15024) marginally affected restriction of P276 and P162 but caused a pronounced (>10-fold) increase in LMA2 restriction (**Fig. 1F**). We propose that MtsR acts as a repressor of MANTIS activity, and alleviating this control permits cleavage of otherwise resistant direct-repeat sites. This suggests control of restriction by site orientation is not absolute as with Type III and ISP RM systems^18–21^.

### Inverted site repeats stimulate phage restriction by MANTIS

To further explore the recognition site requirements and breadth of MANTIS restriction activity, we performed 96-well format liquid culture infection assays using a defence system-free and prophage-free *P. aeruginosa* strain (viPAO1;^22^) carrying the wt AZPAE15024 plasmid and a panel of 60 virulent and 92 temperate *Pseudomonas* phages at a multiplicity of infection (MOI) of 0.1 (**Fig. 2A**, **Supplementary Table 1**). From changes in bacterial growth in the presence of each phage (**Supplementary Fig. 3**), we quantified the effect of MANTIS, which ranged from protective (positive values, indicating restriction) to costly (negative values, indicating reduced bacterial growth) (**Fig. 2A**).

**Figure 2.**
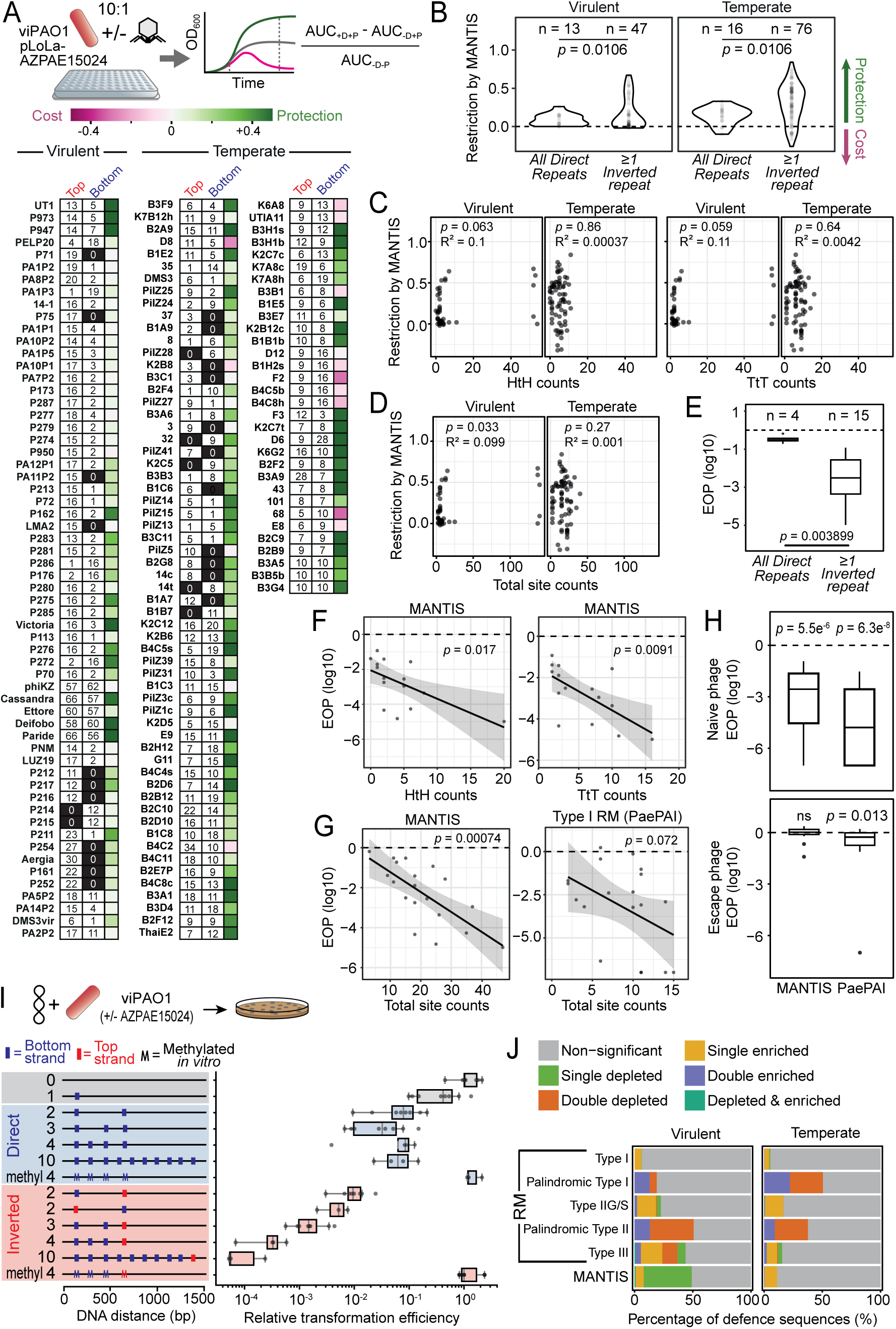
The number and orientation of MANTIS recognition sites influence restriction efficiency. (A) Liquid infection assay schematic (top) and summary of phages analysed, showing. MANTIS site distribution and the effect of MANTIS on host growth during infection (bottom). Colours indicate protection (green) or fitness cost (pink). **(B)** Restriction scores for virulent and temperate phages containing only direct-repeat sites or at least one inverted-repeat pair. Points represent phage means from 3 biological replicates; *n* indicates the number of phages analysed. **(C,D)** Relationships between restriction score and the number of inverted-repeat site pairs (**C**) or total MANTIS site count (**D**). *R*² and *p* denote Pearson correlation and significance, respectively. **(E)** Log,。 EOP of phages on MANTIS-expressing dAPO1 relative to a defenceless control for phages with direct-repeat sites only or at least one inverted-repeat pair. Points represent EOP means from 3 biological replicates; *n* indicates the number of phages analysed. **(F)** Relationship between log,。 EOP and the number of inverted-repeat site pairs. **(G)** Relationship between log,。 EOP and total site count for MANTIS and the Type I RM system PaePAI. **(H)** EOP of naïve and escape phages isolated from MANTIS-or PaePAI-expressing hosts. *p* indicates significance relative to zero. In (**E**) and (**H**), boxplots show median, IQR and 1.5×IQR whiskers; dashed line indicates equal plaque counts on defence and control lawns. **(I)** Relative transformation efficiency of plasmids differing in MANTIS site number, orientation and methylation status. Values were normalised to a defenceless control strain. **(J)** Depletion and enrichment of MANTIS and RM system recognition motifs (grouped by RM type, **Supplementary Data File 1**) across analysed phage genomes, calculated independently for top and bottom strands (**Supplementary** Fig. 5). Colours indicate non-significant, depleted, enriched, or mixed depletion/enrichment patterns.

Overall, MANTIS was not associated with detectable fitness costs in the absence of phage (**Supplementary Fig. 3A**) and provided effective protection against a broad range of phages (**Fig. 2A**); however, some phages remained resistant. To determine whether protection correlated with recognition-site architecture, we grouped virulent and temperate phages according to the presence or absence of inverted site-repeat pairs and compared scores (**Fig. 2B**). Regardless of phage lifestyle, MANTIS conferred stronger restriction of phages containing at least one pair of inverted sites. In virulent phages, restriction was strictly dependent on inverted sites, whereas temperate phages were still restricted when sites were arranged as direct repeats, although less efficiently than with inverted repeats (**Fig. 2B**). Overall, temperate phages were more susceptible to MANTIS. Ranking phages by virulence based on growth curves further showed that restriction of phages lacking inverted site repeats decreased significantly with increasing virulence, whereas the effect on phages containing inverted site repeats was only marginal (**Supplementary Fig. 4**). Together, these data indicate that MANTIS phenotypes depend on both phage genetics and life-history traits.

Restriction might be expected to scale with recognition site number. However, the liquid growth assay scores showed no significant relationship with head-to-head or tail-to-tail inverted site pair counts across either virulent or temperate phages (**Fig. 2C**). A borderline association with total (unpaired) site number was observed for virulent phages, but this effect was driven by three, non-nucleus forming, jumbo phages with unusually high site counts and was lost upon their data exclusion (**Fig. 2D**).

In contrast, plaque assays revealed a clearer dependence on site count. Measuring EOPs for 20 phages (**Supplementary Table 1**) on lawns expressing MANTIS AZPAE15024 or no defence, we found the expected significant dependence of EOPs on inverted sites (**Fig. 2E**) and a clear correlation with total (unpaired) site count (**Fig. 2F,G**), highlighting differences between assay conditions. In comparison, EOPs measured using a native Type I RM system (PaePAI) had only a weak correlation with recognition site count (**Fig. 2G**), highlighting differences between methylation-based defences.

Overall, restriction efficacy was similar between MANTIS and Type I RM but escape phages isolated from plaques showed distinct behaviours (**Fig. 2H**): MANTIS escape phages reached EOPs indistinguishable from no defence (∼1), consistent with efficient (likely methylation-mediated) escape; in contrast, RM escape phages remained significantly below 1, indicating partial escape but some restriction. Incomplete methylation by PaePAI has been observed^23^, which may be sufficient for escape but producing weaker PaePAI evasion upon re-infection.

### Inverted site repeats stimulate plasmid restriction by MANTIS

To test whether MANTIS provides generalist DNA-targeted immunity, we measured transformation efficiencies of viPAO1 carrying a chromosomally-integrated MANTIS AZPAE15024 operon using plasmids containing different numbers and arrangements of MANTIS sites (**Fig. 2I, Supplementary Table 1**). The observed restriction indicates that MANTIS activity does not require phage-specific components or replicative intermediates, and pre-methylation of plasmids conferred the expected complete protection.

A single MANTIS site reduced transformation efficiency, and the addition of a second site in direct repeat further increased restriction, whereas further direct repeats produced no significant additive effect (**Fig. 2I**). In contrast, inverted repeats enhanced restriction, with the effect scaling with site number. Together, these results mirror the phage data and indicate that while both direct and inverted repeats - and even a single site - can support restriction, inverted repeat pairs are markedly more effective. Thus, MANTIS has a graded site orientation preference mechanism, distinct to Type III and ISP RM systems that strictly require inverted repeats^18–21^.

### Phage genomes show strand bias in MANTIS sites

A common evolutionary strategy for MGEs to evade classical RM is recognition sequence loss, although the extent and mode of site avoidance vary with RM type. For enzymes recognising palindromic sequences (Type IIP RM), sites are strongly depleted in phages and plasmids^24–26^. In contrast, asymmetric sites recognised by Type I, IIG and III systems are not uniformly avoided^24^. For Type III RM, which triggers DNA cleavage using inverted site pairs, resistance can instead arise through selective depletion of sites on just one strand^27–29;^ for example, phage T7 has all 36 EcoP15I sites in direct repeat and is resistant.

Given the graded site orientation preference of MANTIS (**Fig. 1F, 2B,F**), and that some *Pseudomonas* phages contain exclusively direct repeat sites (**Fig. 2A**), we predicted that interactions with MANTIS may have driven phage evolution toward selection of protective directly-repeated sites, biasing depletion on one strand. To test this using a panel of phages (**Supplementary Table 1**), we collated strand-specific site counts for MANTIS AZPAE15024 and Pae6684I recognition sequences, together with recognition sequences for PacBio-validated and predicted *P. aeruginosa* RM systems from REBASE^17^ (**Supplementary Data File 1**), thereby allowing comparison of site depletion for MANTIS with systems with similar (RM Type III), and dissimilar (RM Type I & II) site orientation requirements: 38 Type I; 33 Type II and 10 Type III. Sites were further split into palindromic versus non-palindromic. Site counts for each strand were recalculated as standardised residuals relative to expected counts corrected for phage strand-specific nucleotide and triplet composition (**Supplementary Fig. 5**). Sites were then classified as enriched or depleted on each strand using 99% confidence interval thresholds (**Fig. 2J**).

Consistent with selective pressure to avoid inverted site pairs, ∼40% of virulent phages exhibited significant strand-specific single depletion of MANTIS sequences (**Fig. 2J**). The absence of single depletion in temperate phages may reflect reduced selective pressure, as targeting of direct repeats is more likely to block these phages (**Fig. 2B**). Single depletion was also observed for Type III RM sites, as predicted^27–29^. Although less frequent than for MANTIS, this occurred in both virulent and temperate phages, consistent with restriction solely at inverted site pairs^21^. MANTIS sites were additionally single enriched across phage lifestyles, as were asymmetric Type I, IIG/S, and III RM sites. Single enrichment may reflect a model where bacterial selection favours defences whose target sequences overlap essential, and therefore less mutable, phage regulatory regions^30^.

As observed in a previous study^24^, we found significant site avoidance (double depletion) of palindromic Type II RM sites, but not of asymmetric Type IIG/S sites (**Fig. 2J**). Extending these studies, we also detected double enrichment of these sites and double enrichment and depletion of palindromic Type I sites. Although a general elevated loss and gain of palindromic target sites (**Supplementary Fig. 5**) would be consistent with the hypothesis that DNA palindromes are intrinsically more mutagenic during replication^31^, this does not fully explain all our observations: we also detected double depletion of non-palindromic Type III sites in virulent phages, indicating that additional selective forces contribute to shaping RM site distributions.

### DNA methylation by MtsC requires DNA loading by MtsA

To further evaluate the DNA methylation mechanism, we purified AZPAE15024 MtsA-C proteins individually, and reconstituted methylation using plasmid DNA containing overlapping MANTIS and AleI sites (**Fig. 3A**). Under multiple turnover (steady state) conditions ([MtsC] < [DNA]) with ATP and the methyl donor *S*-adenosyl-L-methionine (AdoMet), methylation required all three proteins, consistent with the in-cell data (**Fig. 1E**), and was most efficient when MtsA and MtsB were present in excess over MtsC (**Fig. 3A**). Under single turnover conditions ([MtsC] > [DNA]), all three proteins remained essential and a 1:1:1 stoichiometry was sufficient for complete modification (**Supplementary Fig. 6**).

**Figure 3.**
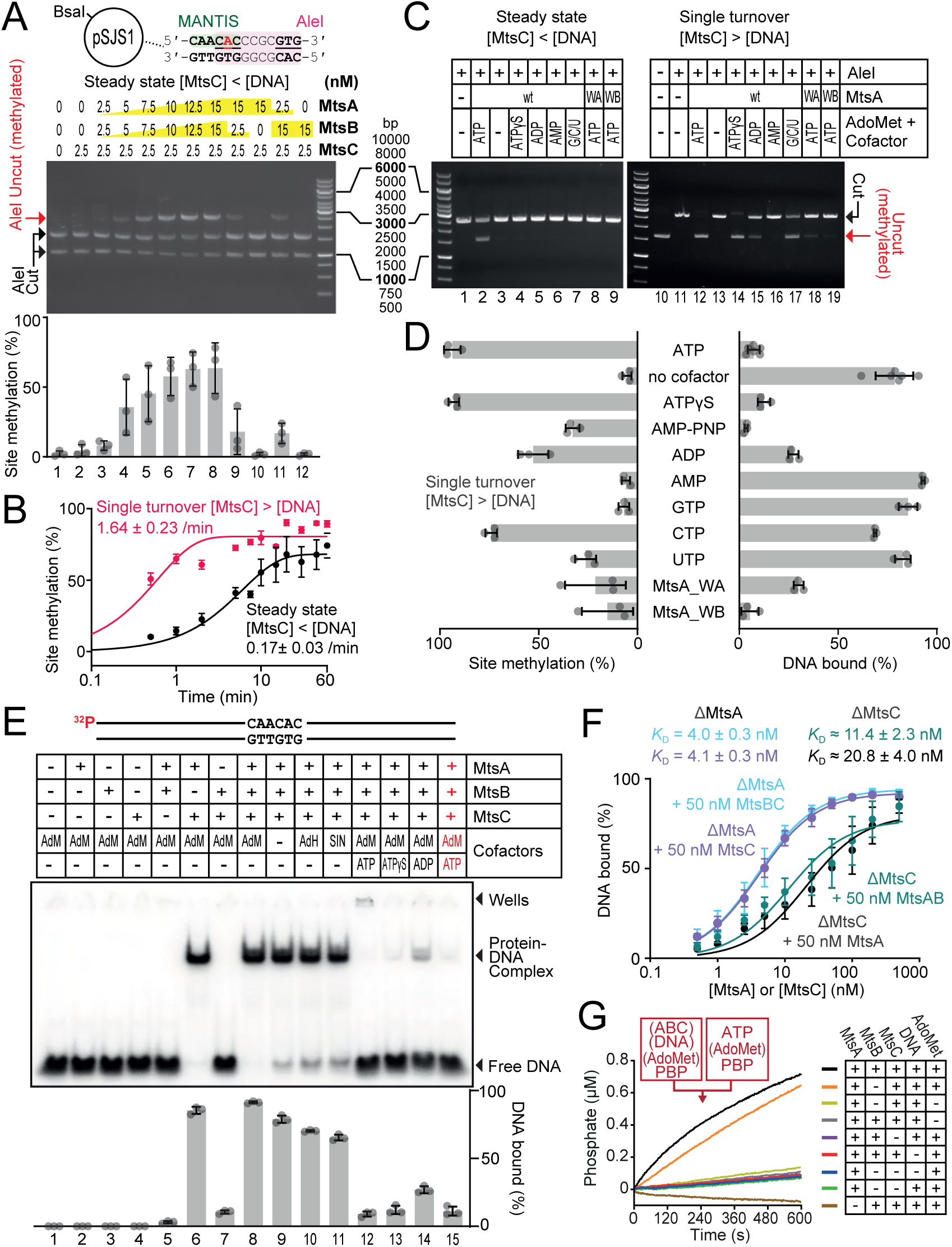
DNA methylation is ATP-dependent and requires DNA loading and activation of MtsC by MtsA and MtsB. **(A)** Representative agarose gel showing that MtsC-dependent DNA methylation requires both MtsA and MtsB. Methylation of pSJS1 following 30 min incubation with ATP and AdoMet was quantified by AleI/BsaI digestion. **(B)** Methylation time courses under (A:B:C:DNA = 6:6:1:2) and single-turnover (120:120:20:1) conditions using pSJS1. Rates were obtained by single-exponential fitting (mean ± SE, *N* = 3, (**Supplementary** Fig. 7)). **(C)** Representative agarose gel electrophoresis showing effects of nucleotide cofactors and MtsA Walker A (WA) and Walker B (WB) mutants on DNA methylation under steady-state and single-turnover conditions. Methylation was assessed after 30 min by AleI digestion. **(D)** DNA methylation inversely correlates with DNA binding. Methylation was measured by plasmid digestion (**Supplementary** Fig. 8) and DNA binding by EMSA (**Supplementary** Fig. 12) (mean ± SD, *N* = 3). **(E)** Representative EMSA showing that DNA binding by MtsC requires MtsA. DNA-protein complexes were analysed by EMSA and quantified (mean ± SD, *N* = 3). In lanes 2–14, DNA and protein were mixed before cofactor addition. Lane 15 is equivalent to lane 12, except that ATP was added 5 min before protein addition. **(F)** Binding affinities of MtsA, MtsB and MtsC determined by EMSA titrations. Dissociation constants were obtained by fitting a one-site binding isotherm (mean ± SE, *N* = 3). **(G)** Stopped-flow phosphate-binding protein (PBP) assay showing ATPase activity of MtsA under the indicated conditions (Average of *N* = 2).

Under steady state conditions, DNA methylation proceeded relatively slowly (**Fig. 3B**, **Supplementary Fig. 7**) and was exclusively ATP-hydrolysis dependent and abolished by Walker A or B mutations in MtsA (**Fig. 3C**). In contrast, ATP-dependent methylation was much faster under single turnover conditions (**Fig. 3B**, **Supplementary Fig. 7**), and was also supported by ATP analogues, and to a lesser extent by ADP or CTP, yet remained blocked by Walker A and B mutations in MtsA (**Fig. 3C,D; Supplementary Fig. 8**). Together, these data indicate that nucleotide binding by MtsA is sufficient to trigger a single round of methylation, whereas ATP hydrolysis is required for “turnover” to methylate another DNA site.

To explore DNA association by MtsC, we separated free and bound species of a 56 bp oligoduplex with a single, central unmethylated site using an electrophoretic mobility shift assay (EMSA) (**Fig. 3E**). Unexpectedly, we did not observe shifted DNA species with MtsA, MtsB or MtsC alone. A protein-DNA complex was observed if MtsA and MtsC were combined; addition of MtsB had no noticeable effect (**Fig. 3F**, **Supplementary Fig. 9**), and only moderate enhancement was observed with AdoMet or other product/inhibitor cofactors (**Fig. 3E**). We identified both MtsA and MtsC, but not MtsB, in the shifted band (**Supplementary Fig. 10**). DNA-bound species were not observed using pre-methylated DNA (**Supplementary Fig. 11**), indicating that the MtsC loading is specific to unmodified sites and that MtsB binding does not appear to be a modification reader.

When ATP or other nucleotides that support single turnover methylation were included in EMSAs, predominantly free DNA was detected (**Fig. 3E**). This outcome was independent of whether ATP was added before MtsA/C or after MtsC-DNA complex formation (compare lanes 12 and 15). We next used EMSAs to quantify MtsC-DNA formation with MtsA or MtsA mutants, using different nucleotides (**Fig. 3D; Supplementary Fig. 12**). Except for the MtsA mutants and CTP, conditions that caused MtsC dissociation correlated with increased DNA methylation (**Supplementary Fig. 13**). The lack of correlation with the ATPase mutants of MtsA may reflect that the coupling between the proteins is disrupted.

Negligible ATP hydrolysis by MtsA is observed in the absence of DNA, MtsC or AdoMet (**Fig. 3G**). Once the DNA binding complex is formed as demonstrated in **Fig. 3E** by combining DNA with MtsA/C, further stabilised by the inclusion of AdoMet, ATPase stimulation is observed. A second, earlier burst is present with the addition of MtsB, conditions that are required for methylation (**Fig. 3A**), which could be a reconfiguration event that is not visible in the EMSA data. Overall, ATP consumption was comparatively slow, supporting a role for MtsA as a molecular switch operating through a single transition rather than as a motor requiring multiple catalytic cycles.

Together, these results suggest that MtsA and MtsC cooperate in site-specific DNA binding. Favourable nucleotide binding by MtsA and the presence of MtsB additionally promotes DNA methylation by MtsC, followed by DNA dissociation that may be triggered by site methylation. While ATP binding can activate methylation, hydrolysis stimulated by MtsBC and DNA is necessary for multiple turnover (**Fig. 3C**). The role of nucleotide hydrolysis may be to “reset” the MtsA conformation following the MtsC loading and MtsB-stimulated methylation event.

### Transient MtsA and MtsB interactions load MtsC with DNA

To further examine interactions between MANTIS proteins, we first determined their individual solution oligomeric states using analytical size-exclusion chromatography (SEC) and mass photometry (**Supplementary Fig. 14A,B**): MtsA was predominantly dimeric but also showed evidence of a higher-order species of unknown stoichiometry; MtsB formed a stable dimer; MtsC behaved as a monomer, consistent with recognition and hemimethylation of an asymmetric target site; and, MtsD formed a stable dimer.

To explore the methylation reaction, we mixed MtsA-C along with combinations of dsDNA, nucleotide and cofactor analogues and separated the resulting species using SEC alongside MtsA as a control. When MtsA-C were combined with a 46 bp oligonucleotide in the presence of ATPɣS and sinefungin, an analogue of AdoMet and inhibitor of methyltransferases, the MtsC peak shifted to a lower elution volume and displayed a markedly higher 260 nm absorbance (**Supplementary Fig. 14C**). Although suggestive of MtsC being loaded onto DNA, higher-order MANTIS complexes were not observed by either SEC or mass photometry, despite extensive variations of conditions, cofactors, MANTIS components and DNA lengths. The MtsA-MtsC-DNA complexes inferred from EMSAs may be weak and only stablished within the gel matrix. As a result, to better understand their functions, we solved the structures of each MANTIS protein in isolation.

#### MtsA structure

Cryo-electron microscopy (Cryo-EM) analysis of MtsA showed an asymmetric dimer interface between N-terminal ATPase domains (aa 1-480) (**Fig. 4A**, **Table 1**; **Supplementary Figs. 15-17**). Due to their flexibility, the C-terminal “tails” were unresolved. The AAA+ ATPase domains and a winged helix-turn-helix-like (wHTH-like) subdomain align to equivalent domains in BrxC (PDB: 9ZDX)^15^ (**Supplementary Fig. 16A**). Residues 1–249 form the large, RecA-like α/β subdomain of a AAA+ ATPase, terminating with the fifth β-strand and the trans-acting arginine-finger (R244) region. Residues 250–350 form a small α-helical AAA+ subdomain containing Sensor 2 (R336). Residues 351–475 form the wHTH-like subdomain. Though MtsA and BrxC have similar N-terminal ATPase domains, AlphaFold3 models of the respective tail regions, also unresolved for BrxC, differ more substantially (**Supplementary Fig. 16B**).

**Figure 4.**
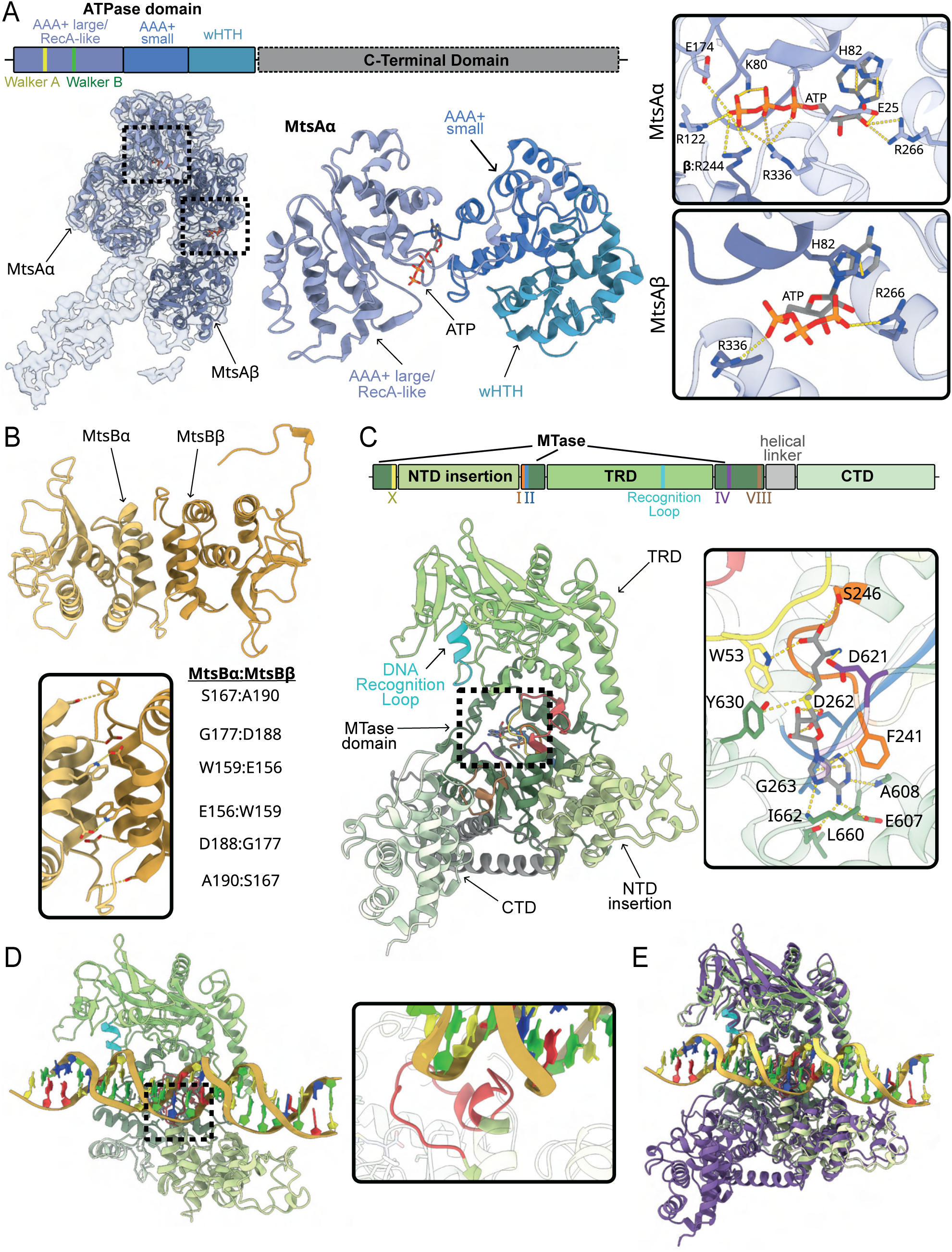
Structural analysis of MtsA, MtsB and MtsC. (**A**) Cryo-electron microscopy structure of the MtsA ATPase-domain dimer at 4 Å resolution with bound ATPs (other views in Supplementary Fig. 14). The domain map shows resolved regions of MtsA in blue together with the positions of the Walker A and Walker B motifs, and unresolved regions in grey (see Supplementary Fig. 15A). Inset: Zoomed view of ATP binding in the MtsAα and MtsAβ protomers (yellow dashed lines indicate ligand–protein interactions identified by PoseView; Supplementary Fig. 15A). (**B**) Crystal structure of the MtsB dimer at 2.6 Å resolution. Inset: Residues contributing to the dimerization interface (yellow dashed lines indicate amino acid interactions identified by PDBePISA). (**C**) Crystal structure of the MtsC monomer bound to AdoMet (grey) at 1.7 Å resolution with major domains, features, and motifs coloured according to the domain map. The DNA recognition loop (cyan) was identified using the MtsC chimera in Fig. 1. A DNA-binding loop shown in panel D is highlighted in red. Inset: Position of AdoMet relative to Motif I (FxGxG) and Motif IV (DPPY). (**D**) Cryo-electron microscopy structure of the MtsC monomer bound to a 46 bp Oligoduplex at 2.8 Å (Supplementary Fig. 14) with the DNA recognition loop (cyan) and DNA-binding loop (red) shown. Note that the CTD is not resolved. Inset: Zoomed view of the DNA-binding loop positioned within the DNA minor groove. (**E**) Structural alignment of MtsC–AdoMet (purple) and MtsC–DNA (green shades).

**Table 1.**
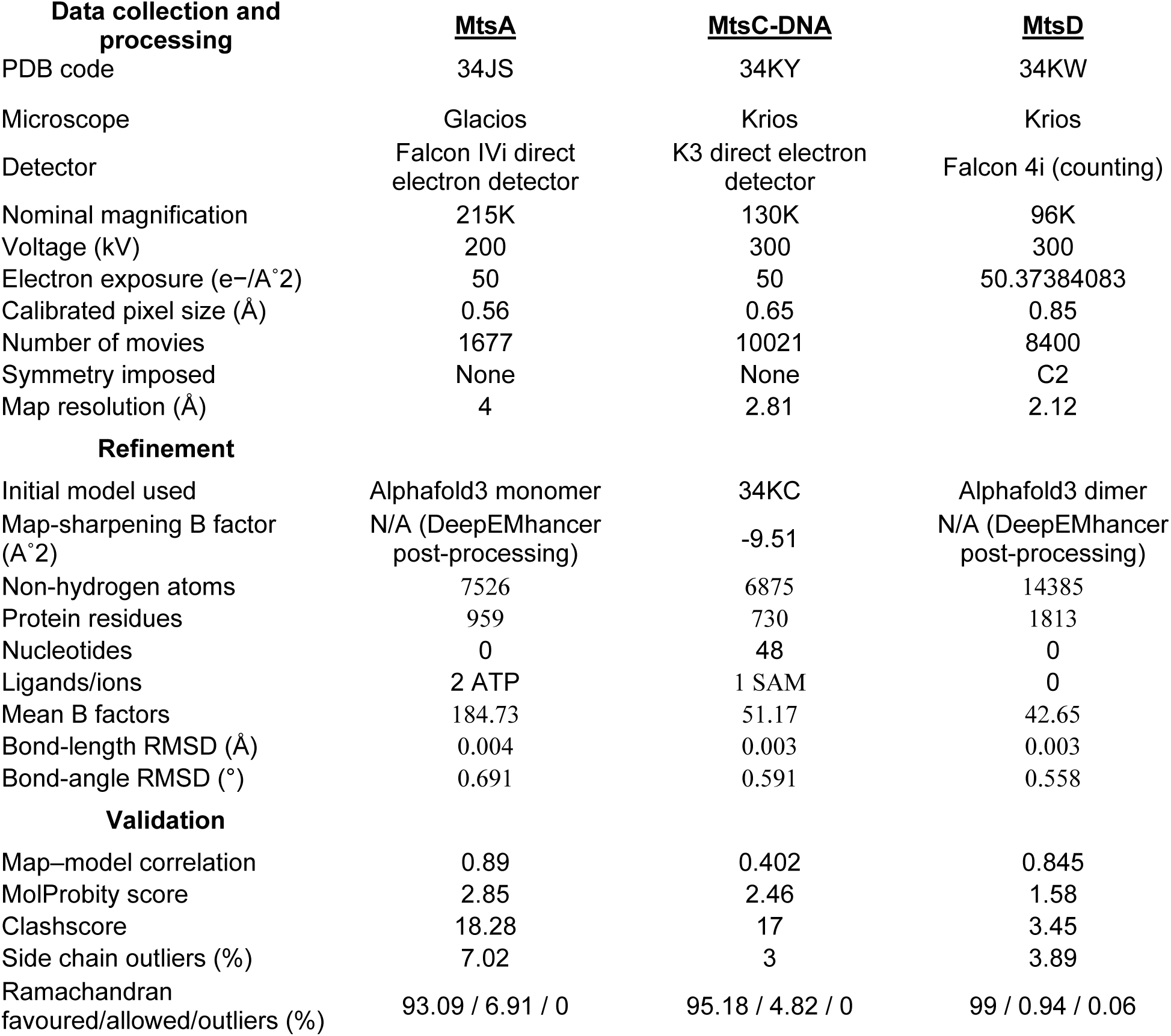
Cryo-EM data collection and refinement statistics.

| **Data collection and processing** | **<u>MtsA</u>** | **<u>MtsC-DNA</u>** | **<u>MtsD</u>** |
| --- | --- | --- | --- |
| PDB code | 34JS | 34KY | 34KW |
| Microscope | Glacios | Krios | Krios |
| Detector | Falcon IVi direct electron detector | K3 direct electron detector | Falcon 4i (counting) |
| Nominal magnification | 215K | 130K | 96K |
| Voltage (kV) | 200 | 300 | 300 |
| Electron exposure (e-/Å<sup>2</sup>) | 50 | 50 | 50.37384083 |
| Calibrated pixel size (Å) | 0.56 | 0.65 | 0.85 |
| Number of movies | 1677 | 10021 | 8400 |
| Symmetry imposed | None | None | C2 |
| Map resolution (Å) | 4 | 2.81 | 2.12 |
| **Refinement** |  |  |  |
| Initial model used | AlphaFold3 monomer | 34KC | AlphaFold3 dimer |
| Map-sharpening B factor (Å<sup>2</sup>) | N/A (DeepEMhancer post-processing) | -9.51 | N/A (DeepEMhancer post-processing) |
| Non-hydrogen atoms | 7526 | 6875 | 14385 |
| Protein residues | 959 | 730 | 1813 |
| Nucleotides | 0 | 48 | 0 |
| Ligands/ions | 2 ATP | 1 SAM | 0 |
| Mean B factors | 184.73 | 51.17 | 42.65 |
| Bond-length RMSD (Å) | 0.004 | 0.003 | 0.003 |
| Bond-angle RMSD (°) | 0.691 | 0.591 | 0.558 |
| **Validation** |  |  |  |
| Map-model correlation | 0.89 | 0.402 | 0.845 |
| MolProbity score | 2.85 | 2.46 | 1.58 |
| Clashscore | 18.28 | 17 | 3.45 |
| Side chain outliers (%) | 7.02 | 3 | 3.89 |
| Ramachandran favoured/allowed/outliers (%) | 93.09 / 6.91 / 0 | 95.18 / 4.82 / 0 | 99 / 0.94 / 0.06 |

MtsA dimers form an interface equivalent to that of BrxC dimers^15^. Despite the modest overall cryo-EM resolution (4 Å), DeepEMhancer sharpening revealed density for ATP and nearby sidechains in both MtsA protomers (**Fig. 4A; Supplementary Fig. 17**). In both sites, ATP is anchored by base stacking with H82. In the first protomer, MtsAα, ATP binds at the interface with the second protomer, MtsAβ, with the γ-phosphate extending towards the arginine finger of MtsAβ. The ribose is stabilised by E25 within loop 15–27 of MtsAα, which is displaced inward through contacts with MtsAβ (helix 229–245 and loop 62–69). In contrast, the ATP in MtsAβ is solvent-exposed because it lacks an equivalent protomer contact. Consequently, loop 15–27 adopts a different conformation, the arginine finger interaction is absent, and the ATP molecule occupies an alternative pose (**Fig. 4A; Supplementary Fig. 17B,C**).

Most AAA+ proteins oligomerise into ring- or spiral-shaped assemblies in which protomer interfaces create multiple nucleotide-binding pockets that can produce sequential waves of NTP hydrolysis^32^. In contrast, the dimer interfaces of both MtsA and BrxC appear to support only a single ATPase-competent active site^15^ (**Supplementary Fig. 17B**). Notably, BrxC can form heptameric rings in the presence of DNA, a property not observed for MtsA. Additionally, BrxC dimers can associate with one or two PglZ–BrxB dimers via an inactive AAA+ domain of BrxB, enhancing both the ATPase and DNA-binding activities of BrxC. The activation of the MtsA ATPase by MtsB, MtsC and DNA (**Fig. 3G**) is suggestive of productive interactions, but the detailed mechanism remains to be determined.

#### MtsB structure

The structure of dimeric MtsB was solved to 2.6 Å resolution by X-ray crystallography (**Fig. 4B**, **Table 2**). The dimer interface is stabilised by three reciprocal hydrogen-bond pairs (W159–E156, S167–A190, and G177–D188) that are conserved among MtsB homologues (**Supplementary Fig. 18A**). When tested, disruption of this interface by targeted mutagenesis prevented transformation with the complete system, suggesting that dimerization is required for host methylation. MtsB does not contain any predicted functional domains and lacks close structural homologues (best DALI Z-score = 3.5)^33^. Its strongly positive surface (**Supplementary Fig. 18B**) may facilitate protein and/or DNA interactions within transient complexes.

**Table 2.**
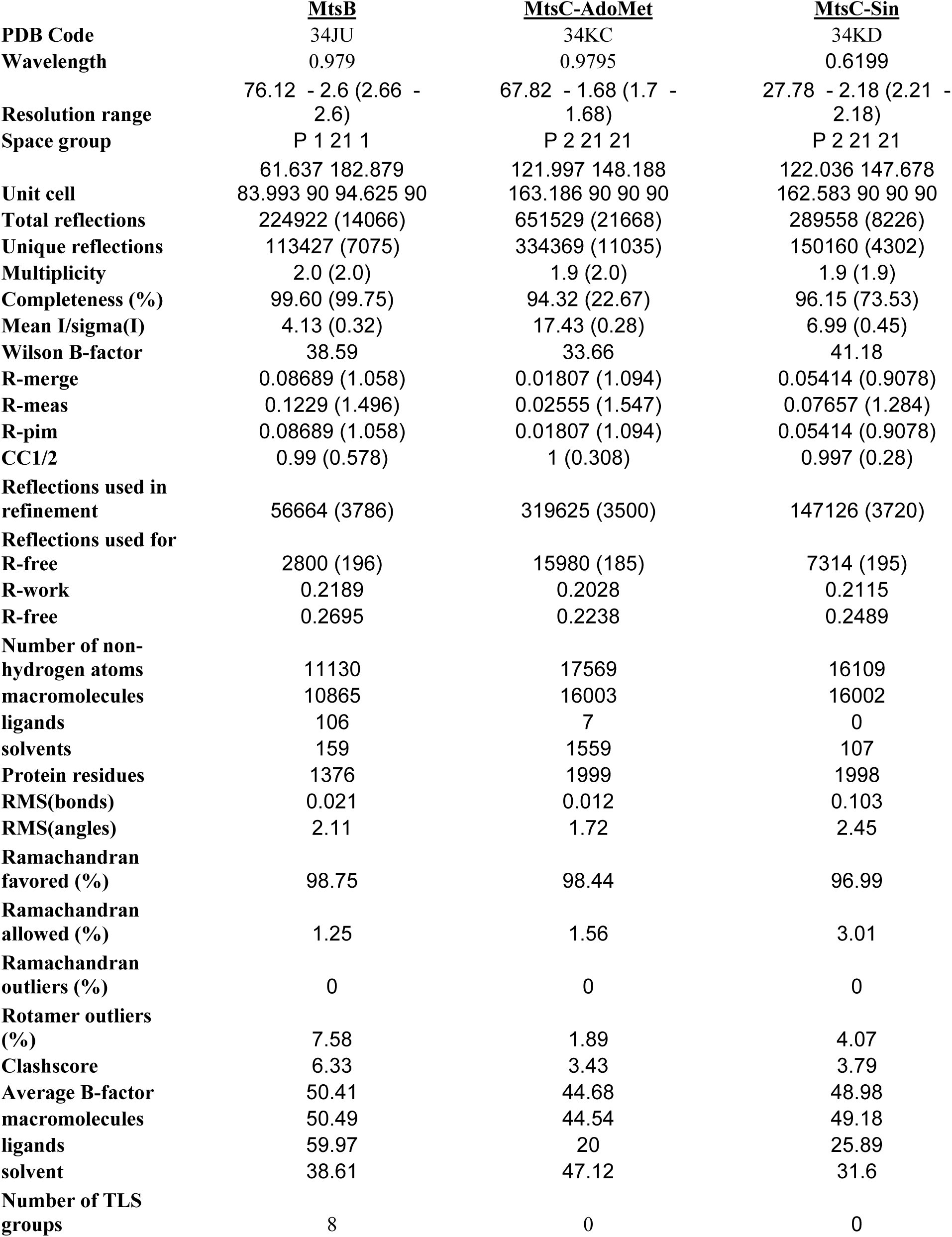
X-ray data collection and refinement statistics.

#### MtsC structure

The structure of monomeric MtsC bound to AdoMet was solved to 1.7 Å resolution by X-ray crystallography (**Fig. 4C**, **Table 2**). MtsC comprises a positively charged and highly conserved group α MTase core with a central target-recognition domain (TRD), and flexible, negatively charged N- and C-terminal domains (NTD and CTD) (**Fig. 4C; Supplementary Fig. 19**). This architecture is distinct from that of the group γ PglX MTase found in classical BREX^34,35^. The MTase core contains a bound AdoMet adjacent to Motif I (F_241_AGGG_245_), which mediates cofactor binding, and Motif IV (D_621_PPY_624_), which is involved in methyl-group transfer. A second structure of MtsC bound to the inhibitor sinefungin was solved to 2.2 Å resolution (**Table 2, Supplementary Fig. 19B**). The overall structures were highly similar (RMSD = 0.2 Å over 6882 atoms), with the ligands adopting near-identical binding conformations; the principal difference was the presence of additional water-mediated interactions stabilising sinefungin binding.

Having obtained individual structures, MtsA, MtsB, MtsC and the 46 bp dsDNA used in **Supplementary Fig. 14** were then combined to generate a 2.8 Å Cryo-EM structure of an MtsC-DNA complex (**Fig. 4D**, **Table 1**; **Supplementary Figs. 20–21)**. Although required for complex formation, density attributable to either MtsA or MtsB could not be identified, consistent with weak or transient interactions between MANTIS components.

MtsC binds DNA within the central positively charged groove of its MTase domain, primarily through a loop (aa 37–58) preceding Motif X that inserts into the minor groove (**Fig. 4D, inset**). Although the density in this region is insufficient to define individual side-chain positions, the loop contains several lysine and arginine residues that likely form non-specific contacts with the DNA phosphate backbone. The MTase domain and TRD are well resolved, whereas the CTD is absent from the cryo-EM density despite focussed refinements and variability analysis using focussed masks, indicating substantial flexibility. This contrasts with the well-defined CTD observed in the crystal structure. The apo and DNA-bound structures align closely (**Fig. 4E**; RMSD = 2.0 Å over 5,448 atoms), indicating that DNA binding induces little overall conformational change. It is therefore unclear from the structure why MtsA is needed for loading.

The density for bound DNA is weaker than that of MtsC, particularly adjacent to the DNA-recognition loop. Three-dimensional variability analysis revealed DNA bending towards this, but no evidence of target base-flipping (**Supplementary Fig. 21**, **Supplementary Movie 1**). Collectively, these observations suggest that the MtsC–DNA structure represents a primed, pre-catalytic DNA-bound intermediate rather than an active methylation complex. This interpretation is supported by the position of Motif VIII, which stabilises the flipped target adenine during methyl transfer. Motif VIII is located on a flexible loop at the junction of the MTase and CTD regions and is positioned away from the AdoMet-binding pocket in the observed structure. Given that ATP, MtsA, and MtsB are all required for methylation (**Fig. 3A**), we propose that additional interactions between MtsC and other MANTIS components promote conformational rearrangements that position Motif VIII for target-base capture and stabilise the DNA distortions required for methyl-group transfer.

### MtsD is structurally related to motor-driven nucleases

We have not yet succeeded in reconstituting MANTIS restriction activity *in vitro*, despite purifying MtsD to high homogeneity. For example, DNA isolated from sensitive and resistant phages was incubated with MtsA–D, ATP, AdoMet, and Mg²⁺, but no DNA cleavage was detected (**Supplementary Fig. 22**). Extensive variation of buffer conditions (including different divalent metals) and cofactors has likewise failed to produce activity. Analytical SEC also failed to detect interactions between MtsD and MtsA–C (**Supplementary Fig. 14D**).

To gain insight into MtsD function, we solved the structure of dimeric MtsD to 2.2 Å by Cryo-EM (**Fig. 5A**, **Table 1**; **Supplementary Figs. 23–25, Supplementary Movie 2**). The dimer adopts a pyramidal architecture in which the two functionally-distinct SF2 helicase-like ATPase domains of each protomer form the base, while the nuclease domains associate above to generate one PLD active site. Like the Type II restriction enzyme BfiI^36^, this single active site is predicted to cleave dsDNA through sequential strand cleavage. Although no nucleotide is bound, substantial cavities surround both ATPase sites (4178 Å³ and 4020 Å³ in protomers MtsDα and MtsDβ, respectively)^37^. A large channel also runs between the nuclease domains, with the catalytic H122 residues projecting into its lumen. The nuclease domains display weaker density than the ATPase domains, consistent with conformational flexibility. Accordingly, 3D variability analysis revealed opening and closing of the channel, potentially allowing engagement of a nucleic acid substrate (**Supplementary Movie 2**).

**Figure 5.**
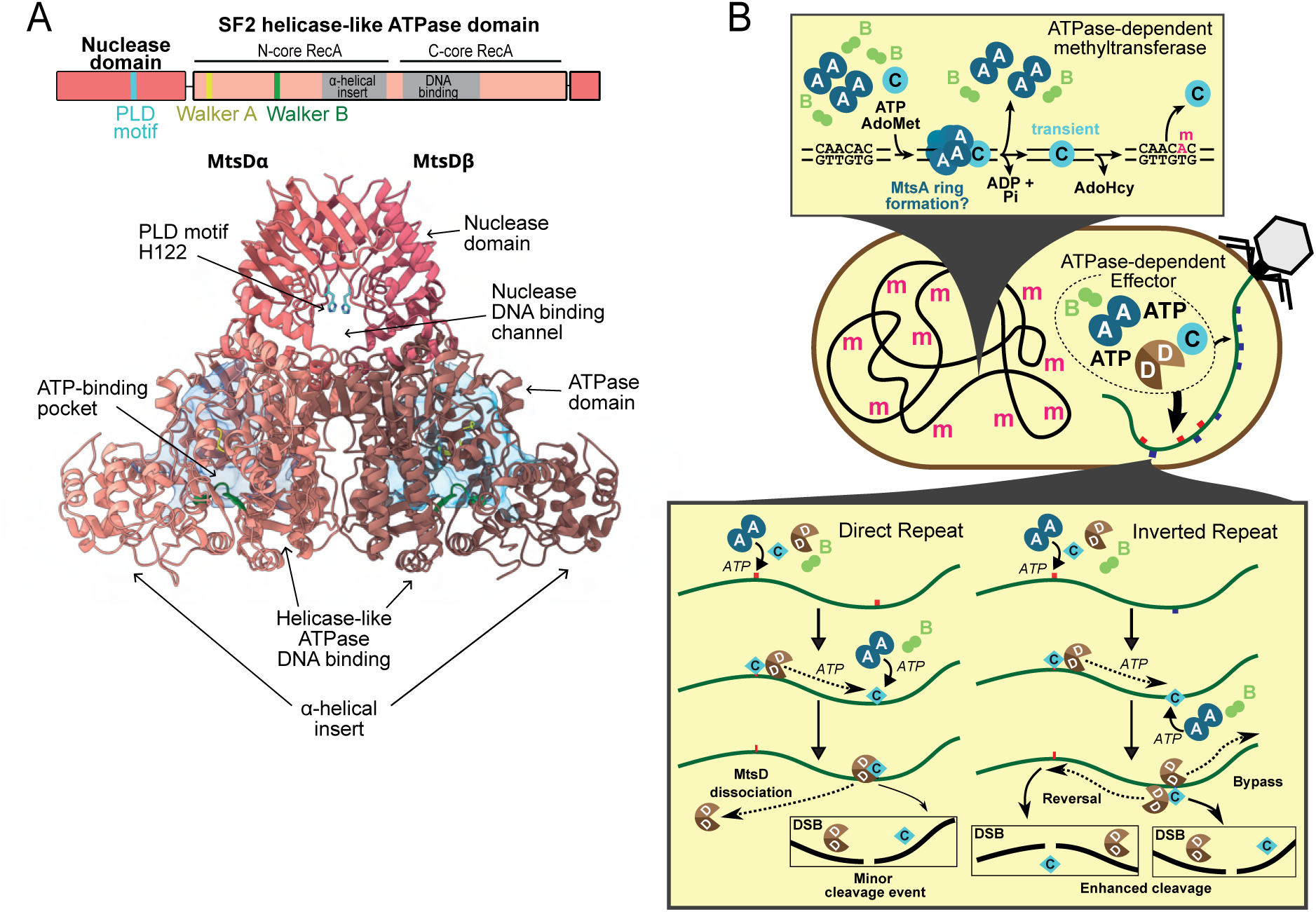
Structural analysis of MtsD and models for DNA methylation and cleavage by MANTIS. (**A**) Cryo-electron microscopy structure of the MtsD dimer at 2.2 Å resolution. The catalytic residue H122 is highlighted in stick form (cyan). The Walker A and Walker B motifs are shown in yellow and green, respectively. Transparent blue surfaces highlight cavities surrounding the ATP-binding pockets^37^. The domain map depicts the relative positions of domains and motifs. (**B**) Models for ATP-dependent DNA methylation at single sites and DNA cleavage dependent on long-range interactions between multiple sites. The central obround represents a *Pseudomonas* cell with a methylated (m) genome (black line) and a phage injecting its unmodified genome (green line) with MANTIS sites on the top (red) or bottom (blue) strands. Top inset: ATP-dependent hemimethylation by MtsABC. Bottom inset: ATP-dependent DNA cleavage by MtsABCD and a mechanism for site-orientation preference. See main text for full explanation.

MtsD shares structural similarity with the Type IV restriction enzyme SauUSI, which targets methylated DNA (PDB: 7CLG^38^; overall RMSD = 3.8 Å over 715 Cα atoms). The ATPase and nuclease folds are individually well conserved (RMSD = 3.6 Å over 714 Cα atoms and 5.1 Å over 413 Cα atoms, respectively). However, MtsD contains unique inserts within the helicase domain (**Supplementary Fig. 25**), whereas SauUSI possesses additional linker and target-recognition domains. These architectural differences likely reflect distinct target-selection mechanisms. SauUSI functions as an autonomous dimer, its TRDs recognising methylated cytosines and producing ATP-driven DNA cleavage. In contrast, MtsD is predicted to act downstream of DNA recognition by MtsABC and to restrict unmethylated targets. MANTIS therefore appears to have repurposed a SauUSI-like ATPase–PLD nuclease module for defence against unmethylated DNA. ATP-dependent control of DNA binding and/or nuclease activation may provide an additional safeguard against autorestriction, although the precise role of ATP hydrolysis in MtsD function remains unclear.

## DISCUSSION

Here, we characterise the MANTIS anti-phage and anti-plasmid defence system (previously Gao_RL or Gao_29)^6^, a novel member of a broad and widespread set of methylation-based defences such as RM, BREX and DISARM. Beyond shared protein architectures, MANTIS displays key BREX-like features, including hemimethylation of 6 bp sequences, MTase-directed specificity for both modification and restriction, and ATPase-dependent multi-protein assembly. However, unlike BREX^39^, susceptibility to MANTIS depends strongly on inverted recognition motifs. Although this site orientation preference resembles that of Type III and ISP RM systems^18–21^, MANTIS lacks their strict requirements: DNA with single or directly repeated sites can still be restricted, albeit less efficiently. While this graded response could imply weaker protection against autoimmunity, partial methylation does not impair host viability, as also observed in BREX^40^. This suggests differential handling of self versus non-self DNA, potentially aided by DNA packaging or other metabolic constraints^40^, or spatial separation of the defence machinery from the host chromosome^41,42^.

Consistent with evolutionary pressure applied by MANTIS, virulent phage genomes show a strand-biased depletion of inverted MANTIS sites that counters restriction. This evolutionary pattern distinguishes MANTIS from Type I and Type II RM, where selection favours site loss on both strands. Strand bias is also seen for Type III RM sites in both virulent and temperate phages. A less active restriction of direct repeats by MANTIS may be sufficient for temperate phage restriction so bias is not selective. Additionally, exceptions where phages evade MANTIS despite carrying inverted target pairs point to the existence of additional anti-MANTIS mechanisms, analogous to anti-RM, anti-BREX, or anti-DISARM^43^.

We propose a previously undescribed ATPase-activated DNA methylation that may have mechanistic parallels in other BREX systems (**Fig. 5**). MtsC cannot bind DNA alone but must be “loaded” by MtsA, reminiscent of roles for related ORC/Cdc6 clade AAA+ proteins in replication initiation and replisome clamp loading^44^. Nucleotide binding by MtsA and the presence of MtsB is required to trigger DNA methylation, leading to MtsC dissociation. ATP hydrolysis by MtsA enables multiple rounds of methylation. Recent structural data on the related AAA+ protein BrxC suggested a dimer and heptamer can interconvert and dynamic protein-protein interactions, which are also hinted at here. Further structural work will be required to resolve the conformational transitions underlying loading and methylation by MANTIS. The dimeric MtsB clearly contributes to methylation, but its precise mechanism is not evident from the isolated dimer structure. Although MtsB is unique to MANTIS, its vital role in activating methylation may be fulfilled by alternative proteins or domains in other BREX-related systems^8^.

We have yet to directly measure nuclease activity for MtsD but based on its structural similarity to the dimeric helicase–nuclease SauUSI^38^ (**Fig. 5**), we propose a working model that accounts for the graded dependence of restriction on site number and orientation (**Fig. 5**). For SauUSI, recognition of methylated cytosines (5mCs) activates directional ATP-dependent translocation along dsDNA using one helicase motor of the dimer^12^. Cleavage is triggered when 5mCs are encountered during translocation: 5mCs in direct repeat (same strand) support cleavage, but inverted repeats (opposite strands) either stimulate cleavage more strongly or activate the second helicase motor to engage DNA and translocate in the opposite direction. These translocation reversals generate a “ping-pong” motion between inverted sites, further promoting DNA cleavage.

By analogy, we propose that an MtsD dimer, loaded by MtsABC, tracks DNA and cleaves upon encountering another site-bound MtsABC (**Fig. 5**). Direct repeats produce “rear-end” collisions that yield dissociation or low levels of cleavage scaling linearly with site number. This accounts for occasional single-site plasmid digestion by motors travelling around the ring. In contrast, inverted repeats generate “head-on” collisions that support higher levels of cleavage or trigger reversals, leading to repeated site encounters and an exponential scaling of cleavage probability. Although restriction mechanisms in other BREX systems remain unclear and utilise different protein effectors, it would be informative to test if graded, site-orientation dependent restriction could be a broader feature; for example, by systematically varying BREX site number and orientation in plasmids, as here.

Many methylation-based defence systems, including MANTIS, BREX, MADS and some RM systems, encode ATPases despite DNA cleavage and methylation not being intrinsically ATP-dependent. We propose that ATP hydrolysis provides layered checkpoints that enhance defence specificity. In MANTIS, AAA+-dependent loading is required for MTase-mediated recognition of unmodified DNA, while a helicase-like ATPase activates nuclease activity and promotes ATP-driven communication between target sites. More broadly, multiple ATPases may improve self/non-self discrimination, reduce autoimmunity and favour defence retention. As ATP levels frequently decline during phage infection^45,46^, ATP-dependent modification and restriction may both be alleviated. This would allow viral propagation but would also reduce release of methylated escape progeny capable of infecting neighbouring cells with the same defence specificity. ATPase-based defences therefore may benefit both microbial individuals and communities.

**Limitations** – difficult to disentangle anti-MANTIS effects from site orientation preference; mechanism for MtsB activation of methylation unclear; order of DNA methylation and cleavage unclear as MtsABC needed for restriction and methylation at unmodified sites; structural interactions are transient, so mechanism needs more work; no activity of MtsD, so yet to confirm the SauUSI-like ping-pong mechanism; Role of MtsB unclear.

## METHODS & MATERIALS

### Plasmid construction, mutagenesis and tritium-labelling

DNA sequences are listed in **Supplementary Data File 2**. pLoLa^10^, an *Escherichia coli*/*Pseudomonas aeruginosa* shuttle vector designed according to the pSEVA vector architecture guidelines^47^, was used as the cloning backbone. The complete MANTIS defence system from strain AZPAE15024, including 240 bp upstream of the operon to preserve the native promoter region, was synthesised as DNA fragments (Twist Bioscience) and assembled into pLoLa using NEBuilder HiFi DNA Assembly Master Mix (New England Biolabs, NEB), generating pLoLa MANTIS. To generate the chimeric MANTIS construct (pLoLa u6684), a synthetic DNA fragment (Integrated DNA Technologies) containing the region encoding amino acids 431–544 of MtsC from PSE6684 and PCR-amplified overlapping sequences to the pLoLa MANTIS backbone were assembled using NEBuilder HiFi DNA Assembly Master Mix, replacing the equivalent region of the AZPAE15024 *mtsC* gene. The Type I RM system PaePAI from the *P. aeruginosa* PAO1 genome, including its native promoter region, was PCR-amplified from PAO1 genomic DNA and assembled into pLoLa using NEBuilder HiFi DNA Assembly Master Mix (NEB). A pLoLa plasmid carrying the *mCherry* gene in place of a defence system was used as a negative control in phage infection assays.

Targeted mutations were introduced into the MANTIS operon to generate individual gene knockouts. For *mtsA*, the coding sequence was deleted while retaining the final 24 bp to minimise disruption of potential regulatory elements controlling expression of downstream genes within the operon. Similarly, *mtsC* was deleted while retaining the final 42 bp. For *mtsD*, a premature stop codon was introduced by replacing the eighth codon (TCA) with TAA, resulting in truncation of the encoded protein. The mutations were generated by PCR using overlapping mutagenic primers. PCR products were resolved on a 1% (w/v) agarose gel, purified by gel extraction, and treated with DpnI (NEB) according to the manufacturer’s instructions, followed by heat inactivation. The resulting reaction mixture was transformed into chemically competent *E. coli* DH5α cells. The same approach was used to introduce a MANTIS site overlapping with an AleI site in the plasmid backbone. In this case the site sequences were added to overlapping regions of the PCR primers. Alanine point mutations were made using overlapping primers with Phusion polymerase (Thermo Fisher Scientific) using pLoLa AZPAE15024 with the overlapping MANTIS AleI site. This approach was also used with the methionine start codon mutants of *mtsR* and *mtsB* to introduce a premature stop codon; Δ*mtsB* required mutagenesis of two methionine residues (M1 and M12) to knock out protein activity. PCR products were treated with DpnI as described above and transformed into One shot Top10 chemically competent *E. coli* cells (Thermo Fisher Scientific).

To generate the pLoLa AZPAE15024 *mtsA* and *mtsC* knockout variants containing the overlapping MANTIS AleI site, the original Δ*mtsA* and Δ*mtsC* vectors were digested with HindIII-HF and NotI-HF (NEB). The 7041 bp and 7121 bp fragments corresponding to Δ*mtsA* and Δ*mtsC* respectively were band extracted from a 1% (w/v) agarose gel and purified using the Zymo Gel DNA Recovery kit via the manufacturer’s instructions. The same process was repeated for the pLoLA AZPAE15024 wt plasmid containing the overlapping site, with the 3789 bp backbone being recovered. Ligation was performed with T4 DNA Ligase (NEB) via the manufacturer’s instructions, and TOP10 cells were transformed prior to plasmid recovery.

pSJS1 was generated by site-directed mutagenesis of pUC19^48^ using overlapping primers and Phusion DNA polymerase (Thermo Fisher Scientific) to introduce a MANTIS site overlapping an AleI recognition sequence. The single-site transformation assay substrate (pLoLa SJS1) was generated by assembling the pLoLa backbone (10307 – 79 bp) with the pSJS1 target region (2652 - 790 bp) using NEBuilder HiFi DNA Assembly Master Mix (NEB). Additional substrates containing 0, 2, 4, or 10 target sites were generated by sequential modification of the target region. Direct-repeat and inverted-repeat variants containing two target sites were produced by site-directed mutagenesis using overlapping primers and Phusion DNA polymerase. Four-site variants were subsequently derived from the corresponding two-site constructs by further rounds of PCR mutagenesis. The zero-site control construct was generated from the single-site plasmid by mutagenesis to remove the target sequence. Construction of the three-site substrates was performed by insertion of annealed oligonucleotides into BamHI- and SbfI-digested two-site plasmids (pLoLa SJS2A and pLoLa SJS2B). Digested oligonucleotides and vector backbones were purified and ligated using T4 DNA ligase (NEB) according to the manufacturer’s instructions. Direct-repeat and inverted-repeat ten-site substrates were generated from the four-site constructs by NEBuilder HiFi DNA Assembly using a synthetic DNA fragment (IDT) containing six additional target sites.

Methylated 4-site direct and inverted plasmids were generated by incubation of 5 nM DNA with 300 nM MtsA, 300 nM MtsB and 50 nM MtsC in rCutSmart buffer (NEB) supplemented with 200 µM *S*-adenosyl methionine (AdoMet) and 2 mM ATP for 1 h at 37°C. Reactions were stopped at 65°C for 20 minutes before purification by addition of 1 volume of AMPure XP beads (Beckman) using the manufacturer’s protocol.

To produce expression constructs, synthetic genes encoding *mtsA-D*, codon optimised for expression in *E. coli* (Twist Bioscience for A, C and D, and Integrated DNA Technologies for B), were cloned into the pEHisV5TEV vector^49^ between the NcoI and BamHI sites and propagated using *E. coli* DH5α. Walker A (K80A) and Walker B (E174A) mutants of MtsA were made by Quikchange mutagenesis of the *mtsA* plasmid. Construct integrity was confirmed by sequencing (Eurofins Genomics, DE).

For tritium labelling, One Shot™ TOP10 Chemically Competent *E. coli* (Invitrogen) were transformed with pSJS1 and plated onto LB/agar supplemented with 50 µg/mL ampicillin. Single colonies were used to inoculate 2YT media and grown overnight at 37 °C. 500 mL of M9 media supplemented with 50 µg/mL ampicillin was inoculated with 4 mL of the 2YT overnight cultures. Cells were grown at 37 ^◦^C until OD600 was between 0.6-0.8 the cultures were supplemented with 170 µg/mL chloramphenicol and 0.5 mL 200 µM, 37 mBq/mL ^3^H-Thymidine (Perkin Elmer). Cultures were grown at 37 ^◦^C for a further 16-18 h. Cultures were pelleted at 6238*g* for 30 minutes at 4 °C before the plasmid was extracted using the Qiagen Plasmid Maxi Kit (Qiagen) following the manufacturer’s protocol except that the plasmid was eluted in 50 mM Tris (pH 8.0) and 50 mM EDTA. 1.0 g CsCl was added per mL of DNA and the DNA:CsCl solution added to an ultracentrifuge tube along with 0.5 mL liquid paraffin and 1 mg EtBr. After vigorous mixing, a gradient was generated using a VTi90 ultracentrifuge rotor at 55,000 rpm for 12 h at 15 °C. The supercoiled band was isolated and EtBr removed using repeated washes with CsCl-saturated isopropanol. Finally, supercoiled DNA was dialysed and stored in 10 mM Tris (pH 8.0) and 1 mM EDTA.

### Pseudomonas aeruginosa strains and phages

Strains and phages used are listed in **Supplementary Table 1**. The *Pseudomonas aeruginosa* PAO1-derived strains dPAO1 and viPAO1 were used throughout this study^22^.

For transformation, strains were grown overnight in LB broth at 37°C with shaking. 1 mL of culture was harvested by centrifugation, washed twice with 1 mM MgSO₄, and resuspended in 50 µL of 1 mM MgSO₄. 50 ng of plasmid DNA was added to the cell suspension, which was transferred to a 1 mm electroporation cuvette and electroporated at 1.8 kV. Immediately after electroporation, 1 mL of LB broth was added, and cells were allowed to recover for 1 h at 37°C with shaking before plating 100 µL onto LB agar supplemented with gentamicin (50 µg/mL).

For chromosomal integration of the MANTIS system into viPAO1, the complete locus was subcloned into a mini-Tn7 transposon carried on a pUC18-derived plasmid by restriction–ligation cloning using AscI and SpeI. Mini-Tn7 insertion was performed as previously described^50^ by co-transforming the mini-Tn7 construct with the helper plasmid pTNS2. For these transformations, 4 mL of overnight culture was used instead of 1 mL, and the entire recovery culture was plated onto selective LB agar.

The phage library used in this study comprised 163 phages, including 58 virulent and 105 temperate. Of these, 152 phages were genome-sequenced. The temperate phage collection was recently isolated in the Westra lab^22^, while the virulent phages include published examples such as DMS3^51,52^ or were sourced primarily from the Temperton lab (University of Exeter, UK), the Basel phage collection (ETH Zürich), the Brockhurst lab (University of Manchester), and the Lavigne lab (University of Leuven).

### Nanopore methylation sequencing

Transformation of *P. aeruginosa* with pLoLa MANTIS was carried out as above. Following transformation of One Shot™ TOP10 chemically competent *E. coli* (Invitrogen) with pLoLa MANTIS or pLoLa u6684, a single colony was used to inoculate a 5 mL overnight of LB supplemented with 5 µg/mL gentamicin. Saturated overnight cultures were harvested, and the plasmids were extracted using the QIAprep Spin Miniprep kit (Qiagen). Purified plasmids were linearised using Eco53kI using the manufacturer’s protocol (NEB) before purification with 1x volumes of AMPure XP beads (Beckman) using the manufacturer’s protocol.

Purified plasmid DNA was prepared and sequenced using an Oxford Nanopore Technologies (ONT) MinION with the Native Barcoding Kit 24 V14 using the manufacturer’s instructions. In brief, 1 µg purified plasmid DNA was end-repaired and dA-tailed using a combination of the NEBNext® Ultra™ II End Repair/dA-Tailing Module and NEBNext FFPE DNA Repair Mix (both NEB) by incubation for 5 minutes at 20 ^◦^C and 5 minutes at 65 ^◦^C before purification using AMPure XP beads (Beckman) using the manufacturer’s protocol. Each purified DNA was ligated to a Native Barcode using the Blunt/TA Ligase Master Mix (NEB) for 20 minutes at room temperature and pooled before purification using AMPure XP beads (Beckman). Finally, pooled barcoded DNA was ligated to the Native Adapter using the NEBNext Quick Ligation Module (NEB) for 20 minutes at room temperature before purification with AMPure XP beads, using the Short Fragment Buffer instead of ethanol.

Prepared DNA was loaded onto a R10.4.1 flowcell (ONT) and sequenced using MinKNOW for 2 hours. Raw pod5 files were basecalled and demultiplexed using Dorado (https://github.com/nanoporetech/dorado/, V0.5.0) with the “<u>dna_10.4.1_e8.2_400bps_sup@v4.2.0</u>” model with 6mA modification enabled. Demultiplexed data was aligned using Dorado, sorted and indexed using SAMtools ( https://github.com/samtools/samtools, V1.19) before a bed file was generated using Modkit (https://github.com/nanoporetech/modkit, V0.2.4). Low coverage events were filtered using Excel (Microsoft) before normalisation against a plasmid with the same backbone lacking the MANTIS system.

### DNA cleavage assay for testing *in vivo* plasmid methylation

One Shot™ TOP10 chemically competent *E. coli* (Invitrogen) was transformed with wt or mutant pLoLa MANTIS plasmids, and 3 colonies used to inoculate 5 mL LB supplemented with 5 mg/mL gentamicin. Following overnight incubation at 13 ^◦^C, plasmid DNA was extracted using the QIAprep Spin Miniprep kit (Qiagen). 300 ng of plasmid DNA was incubated at 37 ^◦^C for 1 hour with AleI and, where noted, NotI (both NEB). Reactions were quenched with 0.2 volumes of DNA:Protein loading dye (10 mM Tris (pH 8.0), 100 mM EDTA, 15% (v/v) Ficoll-400, 1% (w/v) SDS, 0.03% (w/v) Bromophenol blue and 0.03% (w/v) Xylene Cyanol FF). Samples were separated by agarose gel electrophoresis and imaged using a EpiChem3 Darkroom (UVP bioimaging) using 326 nm illumination. Band intensity was quantified using ImageJ v. 2.16.0 (National Institutes of Health).

### Efficiency of plating assays

Plaque assays were conducted on lawns of dPA01^22^ carrying the pLola plasmid expressing the relevant defence system or mCherry as a control. Cultures were inoculated from single colonies into LB supplemented with 50 µg/mL gentamicin and were incubated overnight at 37 °C and 180 rpm. Bacterial lawns were prepared by combining 300 µL of culture with 10 mL of 0.6% (w/v) LB agar supplemented with 50 µg/mL gentamicin. Phage stocks were serial diluted in M9 media and 3 µL of each dilution spotted onto each lawn. At least three technical replicates were conducted for each phage-defence system combination. Plates were incubated overnight at 37 °C and the following day plaques were counted. Efficiency of plating (EOP) was calculated by dividing the phage titre on the defence system lawn by the titre on the control lawn.

To verify if plaques on lawns of dPAO1 strains carrying MANTIS or type I RM system PaePAI represented escape phages, three plaques for each phage-defence combination were randomly picked and diluted in M9 media, and serial dilutions spotted onto lawns carrying the respective defence system or the defenceless control, and EOPs calculated.

For 3/20 phages, there was visible lysis at higher phage concentrations but no discernible plaques at lower concentrations. Here, a score of 0.5 plaques was recorded for the first dilution for which there was no visible lysis. For phage PA11P2, there was visible lysis at higher phage concentrations but no discernible plaques at lower concentrations on control and defence system-carrying lawns. This is consistent with a ‘Lysis from without’ phenotype^53^ that can occur when high levels of phage adsorption cause bacterial cell death, without phage replication. Due to this, phage PA11P2 was removed from the analysis. For phage P176 this phenotype was observed on MANTIS and PaePAI lawns, and for phage P276, only on MANTIS lawns. As both P176 and P276 produced countable distinct plaques on the defenceless control lawn, the lack of plaques on the defence-carrying lawns suggests a real defensive phenotype, and thus here we retained the data in our analyses and recorded 0.5 plaques for the first dilution for which there was no visible lysis.

### Liquid media infection assays

Phage infection assays were conducted in liquid media in 96-well plates with optical density measured over time. Specifically, LB supplemented with 50 µg/mL gentamicin was inoculated with a single colony of viPAO1 carrying pLoLa MANTIS or pLoLa mCherry, and cultures incubated overnight at 37 °C and 180 rpm. 50 µl of overnight culture was diluted into 5 mL of LB and subsequently incubated at 37 °C and 180 rpm for 3 hours. The culture was diluted to an OD600 of 0.03 using LB supplemented with 50 µg/mL gentamicin to a final volume of 15 mL. In a pilot experiment, this culture was serially diluted and 50 µL of dilutions plated onto LB agar and incubated at 37 °C overnight. Colonies were counted and colony forming units (CFU) per mL of the original cultures calculated. 180 µL of this culture was transferred to each of the inner 60 wells of a 96 well plate, along with 20 µL of phage stock diluted to ensure a multiplicity of infection (MOI) of 0.1, based on the predicted count of CFUs in the inoculating culture. Outer wells were filled with 100 µl of LB to ensure stable temperature across the plate. A CLARIOstar plate reader (BMG Labtech) was used to measure OD600 of each well every 30 minutes for 18 h. A minimum of three biological replicates of each phage-defence system combination were conducted.

From the OD600 measurements, per-well growth curves were fit and area under the curve (AUC) calculated using the *trapezoid* function of the *integrate* module of the python library *SciPy*^54^. Due to high noise at the start of the infection assay, and the tendency for phage-resistant surface mutants to evolve at later timepoints, only data from 3 to 10 h post-inoculation were used to calculate the AUC. For a given phage, resistance was calculated by subtracting the AUC for the mCherry control strain in the presence of phage from the AUC for the MANTIS carrying viPAO1 strain in the presence of phage, and then this value was divided by the AUC for the mCherry control strain in the absence of phage (**Fig. 2A**).

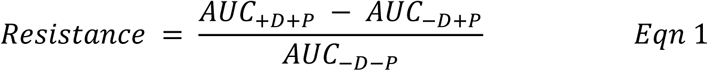

Similarly, phage strength was calculated as the reduction in AUC the phage caused in the mCherry carrying viPAO1 control strain.

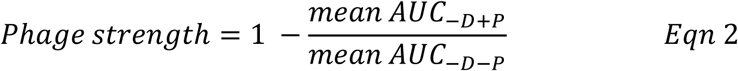

### Plasmid transformation assay

viPAO1 strains carrying either a chromosomal copy of the MANTIS defence system or no defence system (control) were transformed with plasmid DNA with varying numbers of MANTIS sites using the electroporation protocol described above. Following the 1 h recovery period, serial dilutions of the cultures were prepared in fresh LB, and 5 µL drops were plated onto both selective LB agar to enumerate transformants and non-selective LB agar to determine the total number of colony-forming units (CFU). Transformation efficiency was calculated as the ratio of transformants to total CFU. Relative transformation efficiency was calculated by normalising the transformation efficiency of the MANTIS-containing strain to that of the control viPAO1 strain transformed with the same plasmid.

### Evaluating depletion of *P. aeruginosa* MANTIS and RM system motifs from phage genomes

To investigate whether *P. aeruginosa* MANTIS (AZPAE15024 or Pae6684I) or RM system motifs (PacBio-validated and predicted recognition motifs for *P. aeruginosa* strains obtained from REBASE (**Supplementary Data File 1**) occur in phage genomes at rates different than expected by chance, the expected counts of sites per defence system per phage were calculated under a second-order Markov model, using observed genomic tri- and di-nucleotide frequencies. For a motif *M* = *m*_1_…*m*_L_ of length *L*, the probability *P* was estimated as:

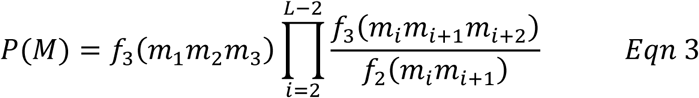

where *m*_i_ is the nucleotide at position i in the motif, and the quotient of *f*_3_(*xyz*) (the observed genomic frequency of trinucleotide *xyz*) and *f*_3_(*xy*) (the observed genomic frequency of dinucleotide *xyz*) estimates the conditional probability of nucleotide *z* given the preceding nucleotides *xy*. The product of Eqn 3 therefore represents the sequential multiplication of conditional probabilities under a second-order Markov chain. Expected counts of the motif in a phage genome are then given by multiplying *P*(*M*) by (*N* – *L* + 1). To get expected counts of degenerate motifs, any motif containing an IUPAC nucleotide code was expanded, and expected counts for all possible motifs were calculated and summed. Forward and reverse strand values were calculated separately.

Motif enrichment or depletion was quantified using a standardized residual (SR) approach. For each motif on each genome stand, the SR was calculated as:

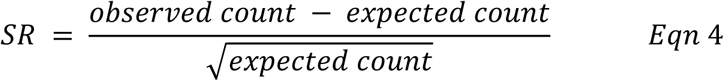

which approximates a z-score under a Poisson model. To incorporate the direction of deviation, a signed log-transformed SR was computed with:

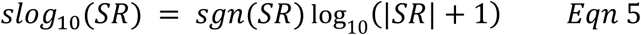

Motifs were classed as significantly depleted or enriched based on whether the slog_10_(SR) for a given strand exceeded the threshold of ±log_10_(2.576 + 1), corresponding approximately to a two-tailed 99% confidence interval on the underlying z-score. Motifs were classified as “Double enriched”. “Double depleted”. “Single enriched”, “Single depleted”, “Depleted and enriched”, or “Non-significant”, depending on the significance of depletion and/or enrichment on each strand.

### Expression and purification of MANTIS proteins

*E. coli* C43 (DE3) was transformed with an expression construct and protein expressed according to a previously described protocol^49^. For MtsA, -C and -D, 6 L of culture grown in LB medium were induced with 0.4 mM isopropyl-β-D-1-thiogalactoside (IPTG) at an OD600 of ∼0.8 and grown overnight at 16 °C with shaking. For Mts B, induction was carried out for 4 h at 37 °C. Cells were harvested (4000 rpm, 10 min, 4°C, Avanti JXN-26, Beckman Coulter, JLA-8.1 rotor) and resuspended in lysis buffer (50 mM Tris-HCl pH 7.5, 0.5 M NaCl, 10 mM imidazole,10% (v/v) glycerol, lysozyme, protease inhibitors (cOmplete EDTA-free protease inhibitor, Roche). Cells were lysed by sonicating for six times for 1 min on ice with 1 min rest intervals (Soniprep 150, MSE). Lysates were clarified by ultracentrifugation (40,000 rpm, 30 min, 4 °C Optima L-90K, 70Ti rotor) and filtered.

Proteins were purified using an immobilised metal affinity chromatography (IMAC) column (HisTrapFF crude, Cytiva), washed with 20 column volumes (CV) of buffer containing 50 mM Tris-HCl pH 7.5, 0.5 M NaCl, 30 mM imidazole and 10% (v/v) glycerol, followed by a step elution with buffer containing 50 mM Tris-HCl pH 7.5, 0.5 M NaCl, 0.5 M imidazole and 10% (v/v) glycerol on an NGC Chromatography System (BioRad). Protein containing fractions were concentrated and the affinity tag was removed by incubation of protein with Tobacco Etch Virus (TEV) protease (10:1) overnight at room temperature. Cleaved protein was isolated from TEV by repeating the IMAC step and collection of the unbound fraction. Size exclusion chromatography (Superdex 200 16/60 or 26/60, Cytiva) was used to further purify the proteins, with proteins eluted isocratically in a buffer containing 20 mM Tris-HCl pH 7.5, 250 mM NaCl. The proteins were concentrated using a centrifugal concentrator (Amicon Ultra-15, Merck) and stored frozen at -70°C.

For the His-tagged MtsA, -B or –C protein variants, *E. coli* BL21 Star (DE3) was transformed with the same expression vectors described above. Cultures in 1L LB medium were incubated at 37 °C until an OD600 of ∼0.8 before addition of 0.4 mM IPTG final followed by further incubation overnight at 16 °C with shaking. Cultures were harvested (5000 rpm, 30 min, 4°C, Avanti ??, Beckman Coulter, JLA-8.1 rotor) and resuspended in cell wash buffer (10 mM Tris-HCl pH 8.0, 100 mM NaCl, 1 mM EDTA) followed by a second centrifugation at 5000 rpm for 30 minutes at 4°C. Cells were resuspended in lysis buffer. Cells were lysed by sonication as described above but using a Sonics Vibra Cell ultrasonic homogenizer before clarification (21500 rpm, 30 min, 4°C, Avanti J-26 XP Centrifuge, Beckman Coulter, JA25.5 rotor). Following clarification, supernatant was dialysed using SnakeSkin™ 10kDa MWCO Dialysis Tubing into buffer containing 50 mM Tris-HCl pH 7.5, 0.5 M NaCl, 10 mM imidazole, 10% (v/v) glycerol and 1 mM p-mercaptoethanol (βME) for 2 hours at 4°C.

Dialysate was filtered and protein was purified using an IMAC column (HiTrap TALON crude, Cytiva), washed with 20CVs of buffer containing 50 mM Tris-HCl pH 7.5, 0.5 M NaCl, 20 mM imidazole and 10% (v/v) glycerol followed by a step elution with buffer containing 50 mM Tris-HCl pH 7.5, 0.5 M NaCl, 0.25 M imidazole and 10% (v/v) glycerol on a AKTA Prime system (Cytiva Life Science). Fractions containing protein were dialysed into buffer containing 50 mM Tris-HCl pH 7.5, 75 mM NaCl, 0.1 mM EDTA, 10% (v/v) glycerol and 1 mM βME before further purification on a Mono Q 5/50 GL (MtsA and –C) or Mono S 5/50 GL (MtsB). Columns were washed with 10 CVs of buffer containing 50 mM Tris-HCl pH 7.5, 125 mM NaCl, 0.1 mM EDTA, 10% (v/v) glycerol and 1 mM βME before elution with a NaCl gradient from 125-500 mM. The proteins were concentrated using a centrifugal concentrator (Amicon Ultra-15, Merck), adjusted to 50% (v/v) glycerol and stored at -20°C

### *In vitro* methylation assays

*In vitro* methylation assays were performed with 5 nM tritiated pSJS1 in rCutSmart buffer (NEB) with 200 µM AdoMet and 2 mM nucleotide cofactor (e.g. ATP, ATPvS, ADP, etc), as indicated. After addition of nucleotide, reactions were incubated for 30 min at 37 ° C before being stopped by heating at 65 ^◦^C for 20 minutes. DNA was treated with a combination of BsaI-HFv2 and AleI-v2 (NEB) for 30 min at 37 ^◦^C. Following digestions, samples were quenched with 0.2 volumes DNA:Protein loading dye, separated by agarose gel electrophoresis and imaged using a EpiChem3 Darkroom (UVP bioimaging) using 326 nm illumination. Bands were quantified using tritiated DNA quantification by scintillation counting.

Steady state conditions were then defined as 15 nM MtsA, 15 nM MtsB and 2.5 nM MtsC and single turnover conditions were defined as 300 nM MtsA, 300 nM MtsB and 50 nM MtsC. Reactions were quenched at the indicated time points by the addition of EDTA (pH 8.0) to a final concentration of 31.3 mM, before purification with 1 volume of AMPure XP beads using the manufacturer’s protocol. Methylation was validated by digestion of 120 ng DNA in rCutSmart buffer with AleI-v2 with or without BsaI-HFv2, with separation and quantification as described above. Nucleotide assays were performed as described above, with incubation of reactions with 2 mM nucleotides for 30 minutes at 37 ^◦^C.

### Electrophoretic Mobility Shift Assays

DNA oligonucleotides were labelled using T4 polynucleotide kinase (T4 PnK) and [v-32P] ATP (0.74 MBq) at 37 °C for 1 h in T4 PnK Buffer (NEB). Following incubation at 80°C for 10 min, labelled DNA was purified using Micro Bio-Spin chromatography columns (Bio-Rad Laboratories). Duplex DNA was annealed by mixing the ^32^P-labelled oligonucleotide with 1.05-fold of its unlabelled complementary strand in 10 mM Tris (pH 8.0), 1 mM EDTA and 50 mM NaCl before incubation for 5 minutes at 95 ^◦^C followed by slow cooling to ambient temperature.

For protein variation assays (**Fig. 3E**), 0.5 nM ^32^P-labelled duplex DNA was incubated in rCutSmart buffer (NEB) with 200 nM MtsA, MtsB and/or MtsC, plus 200 µM AdoMet/sinefungin/*S*-adenosylhomocysteine (AdoHcy) and/or 2 mM nucleotides, as indicated. For protein titration assays (Fig. 3F), 0.5 nM ^32^P-labelled duplex DNA was incubated in rCutSmart plus 200 µM AdoMet with either 50 nM MtsA, MtsAB, MtsC or MtsBC and varying concentrations of MtsC or MtsA, as indicated. Nucleotide displacement assays (Fig. 3C) were performed by incubation of 0.5 nM DNA with 50 nM MtsABC with 200 µM AdoMet and 2 mM nucleotide, as indicated.

In all cases, reactions were incubated at room temperature for 20 min before addition of 6x EMSA loading dye (rCutSmart supplemented with 48% (v/v) sucrose and 0.04% (w/v) bromophenol blue). Free and bound species were separated using a 5% (w/v) 29:1 polyacrylamide gel in TBM buffer (89 mM Tris, 89 mM boric acid and 2 mM MgCl_2_) supplemented with Rhinohide (Invitrogen) at 12.5 V/cm for 1 h at 8 °C. Gels were dried onto 3MM Chr blotting paper (Whatman) at 80 °C for 1 hr under vacuum. Gels were imaged using a Typhoon FLA 9500 device (Cytiva) and band intensities were quantified using ImageJ v. 2.16.0 (National Institutes of Health).

For Anti-His antibody supershift experiments, 0.5 nM ^32^P-labelled duplex DNA was incubated in rCutSmart buffer (NEB) with 100 nM MtsA, MtsB and/or MtsC +/-His-tag where specified, plus 200 µM AdoMet. After 10 min of incubation at room temperature, 1 µg Anti His-tag (Clontech) was added as indicated. Following a further 10 min incubation samples were separated by EMSA as described above.

### Phosphate Binding Protein ATPase Assay

7-Diethylamino-3-((((2-maleimidyl)ethyl)amino)carbonyl)coumarin-labelled Phosphate Binding Protein (MDCC-PBP) was cultured, purified and labelled as previously described^55^.

Experiments were performed in the SF-61 DX2 with single mixing at 37 °C with a 455GG filter to record MDCC emissions with the excitation wavelength set to 437 nm. MDCC-PBP was calibrated by titrating inorganic phosphate (Pi) against 6 µM PBP and recording the emissions for 15 s with 2048 points. MANTIS reactions were mixed as indicated with final concentrations of 100 nM 100 bp oligoduplex containing a single recognition motif, with 75 nM MtsA/B and 12.5 nM MtsC, as indicated, with the addition of 6 µM PBP, 200 µM AdoMet and 2 mM ATP in rCutSmart Buffer (NEB).

Average photomultiplier signal at varying Pi concentrations with PBP were used as calibration to determine a linear regression in GraphPad Prism V11.0.2. Pi contamination from ATP stock was removed, and samples were converted to Pi release in µM using the linear regression described.

### Analytical size exclusion chromatography

Purified proteins were first buffer exchanged into the required buffer conditions and concentrations using centrifugal concentrators (Cytiva). For individual MANTIS protein stoichiometries, proteins in A150 buffer (20 mM Tris pH 7.9, 150 mM NaCl) were concentrated and combined to 10 µM final concentration. For assaying complex formation and DNA binding, proteins in A40 buffer (20 mM Tris pH 7.9, 40 mM NaCl) were combined at equimolar ratios. Where indicated, protein mixtures also included combinations of ATP (4 mM), ATPoS (4 mM), AdoMet (1 mM), sinefungin (1 mM) and DNA oligonucleotides. For data shown in Fig. 4B, protein and DNA samples were at 5 µM and used a 46 bp version of the DNA oligonucleotide version of that used in biochemical assays (DNA_46bp_). Separation was performed on an Akta Pure (Cytiva) using a Superdex 200 increase 16/160 column at 0.25 ml/min.

### Mass photometry

Mass photometry experiments were performed on a TwoMP (Refeyn). The instrument was calibrated using Thyroglobulin, aldolase and conalbumin from a high molecular weight Gel Filtration Calibration Kit (Cytiva). Samples and calibrants were buffer exchanged into either A40 (20 mM Tris pH 7.9, 40 mM NaCl) or A150 (20 mM Tris pH 7.9, 150 mM NaCl) and diluted to 5 – 10 nM final concentration. For combinations of purified proteins, samples were mixed and incubated for 30 minutes at room temperature before measurement. Samples were measured on poly-lysine treated high-sensitivity microscope slides. One-minute videos were recorded on AcquireMP v2.5 software (Refeyn) and resulting data were analysed using DiscoverMP v2.5 software (Refeyn)

### Cryo-electron microscopy

Cryo-Electron Microscopy experiments were performed at the York Structural Biology Laboratory (York University), eBIC (Diamond Light Source) and the Astbury Centre for Structural Biology (Leeds University) for MtsA, MtsC-DNA and MtsD, respectively. Data collection and processing workflows are described below and shown in Supplementary Figs. 15, 20 and 23, respectively.

#### MtsA

MtsA was buffer exchanged into A150 ( 20 mM Tris pH 7.9, 150 mM NaCl), diluted to 1 mg/ml. and 3 μL added to a plasma-cleaned and carbon film-coated 2/2 holey carbon 200 nm mesh copper grid. Grids were blotted for 5s and plunge-frozen in liquid ethane (Vitrobot; Thermofisher). Grids were loaded onto a Glacios 200 kV electron microscope (Thermofisher), a dataset of 1677 movies collected at a resolution of 0.56 Å, and data transferred to cryoSPARC^56^ for processing. Micrographs were motion and CTF corrected, and then manually curated to 1399 micrographs before being denoised using a training set of 100 micrographs. An automated gaussian search and curation produced ∼78,000 initial particles, which were extracted at 700 pixels with Fourier cropping to 256 pixels. Iterative rounds of 2D classification and selection produced 9000 particles across 10 classes which were used for template picking. Manual particle curation and extraction produced ∼220,000 particles from the complete dataset which were extracted at 1200 pixels and Fourier cropped to 256 pixels. Iterative rounds of 2D classification and selection produced a final set of 38,488 particles, which were fed into an initial *ab initio* reconstruction. This particle set and volume were then subjected to iterative rounds of heterologous refinement, with one correct starting volume and one junk starting volume, to clean up the particle stack and reduce noise. This resulted in a final set of 22,717 particles, which were non-uniform refined to 4 Å. An AlphaFold3^57^ model of the MtsA1-480 dimer was fitted to the density map in ChimeraX^58^ and iterative rounds of building and refinement were carried out using Coot^59^ and Phenix^60^, respectively. DeepEMhancer^61^ was used for model sharpening and map visualisation. Ligand density Polder Omit maps were produced with Phenix.

#### MtsC-DNA

MtsC was mixed with purified MtsA_E174A_ and DNA_Mts_46_ in A40 (20 mM Tris pH 7.9, 40 mM NaCl) buffer at equimolar (5 µM) concentration in the presence of 1 mM ATP and 4 mM AdoMet and incubated at 37°C for 30 minutes. Three microlitres of the mixture was then added to a plasma-cleaned and carbon film-coated 2/2 holey carbon 200 nm mesh copper grid, for 5s and plunge-frozen in liquid ethane (Vitrobot, Thermofisher). Grids were loaded onto a Tundra 300 kV electron microscope (Thermofisher), a dataset of 10,022 movies was collected at a resolution of 0.653 Å and data were transferred to cryoSPARC for processing. Following motion and CTF correction, initial processing of and curation utilised 161 micrographs for gaussian particle fitting, producing 22 2D classes (∼18,000 particles) which were used for template picking on the complete dataset. Particle curation resulted in ∼2.3M particles extracted at 400 pixels. Iterative rounds of 2D classification and class building produced 31 classes (696,244 particles) which were used for *ab initio* reconstruction and refined to 2.66 Å. To resolve the region of the map corresponding to DNA, 3D classification was run using a focussed mask to separate particles with consistent DNA density. Resulting maps were inspected manually in ChimeraX and particle classes with homogenous DNA density were binned and non-uniform refined together to a resolution of 2.8 Å. The 2.66 Å map and the 2.8 Å map were then combined in Phenix to merge optimum density for MtsC and DNA regions, respectively. The MtsC crystal structure and a manually-built DNA oligonucleotide were then manually fitted to the density in ChimeraX before iterative rounds of building and refinement in Coot and Phenix, respectively. DeepEMhancer and Phenix were used for model sharpening and map visualisation.

#### MtsD

MtsD was diluted to 5 mg/ml in A150 (20 mM Tris pH 7.9, 150 mM NaCl). Three microlitres of MtsD was added to a plasma-cleaned and carbon film-coated 1.2/1.3 holey carbon 200 nm mesh copper grid. Grids were blotted for 8s and plunge-frozen in liquid ethane (Vitrobot, Thermofisher). Grids were loaded onto a Tundra 300 kV electron microscope (Thermofisher) and a dataset of 8400 movies was collected at a resolution of 0.85 Å and data were transferred to cryoSPARC for processing. Initial processing was performed on 1657 micrographs to produce 2D class templates for template picking on the complete dataset. Particle picking and curation yielded ∼2.5M particles which were extracted at a 400 pixel box size. Iterative rounds of 2D classification and class selection produced 41 2D classes comprising 992,729 particles. *Ab initio* reconstruction and refinement with C2 symmetry generated a 2.12 Å density map. An AlphaFold3 model of the MtsD dimer was fitted to the density map in ChimeraX and iterative rounds of building and refinement were carried out using Coot and Phenix, respectively. To model the flexible nuclease domain, ModelAngelo^62^ was used to trace the peptide backbone through the density and residues were built manually in Coot. DeepEMhancer was used for model sharpening and map visualisation.

### X-ray crystallography

MtsB and MtsC were concentrated to 12 mg/ml in crystal buffer (20 mM Tris pH 7.9, 150 mM NaCl, 2.5 mM DTT) for crystal screening. Crystal screens were set using the sitting drop vapour diffusion method using a Mosquito Xtal3 liquid handling robot (SPT Labtech) and crystal screens were incubated at 18 °C. All commercially available crystal screens were produced by Molecular Dimensions. MtsB initially crystallised in JCSG+ well E2, though crystals diffracted poorly. Further optimisation produced crystals in 50 mM Tris pH 7.75, 1.125 M LiSO4. MtsC was incubated with 1 mM AdoMet or sinefungin for 30 minutes on ice before crystallisation and crystals were observed in Morpheus I well G10 (0.1 M carboxylic acids, 0.1 M buffer system 3 pH 8.5, 30% v/v precipitant mix 2). Crystallisation was confirmed by microscopy, and crystals extracted using nylon loops and flash frozen in liquid nitrogen. Data collection was carried out remotely at Diamond Light Source, Oxford, UK on beamline I04.

Initial data processing was carried out with automated IspyB processes using Xia2-DIALS for space group determination and dataset merging^63^. Further data reduction was carried out using AIMLESS in CCP4i2^64^. Both MtsB and MtsC structure solutions were produced by molecular replacement with AlphaFold3 models using Phaser^65^. Iterative rounds of building and refinement were then carried out using Coot and Refmac, respectively. For MtsB, the asymmetric crystal unit contained 8 copies of MtsB, each forming dimer pairs with MtsB molecules in adjacent asymmetric units. Structure validation was performed with CCP4i2, Phenix, Coot and the PDB validation server. Ligand density Polder Omit maps were produced with Phenix. Visualisation and figure creation was performed in ChimeraX.

### Statistical analysis and data fitting

Unless stated otherwise, a minimum of three technical repeats were carried out for experiments and mean values and errors are indicated in the relevant legends.

All statistical analyses relating to the infection assays and efficiency of plating experiments were conducted in R v. 4.3.1 (available from https://www.R-project.org/) and utilized Generalized Linear Models (GLMs), that were built using the base R function *glm*. Initially global GLMs were constructed, and significance testing was performed via model simplification.

Terms were retained in the minimal adequate model if their removal significantly reduced model fit (p < 0.05). Model assumptions were checked using the *DHARMa* package (available from https://cran.r-project.org/web/packages/DHARMa/index.html), which uses simulation-based residual diagnostics. Overdispersion, model fit (using a Kolmogorov-Smirnov test), influential outliers, and heteroscedasticity of residuals, were all tested for. For all post-hoc analyses the R package *emmeans* was utilised (available from https://github.com/rvlenth/emmeans), including the test of whether resistance conferred by a defense system in the liquid media infection assays is significantly different to 0, and if EOP is significantly different to 1.

The *tidyverse* packages^66^ were used for data handling. To create figures the R packages *ggplot2*^66^, *ggpubr* (available from https://cran.r-project.org/web/packages/ggpubr/index.html), *cowplot* (available from https://cran.r-project.org/web/packages/cowplot/index.html), *RColorBrewer* (available from https://cran.r-project.org/web/packages/RColorBrewer/index.html), *gridExtra* (available from https://cran.r-project.org/web/packages/gridExtra/index.html), and *patchwork* (available from https://cran.r-project.org/web/packages/patchwork/index.html) were used. Kinetic and thermodynamic data were analysed using GraphPad Prism version 11.0.2 for Windows (GraphPad Software, Boston, Massachusetts USA)

## Acknowledgements

We thank Ben Temperton and Christian Fitch (University of Exeter), Mike Brockhurst and Rosanna Wright (University of Manchester), Alexander Hams (ETH Zürich), and Rob Lavigne (University of Leuven) for providing virulent phages used in this study. We thank Durham University and the Centre for Programmable Matter for the use of mass photometry (Refeyn TwoMP) equipment. We gratefully acknowledge Diamond Light Source for time on beamlines I04 and I24 under proposal MX32736. We thank the York Structural Biology Laboratory (York University), eBIC (Diamond Light Source) and the Astbury Centre for Structural Biology (Leeds University) for access to their respective cryo-EM facilities.

## Author Contributions

**Shaw** – Investigation, methodology, formal analysis, data curation, visualisation, writing – review and editing

**Went** – Investigation, methodology, formal analysis, data curation, visualisation, writing original draft, writing – review and editing

**Agapov** – Investigation, methodology, formal analysis, data curation, visualisation, writing – review and editing

**Westley** – Investigation, methodology, formal analysis, data curation, visualisation, writing original draft, writing – review and editing

**Olina** – Investigation, methodology, writing – review and editing

**Graham** – Investigation, methodology, writing – review and editing

**Chong** – Investigation, methodology, formal analysis, data curation, visualisation, writing original draft, writing – review and editing

**MultiDefence Consortium** - funding Acquisition, project administration, methodology

**Baker** - Supervision, funding acquisition, writing – review and editing

**White** – Conceptualization, supervision, funding acquisition, writing – review and editing

**Van Houte** - Supervision, funding acquisition, resource provision, writing – review and editing

**Westra** - Supervision, funding acquisition, resource provision, writing – review and editing

**Blower** - Conceptualization, formal analysis, visualisation, supervision, funding acquisition, writing original draft, writing – review and editing

**Szczelkun** - Conceptualization, formal analysis, visualisation, supervision, funding acquisition, writing original draft, writing – review and editing

## REFERENCES

1. Tesson, F. et al. Systematic and quantitative view of the antiviral arsenal of prokaryotes. Nat. Commun. 13, 2561 (2022).

2. Mordret, E. et al. Protein and genomic language models uncover the unexplored diversity of bacterial immunity. Science (1979). 392, (2026).

3. Néron, B., et al. MacSyFinder v2: Improved modelling and search engine to identify molecular systems in genomes. Peer Community Journal 3, (2023).

4. Tesson, F. et al. A Comprehensive Resource for Exploring Antiphage Defense: DefenseFinder Webservice,Wiki and Databases. Peer Community Journal 4, 000–000 (2024).

5. Payne, L. J. et al. PADLOC: a web server for the identification of antiviral defence systems in microbial genomes. Nucleic Acids Res. 50, W541–W550 (2022).

6. Gao, L. et al. Diverse enzymatic activities mediate antiviral immunity in prokaryotes. Science 369, 1077–1084 (2020).

7. Goldfarb, T. et al. BREX is a novel phage resistance system widespread in microbial genomes. EMBO J. 34, 169–83 (2015).

8. Rakesh, S. & Krishnan, A. Expanding the landscape of BREX diversity: uncovering multi-layered functional frameworks and identification of novel BREX-related defense systems. Nucleic Acids Res. 54, (2026).

9. Tock, M. R. & Dryden, D. T. The biology of restriction and anti-restriction. Curr. Opin. Microbiol. 8, 466–472 (2005).

10. Maestri, A. et al. The bacterial defense system MADS interacts with CRISPR-Cas to limit phage infection and escape. Cell Host Microbe 32, 1412–1426.e11 (2024).

11. Ofir, G. et al. DISARM is a widespread bacterial defence system with broad anti-phage activities. Nat. Microbiol. 3, 90–98 (2018).

13. Shaw, S. J. et al. Individual Staphylococcus aureus SauUSI restriction endonuclease motors fragment methylated DNA. bioRxiv 2025.10.16.682607 (2025) doi:10.1101/2025.10.16.682607.

13. Toliusis, P., Zaremba, M., Silanskas, A., Szczelkun, M. D. & Siksnys, V. CgII cleaves DNA using a mechanism distinct from other ATP-dependent restriction endonucleases. Nucleic Acids Res. 45, 8435–8447 (2017).

14. Toliusis, P. et al. The H-subunit of the restriction endonuclease CglI contains a prototype DEAD-Z1 helicase-like motor. Nucleic Acids Res. 46, 2560–2572 (2018).

15. Kaiser, A. J. et al. Multiple forms of protein–protein and DNA binding are exhibited by BrxC from the BREX phage restriction system. Nucleic Acids Res. 54, (2026).

16. Chong, C. E. et al. Defence systems drive accessory genome interactions in Pseudomonas aeruginosa. ISME Communications 6, 130 (2026).

17. Roberts, R. J., Vincze, T., Posfai, J. & Macelis, D. REBASE: a database for DNA restriction and modification: enzymes, genes and genomes. Nucleic Acids Res. 51, D629–D630 (2023).

18. Meisel, A., Bickle, T. A., Kriiger, D. H. & Schroeder, C. Type III restriction enzymes need two inversely oriented recognition sites for DNA cleavage. Nature 1992 355:6359 355, 467–469 (1992).

19. Meisel, A., Mackeldanz, P., Bickle, T. A., Krüger, D. H. & Schroeder, C. Type III restriction endonucleases translocate DNA in a reaction driven by recognition site-specific ATP hydrolysis. EMBO Journal 14, 2958–2966 (1995).

20. Smith, R. M., Diffin, F. M., Savery, N. J., Josephsen, J. & Szczelkun, M. D. DNA cleavage and methylation specificity of the single polypeptide restriction–modification enzyme LlaGI. Nucleic Acids Res. 37, 7206–7218 (2009).

21. Van Aelst, K. et al. Type III restriction enzymes cleave DNA by long-range interaction between sites in both head-to-head and tail-to-tail inverted repeat. Proc. Natl. Acad. Sci. U. S. A. 107, 9123–9128 (2010).

22. Olina, A. et al. Engineering the Pseudomonas aeruginosa virus-isolation host viPAO1 reveals how host genetic barriers shape recoverable phage diversity. bioRxiv 2026.09.16.752016 (2026) doi:10.64898/2026.09.16.752016.

23. Doberenz, S. et al. Identification of a Pseudomonas aeruginosa PAO1 DNA Methyltransferase, Its Targets, and Physiological Roles. mBio 8, e02312–16 (2017).

24. Rusinov, I. S., Ershova, A. S., Karyagina, A. S., Spirin, S. A. & Alexeevski, A. V. Avoidance of recognition sites of restriction-modification systems is a widespread but not universal anti-restriction strategy of prokaryotic viruses. BMC Genomics 2018 19:1 19, 885-(2018).

25. Shaw, L. P., Rocha, E. P. C. & Maclean, R. C. Restriction-modification systems have shaped the evolution and distribution of plasmids across bacteria. Nucleic Acids Res. 51, 6806–6818 (2023).

26. Shaw, L. P. et al. The leading region of many plasmids is adapted for translational efficiency. Nucleic Acids Res. 54, 13–14 (2026).

27. Krüger, D. H., Kupper, D., Meisel, A., Reuter, M. & Schroeder, C. The significance of distance and orientation of restriction endonuclease recognition sites in viral DNA genomes. FEMS Microbiol. Rev. 17, 177–184 (1995).

28. Schroeder, C., Jurkschat, H., Meisel, A., Reich, J. G. & Krüger, D. Unusual occurrence of EcoP1 and EcoP15 recognition sites and counterselection of type II methylation and restriction sequences in bacteriophage T7 DNA. Gene 45, 77–86 (1986).

29. Sears, A., Peakman, L. J., Wilson, G. G. & Szczelkun, M. D. Characterization of the Type III restriction endonuclease PstII from Providencia stuartii. Nucleic Acids Res. 33, 4775–4787 (2005).

30. Krüger, D. H. & Bickle, T. A. Bacteriophage survival: multiple mechanisms for avoiding the deoxyribonucleic acid restriction systems of their hosts. Microbiol. Rev. 47, 345–360 (1983).

31. Lindsey, J. C. & Leach, D. R. F. Slow replication of palindrome-containing DNA. J. Mol. Biol. 206, 779–782 (1989).

32. Carver, A., Zhang, B. & Zhang, X. Structures and mechanisms of AAA+ protein complexes in DNA processing. Curr. Opin. Struct. Biol. 92, 103056 (2025).

33. Holm, L., Laiho, A., Törönen, P. & Salgado, M. DALI shines a light on remote homologs: One hundred discoveries. Protein Sci. 32, (2023).

34. Went, S. C. et al. Structure and rational engineering of the PglX methyltransferase and specificity factor for BREX phage defence. Nat. Commun. 15, (2024).

35. Drobiazko, A. et al. Molecular basis of foreign DNA recognition by BREX anti-phage immunity system. Nat. Commun. 16, 1825 (2025).

36. Sasnauskas, G., Halford, S. E. & Siksnys, V. How the BfiI restriction enzyme uses one active site to cut two DNA strands. Proc. Natl. Acad. Sci. U. S. A. 100, 6410–6415 (2003).

37. Guerra, J. V. da S., et al. pyKVFinder: an efficient and integrable Python package for biomolecular cavity detection and characterization in data science. BMC Bioinformatics 2021 22:*1* 22, 607-(2021).

38. Tumuluri, V. S., Rajgor, V., Xu, S. Y., Chouhan, O. P. & Saikrishnan, K. Mechanism of DNA cleavage by the endonuclease SauUSI: a major barrier to horizontal gene transfer and antibiotic resistance in Staphylococcus aureus. Nucleic Acids Res. 49, 2161–2178 (2021).

39. Kelly, A. et al. Diverse Durham collection phages demonstrate complex BREX defense responses. Appl. Environ. Microbiol. 89, (2023).

40. Isaev, A. et al. Phage T7 DNA mimic protein Ocr is a potent inhibitor of BREX defence. Nucleic Acids Res. 48, 5397–5406 (2020).

41. Zhang, Z. et al. Kiwa is a membrane-embedded defense supercomplex activated at phage attachment sites. Cell 188, 5862–5877.e23 (2025).

42. Sanchez-Londono, M. et al. Visualization of Type IV-A1 CRISPR-mediated repression of gene expression and plasmid replication. Nucleic Acids Res. 52, 12592–12603 (2024).

43. Niault, T., van Houte, S., Westra, E. & Swarts, D. C. Evolution and ecology of anti-defence systems in phages and plasmids. Current Biology 35, R32–R44 (2025).

44. Bleichert, F., Botchan, M. R. & Berger, J. M. Crystal structure of the eukaryotic origin recognition complex. Nature 519, 321–326 (2015).

45. Li, J. et al. Structures and activation mechanism of the Gabija anti-phage system. Nature 629, 467–473 (2024).

46. Gaskell-Mew, L. et al. Inhibition by ATP regulates the activity of a CBASS anti-phage nucleotide cyclase. Biochem. J. 483, 1617–1630 (2026).

47. Martínez-García, E. et al. SEVA 4.0: an update of the Standard European Vector Architecture database for advanced analysis and programming of bacterial phenotypes. Nucleic Acids Res. 51, D1558–D1567 (2023).

48. Norrander, J., Kempe, T. & Messing, J. Construction of improved M13 vectors using oligodeoxynucleotide-directed mutagenesis. Gene 26, 101–106 (1983).

49. Rouillon, C., Athukoralage, J. S., Graham, S., Grüschow, S. & White, M. F. Investigation of the cyclic oligoadenylate signaling pathway of type III CRISPR systems. Methods Enzymol. 616, 191–218 (2019).

50. Choi, K. H. & Schweizer, H. P. mini-Tn7 insertion in bacteria with single attTn7 sites: example Pseudomonas aeruginosa. Nature Protocols 2006 1:1 1, 153–161 (2006).

51. Budzik, J. M., Rosche, W. A., Rietsch, A. & O’Toole, G. A. Isolation and Characterization of a Generalized Transducing Phage for Pseudomonas aeruginosa Strains PAO1 and PA14. J. Bacteriol. 186, 3270–3273 (2004).

52. Cady, K. C., Bondy-Denomy, J., Heussler, G. E., Davidson, A. R. & O’Toole, G. A. The CRISPR/Cas adaptive immune system of Pseudomonas aeruginosa mediates resistance to naturally occurring and engineered phages. J. Bacteriol. 194, 5728–5738 (2012).

53. Abedon, S. T. Lysis from without. Bacteriophage 1, 46–49 (2011).

54. Virtanen, P. et al. SciPy 1.0: fundamental algorithms for scientific computing in Python. Nature Methods 2020 17:3 17, 261–272 (2020).

55. Hirshberg, M. et al. Crystal Structure of Phosphate Binding Protein Labeled with a Coumarin Fluorophore, a Probe for Inorganic Phosphate,. Biochemistry 37, 10381–10385 (1998).

56. Punjani, A., Rubinstein, J. L., Fleet, D. J. & Brubaker, M. A. cryoSPARC: algorithms for rapid unsupervised cryo-EM structure determination. Nature Methods 2017 14:3 14, 290–296 (2017).

57. Abramson, J. et al. Accurate structure prediction of biomolecular interactions with AlphaFold 3. Nature 2024 630:8016 630, 493–500 (2024).

58. Goddard, T. D. et al. UCSF ChimeraX: Meeting modern challenges in visualization and analysis. Protein Science 27, 14–25 (2018).

59. Emsley, P. & Cowtan, K. Coot: model-building tools for molecular graphics. Acta Crystallogr D Biol Crystallogr 60, 2126–2132 (2004).

60. Adams, P. D. et al. PHENIX: a comprehensive Python-based system for macromolecular structure solution. Acta Crystallogr. D Biol. Crystallogr. 66, 213–21 (2010).

61. Sanchez-Garcia, R. et al. DeepEMhancer: a deep learning solution for cryo-EM volume post-processing. Communications Biology 2021 4:1 4, 874- (2021).

62. Jamali, K. et al. Automated model building and protein identification in cryo-EM maps. Nature 2024 628:8007 628, 450–457 (2024).

63. Winter, G. *xia2* : an expert system for macromolecular crystallography data reduction. J. Appl. Crystallogr. 43, 186–190 (2010).

64. Winn, M. D. et al. Overview of the CCP4 suite and current developments. Acta Crystallogr. D Biol. Crystallogr. 67, 235–42 (2011).

65. McCoy, A. J. et al. Phaser crystallographic software. J. Appl. Crystallogr. 40, 658–674 (2007).

66. Wickham, H. et al. Welcome to the Tidyverse. J. Open Source Softw. 4, 1686 (2019).

