## Supplementary Table 1 for "Dual ATPase-activated DNA hemimethylation and cleavage by the MANTIS defence system"

| Type | Name | Description | Use in this study | Source |
|---|---|---|---|---|
| Bacteria | *P. aeruginosa* dPAO1 | PAO1 with Δ*gabija*, Δ*retron1B*, Δhelduf2290, ΔPaepaI, ΔPf4, ΔCtx | Strain lacking defences for analysing MANTIS activity: Figs. 1 and 2 | [22] |
| Bacteria | *P. aeruginosa* viPAO1 | dPAO1 with ΔPf6 | Strain lacking defences and prophage for analysing MANTIS activity: Fig. 2 | [22] |
| Bacteria | *P. aeruginosa* viPAO1 mCherry | ViPAO1 with mCherry gene in miniTn7 insertion site | Transformation efficiency assay: Fig. 2I | This study |
| Bacteria | *P. aeruginosa* viPAO1 mCherry | ViPAO1 with AZPAE15024 MANTIS operon in miniTn7 insertion site | Transformation efficiency assay: Fig. 2I | This study |
| Bacteria | *E. coli* C43 (DE3) | F – ompT hsdSB (rB- mB-) gal dcm (DE3) | Protein expression | Sigma-Aldrich |
| Bacteria | *E. coli* DH5α | fhuA2::IS2 Δ(mmuP-mhpD)169 ΔphoA8 glnX44 ϕ80d[ΔlacZ58(M15)] rfbD1 gyrA96 luxS11 recA1 endA1 rphWT thiE1 hsdR17 | Plasmid propagation | New England Biolabs |
| Bacteria | *E. coli* Top10 | F–mcrA Δ(mrr-hsdRMS-mcrBC) φ80lacZΔM15 ΔlacX74 recA1 araD139 Δ(ara-leu)7697 galU galK λ–rpsL(StrR) endA1 nupG | Plasmid propagation and MANTIS methylation assays | Invitrogen |
| Phage | B3F9_PAnicatthedisco | Temperate, Abbreviation B3F9 | Fig. 2A, Fig. 2J | [22] |
| Phage | K7B12h_PAritosh | Temperate, Abbreviation K7B12h | Fig. 2A, Fig. 2J | [22] |
| Phage | B2A9_PAw | Temperate, Abbreviation B2A9 | Fig. 2A, Fig. 2J | [22] |
| Phage | D8_PAmela | Temperate, Abbreviation D8 | Fig. 2A, Fig. 2J | [22] |
| Phage | B1E2_Pavel | Temperate, Abbreviation B1E2 | Fig. 2A, Fig. 2J | [22] |
| Phage | 35_PAlosanto | Temperate, Abbreviation 35 | Fig. 2A, Fig. 2J | [22] |
| Phage | DMS3 | Temperate | Fig. 2A, Fig. 2J | [51] |
| Phage | PilZ25_JBD24and2 | Temperate, Abbreviation PilZ25 | Fig. 2A, Fig. 2J | [22] |
| Phage | PilZ24_JBD24and1 | Temperate, Abbreviation PilZ24 | Fig. 2A, Fig. 2J | [22] |
| Phage | 37_PArmigiano | Temperate, Abbreviation 37 | Fig. 2A, Fig. 2J | [22] |
| Phage | B1A9_JBD69and1 | Temperate, Abbreviation B1A9 | Fig. 2A, Fig. 2J | [22] |
| Phage | 8_PAkora | Temperate, Abbreviation 8 | Fig. 2A, Fig. 2J | [22] |
| Phage | PilZ28_PAncetta | Temperate, Abbreviation PilZ28 | Fig. 2A, Fig. 2J | [22] |
| Phage | K2B8_PAnettone2 | Temperate, Abbreviation K2B8 | Fig. 2A, Fig. 2J | [22] |
| Phage | B3C1_PAnettone1 | Temperate, Abbreviation B3C1 | Fig. 2A, Fig. 2E, Fig. 2J | [22] |
| Phage | B2F4_PAlmier | Temperate, Abbreviation B2F4 | Fig. 2A, Fig. 2E, Fig. 2J | [22] |
| Phage | PilZ27_PApadam | Temperate, Abbreviation PilZ27 | Fig. 2A, Fig. 2J | [22] |
| Phage | B3A6_PAnipuri | Temperate, Abbreviation B3A6 | Fig. 2A, Fig. 2J | [22] |
| Phage | 3_PAneer1 | Temperate, Abbreviation 3 | Fig. 2A, Fig. 2J | [22] |
| Phage | 32_PAneer2 | Temperate, Abbreviation 32 | Fig. 2A, Fig. 2J | [22] |
| Phage | PilZ41_PAvlova | Temperate, Abbreviation PilZ41 | Fig. 2A, Fig. 2J | [22] |

| Type | Name | Description | Use in this study | Source |
|---|---|---|---|---|
| Phage | K2C5_PAncake | Temperate, Abbreviation K2C5 | Fig. 2A, Fig. 2J | [22] |
| Phage | B3B3_PAsta | Temperate, Abbreviation B3B3 | Fig. 2A, Fig. 2E, Fig. 2J | [22] |
| Phage | B1C6_PAratha | Temperate, Abbreviation B1C6 | Fig. 2A, Fig. 2J | [22] |
| Phage | PilZ14_PAte2 | Temperate, Abbreviation PilZ14 | Fig. 2A, Fig. 2J | [22] |
| Phage | PilZ15_PAte1 | Temperate, Abbreviation PilZ15 | Fig. 2A, Fig. 2J | [22] |
| Phage | PilZ13_PAte3 | Temperate, Abbreviation PilZ13 | Fig. 2A, Fig. 2J | [22] |
| Phage | B3C11_PArfait | Temperate, Abbreviation B3C11 | Fig. 2A, Fig. 2J | [22] |
| Phage | PilZ5_PApaya | Temperate, Abbreviation PilZ5 | Fig. 2A, Fig. 2J | [22] |
| Phage | B2G8_PArsnip | Temperate, Abbreviation B2G8 | Fig. 2A, Fig. 2J | [22] |
| Phage | 14c_PAsty1 | Temperate, Abbreviation 14c | Fig. 2A, Fig. 2J | [22] |
| Phage | 14t_PAsty2 | Temperate, Abbreviation 14t | Fig. 2A, Fig. 2J | [22] |
| Phage | B1A7_PAnnacotta | Temperate, Abbreviation B1A7 | Fig. 2A, Fig. 2E, Fig. 2J | [22] |
| Phage | B1B7_PAstrami | Temperate, Abbreviation B1B7 | Fig. 2A, Fig. 2J | [22] |
| Phage | K2C12_PAleale | Temperate, Abbreviation K2C12 | Fig. 2A, Fig. 2J | [22] |
| Phage | K2B6_PAralyzer | Temperate, Abbreviation K2B6 | Fig. 2A, Fig. 2E, Fig. 2J | [22] |
| Phage | B4C5s_PAstis | Temperate, Abbreviation B4C5s | Fig. 2A, Fig. 2J | [22] |
| Phage | PilZ39_PAdme | Temperate, Abbreviation PilZ39 | Fig. 2A, Fig. 2J | [22] |
| Phage | PilZ31_PAddington | Temperate, Abbreviation PilZ31 | Fig. 2A, Fig. 2J | [22] |
| Phage | B1C3_PAcman | Temperate, Abbreviation B1C3 | Fig. 2A, Fig. 2J | [22] |
| Phage | PilZ3c_PAcifc2 | Temperate, Abbreviation PilZ3c | Fig. 2A, Fig. 2J | [22] |
| Phage | PilZ1c_PAcific1 | Temperate, Abbreviation PilZ1c | Fig. 2A, Fig. 2J | [22] |
| Phage | K2D5_PAdstow | Temperate, Abbreviation K2D5 | Fig. 2A, Fig. 2J | [22] |
| Phage | E9_PAragon | Temperate, Abbreviation E9 | Fig. 2A, Fig. 2J | [22] |
| Phage | B2H12_PAignton | Temperate, Abbreviation B2H12 | Fig. 2A, Fig. 2J | [22] |
| Phage | G11_PAllmall | Temperate, Abbreviation G11 | Fig. 2A, Fig. 2E, Fig. 2J | [22] |
| Phage | B4C4s_PAlma | Temperate, Abbreviation B4C4s | Fig. 2A, Fig. 2J | [22] |
| Phage | B2D6_PAnama | Temperate, Abbreviation B2D6 | Fig. 2A, Fig. 2J | [22] |
| Phage | B2B12_PAlermo | Temperate, Abbreviation B2B12 | Fig. 2A, Fig. 2J | [22] |
| Phage | B2C10_PAnda | Temperate, Abbreviation B2C10 | Fig. 2A, Fig. 2J | [22] |
| Phage | B2D10_PArakeet | Temperate, Abbreviation B2D10 | Fig. 2A, Fig. 2J | [22] |
| Phage | B1C8_PArrot | Temperate, Abbreviation B1C8 | Fig. 2A, Fig. 2E, Fig. 2J | [22] |
| Phage | B4C2_PAltus | Temperate, Abbreviation B4C2 | Fig. 2A, Fig. 2J | [22] |
| Phage | B4C11_PAngolin | Temperate, Abbreviation B4C11 | Fig. 2A, Fig. 2J | [22] |
| Phage | B2E7P_PApillon | Temperate, Abbreviation B2E7P | Fig. 2A, Fig. 2J | [22] |

| Type | Name | Description | Use in this study | Source |
|---|---|---|---|---|
| Phage | B4C8c_PArtridge | Temperate, Abbreviation B4C8c | Fig. 2A, Fig. 2J | [22] |
| Phage | B3A1_PAnther1 | Temperate, Abbreviation B3A1 | Fig. 2A, Fig. 2J | [22] |
| Phage | B3D4_PAnther3 | Temperate, Abbreviation B3D4 | Fig. 2A, Fig. 2J | [22] |
| Phage | B2F12_PAsadena | Temperate, Abbreviation B2F12 | Fig. 2A, Fig. 2J | [22] |
| Phage | ThaiE2_PAdthai | Temperate, Abbreviation ThaiE2 | Fig. 2A, Fig. 2J | [22] |
| Phage | K6A8_PAssau1 | Temperate, Abbreviation K6A8 | Fig. 2A, Fig. 2J | [22] |
| Phage | UTIA11_PAssau2 | Temperate, Abbreviation UTIA11 | Fig. 2A, Fig. 2J | [22] |
| Phage | B3H1s_PAphos2 | Temperate, Abbreviation B3H1s | Fig. 2A, Fig. 2J | [22] |
| Phage | B3H1b_PAphos1 | Temperate, Abbreviation B3H1b | Fig. 2A, Fig. 2J | [22] |
| Phage | K2C7c_PAdova | Temperate, Abbreviation K2C7c | Fig. 2A, Fig. 2J | [22] |
| Phage | K7A8c_PArma1 | Temperate, Abbreviation K7A8c | Fig. 2A, Fig. 2J | [22] |
| Phage | K7A8h_PArma2 | Temperate, Abbreviation K7A8h | Fig. 2A, Fig. 2J | [22] |
| Phage | B3B1_PAlmyra | Temperate, Abbreviation B3B1 | Fig. 2A, Fig. 2J | [22] |
| Phage | B1E5_PAkhom | Temperate, Abbreviation B1E5 | Fig. 2A, Fig. 2J | [22] |
| Phage | B3E7_PArtisan | Temperate, Abbreviation B3E7 | Fig. 2A, Fig. 2J | [22] |
| Phage | K2B12c_PAthetic2 | Temperate, Abbreviation K2B12c | Fig. 2A, Fig. 2J | [22] |
| Phage | B1B1b_PAthetic1 | Temperate, Abbreviation B1B1b | Fig. 2A, Fig. 2J | [22] |
| Phage | D12_PAncras1 | Temperate, Abbreviation D12 | Fig. 2A, Fig. 2J | [22] |
| Phage | B1H2s_PAncras4 | Temperate, Abbreviation B1H2s | Fig. 2A, Fig. 2J | [22] |
| Phage | F2_PAncras2 | Temperate, Abbreviation F2 | Fig. 2A, Fig. 2J | [22] |
| Phage | B4C5b_PAncras3 | Temperate, Abbreviation B4C5b | Fig. 2A, Fig. 2J | [22] |
| Phage | B4C8h_PAncras6 | Temperate, Abbreviation B4C8h | Fig. 2A, Fig. 2J | [22] |
| Phage | F3_PAlestra | Temperate, Abbreviation F3 | Fig. 2A, Fig. 2E, Fig. 2J | [22] |
| Phage | K2C7t_PAzuzu | Temperate, Abbreviation K2C7t | Fig. 2A, Fig. 2J | [22] |
| Phage | D6_PAndemonium | Temperate, Abbreviation D6 | Fig. 2A, Fig. 2E, Fig. 2J | [22] |
| Phage | K6G2_PAlazzo1 | Temperate, Abbreviation K6G2 | Fig. 2A, Fig. 2E, Fig. 2J | [22] |
| Phage | B2F2_PAchinko1 | Temperate, Abbreviation B2F2 | Fig. 2A, Fig. 2J | [22] |
| Phage | B3A9_PArseltongue | Temperate, Abbreviation B3A9 | Fig. 2A, Fig. 2E, Fig. 2J | [22] |
| Phage | 43_PAssenger3 | Temperate, Abbreviation 43 | Fig. 2A, Fig. 2J | [22] |
| Phage | 101_PAssenger2 | Temperate, Abbreviation 101 | Fig. 2A, Fig. 2J | [22] |
| Phage | 68_PAstor | Temperate, Abbreviation 68 | Fig. 2A, Fig. 2J | [22] |
| Phage | E8_PAsteur2 | Temperate, Abbreviation E8 | Fig. 2A, Fig. 2J | [22] |
| Phage | B2C9_PAgoda2 | Temperate, Abbreviation B2C9 | Fig. 2A, Fig. 2J | [22] |
| Phage | B2B9_PAgoda1 | Temperate, Abbreviation B2B9 | Fig. 2A, Fig. 2J | [22] |

| Type | Name | Description | Use in this study | Source |
|---|---|---|---|---|
| Phage | B3A5_PAparazzi1 | Temperate, Abbreviation B3A5 | Fig. 2A, Fig. 2J | [22] |
| Phage | B3B5b_PAparazzi3 | Temperate, Abbreviation B3B5b | Fig. 2A, Fig. 2J | [22] |
| Phage | B3G4_PAparazzi4 | Temperate, Abbreviation B3G4 | Fig. 2A, Fig. 2J | [22] |
| Phage | 11 | Temperate | Fig. 2J | [22] |
| Phage | 27 | Temperate | Fig. 2J | [22] |
| Phage | 5 | Temperate | Fig. 2J | [22] |
| Phage | B1D3 | Temperate | Fig. 2J | [22] |
| Phage | B2D3 | Temperate | Fig. 2J | [22] |
| Phage | B2G2 | Temperate | Fig. 2J | [22] |
| Phage | B3B5s | Temperate | Fig. 2J | [22] |
| Phage | B3C4 | Temperate | Fig. 2J | [22] |
| Phage | B3F6 | Temperate | Fig. 2J | [22] |
| Phage | B4C7 | Temperate | Fig. 2J | [22] |
| Phage | K2C8 | Temperate | Fig. 2J | [22] |
| Phage | K2G12 | Temperate | Fig. 2J | [22] |
| Phage | Pil3t | Temperate | Fig. 2J | [22] |
| Phage | DMS3vir | Virulent | Fig. 2A | [52] |
| Phage | UT1 | Virulent | Fig. 2A, Fig. 2J | Alexander Hams, ETH Zürich |
| Phage | P973 | Virulent | Fig. 2A, Fig. 2J | Christian Fitch in Ben Temperton's Lab, Exeter |
| Phage | P947 | Virulent | Fig. 2A, Fig. 2J | Christian Fitch in Ben Temperton's Lab, Exeter |
| Phage | PELP20 | Virulent | Fig. 2A, Fig. 2J | Rosanna Wright in Mike Brockhurst's Lab, Manchester |
| Phage | P71 | Virulent | Fig. 2A, Fig. 2J | Christian Fitch in Ben Temperton's Lab, Exeter |
| Phage | PA1P2 | Virulent | Fig. 2A, Fig. 2J | Rosanna Wright in Mike Brockhurst's Lab, Manchester |
| Phage | PA8P2 | Virulent | Fig. 2A, Fig. 2J | Rosanna Wright in Mike Brockhurst's Lab, Manchester |
| Phage | PA1P3 | Virulent | Fig. 2A, Fig. 2J | Rosanna Wright in Mike Brockhurst's Lab, Manchester |
| Phage | 14-1 | Virulent | Fig. 2A, Fig. 2J | Rosanna Wright in Mike Brockhurst's Lab, Manchester |

| Type | Name | Description | Use in this study | Source |
|---|---|---|---|---|
| Phage | P75 | Virulent | Fig. 2A, Fig. 2J | Christian Fitch in Ben Temperton's Lab, Exeter |
| Phage | PA1P1 | Virulent | Fig. 2A, Fig. 2J | Rosanna Wright in Mike Brockhurst's Lab, Manchester |
| Phage | PA10P2 | Virulent | Fig. 2A, Fig. 2J | Rosanna Wright in Mike Brockhurst's Lab, Manchester |
| Phage | PA1P5 | Virulent | Fig. 2A, Fig. 2J | Rosanna Wright in Mike Brockhurst's Lab, Manchester |
| Phage | PA10P1 | Virulent | Fig. 2A, Fig. 2J | Rosanna Wright in Mike Brockhurst's Lab, Manchester |
| Phage | PA7P2 | Virulent | Fig. 2A, Fig. 2J | Rosanna Wright in Mike Brockhurst's Lab, Manchester |
| Phage | P173 | Virulent | Fig. 2A, Fig. 2J | Christian Fitch in Ben Temperton's Lab, Exeter |
| Phage | P287 | Virulent | Fig. 2A, Fig. 2J | Christian Fitch in Ben Temperton's Lab, Exeter |
| Phage | P277 | Virulent | Fig. 2A, Fig. 2J | Christian Fitch in Ben Temperton's Lab, Exeter |
| Phage | P279 | Virulent | Fig. 2A, Fig. 2J | Christian Fitch in Ben Temperton's Lab, Exeter |
| Phage | P274 | Virulent | Fig. 2A, Fig. 2J | Christian Fitch in Ben Temperton's Lab, Exeter |
| Phage | P950 | Virulent | Fig. 2A, Fig. 2J | Christian Fitch in Ben Temperton's Lab, Exeter |
| Phage | PA12P1 | Virulent | Fig. 2A, Fig. 2J | Rosanna Wright in Mike Brockhurst's Lab, Manchester |
| Phage | PA11P2 | Virulent | Fig. 2A, Fig. 2J, (tested but excluded from Fig. 2E) | Rosanna Wright in Mike Brockhurst's Lab, Manchester |
| Phage | P213 | Virulent | Fig. 2A, Fig. 2J | Christian Fitch in Ben Temperton's Lab, Exeter |
| Phage | P72 | Virulent | Fig. 2A, Fig. 2J | Christian Fitch in Ben Temperton's Lab, Exeter |
| Phage | P162 | Virulent | Fig. 1F, Fig. 2A, Fig. 2E, Fig. 2J | Christian Fitch in Ben Temperton's Lab, Exeter |

| Type | Name | Description | Use in this study | Source |
|---|---|---|---|---|
| Phage | LMA2 | Virulent | Fig. 1F, Fig. 2A, Fig. 2E, Fig. 2J | Rosanna Wright in Mike Brockhurst's Lab, Manchester |
| Phage | P283 | Virulent | Fig. 2A, Fig. 2J | Christian Fitch in Ben Temperton's Lab, Exeter |
| Phage | P281 | Virulent | Fig. 2A, Fig. 2J | Christian Fitch in Ben Temperton's Lab, Exeter |
| Phage | P286 | Virulent | Fig. 2A, Fig. 2J | Christian Fitch in Ben Temperton's Lab, Exeter |
| Phage | P176 | Virulent | Fig. 2A, Fig. 2E, Fig. 2J | Christian Fitch in Ben Temperton's Lab, Exeter |
| Phage | P280 | Virulent | Fig. 2A, Fig. 2J | Christian Fitch in Ben Temperton's Lab, Exeter |
| Phage | P275 | Virulent | Fig. 2A, Fig. 2J | Christian Fitch in Ben Temperton's Lab, Exeter |
| Phage | P284 | Virulent | Fig. 2E | Christian Fitch in Ben Temperton's Lab, Exeter |
| Phage | P285 | Virulent | Fig. 2A, Fig. 2J | Christian Fitch in Ben Temperton's Lab, Exeter |
| Phage | Victoria | Virulent | Fig. 2A, Fig. 2J | Alexander Hams, ETH Zürich |
| Phage | P113 | Virulent | Fig. 2A, Fig. 2J | Christian Fitch in Ben Temperton's Lab, Exeter |
| Phage | P276 | Virulent | Fig. 1F, Fig. 2A, Fig. 2E, Fig. 2J | Christian Fitch in Ben Temperton's Lab, Exeter |
| Phage | P272 | Virulent | Fig. 2A, Fig. 2E, Fig. 2J | Christian Fitch in Ben Temperton's Lab, Exeter |
| Phage | P70 | Virulent | Fig. 2A, Fig. 2J | Christian Fitch in Ben Temperton's Lab, Exeter |
| Phage | phiKZ | Virulent | Fig. 2A, Fig. 2J | Rosanna Wright in Mike Brockhurst's Lab, Manchester |
| Phage | Cassandra | Virulent | Fig. 2A, Fig. 2J | Alexander Hams, ETH Zürich |
| Phage | Ettore | Virulent | Fig. 2A, Fig. 2J | Alexander Hams, ETH Zürich |
| Phage | Deifobo | Virulent | Fig. 2A, Fig. 2J | Alexander Hams, ETH Zürich |

| Type | Name | Description | Use in this study | Source |
|---|---|---|---|---|
| Phage | Paride | Virulent | Fig. 2A, Fig. 2J | Alexander Hams, ETH Zürich |
| Phage | PNM | Virulent | Fig. 2A, Fig. 2J | Rosanna Wright in Mike Brockhurst's Lab, Manchester |
| Phage | LUZ19 | Virulent | Fig. 2A, Fig. 2J | Prof. Rob Lavigne, University of Leuven |
| Phage | P212 | Virulent | Fig. 2A, Fig. 2E, Fig. 2J | Christian Fitch in Ben Temperton's Lab, Exeter |
| Phage | P217 | Virulent | Fig. 2A, Fig. 2J | Christian Fitch in Ben Temperton's Lab, Exeter |
| Phage | P216 | Virulent | Fig. 2A, Fig. 2J | Christian Fitch in Ben Temperton's Lab, Exeter |
| Phage | P214 | Virulent | Fig. 2A, Fig. 2J | Christian Fitch in Ben Temperton's Lab, Exeter |
| Phage | P215 | Virulent | Fig. 2A, Fig. 2J | Christian Fitch in Ben Temperton's Lab, Exeter |
| Phage | P211 | Virulent | Fig. 2A, Fig. 2E, Fig. 2J | Christian Fitch in Ben Temperton's Lab, Exeter |
| Phage | P278 | Virulent | Fig. 2J | Christian Fitch in Ben Temperton's Lab, Exeter |
| Phage | PA2P1 | Virulent | Fig. 2J | Rosanna Wright in Mike Brockhurst's Lab, Manchester |
| Phage | PA5P1 | Virulent | Fig. 2J | Rosanna Wright in Mike Brockhurst's Lab, Manchester |
| Phage | phiKZ_2 | Virulent | Fig. 2J | Rosanna Wright in Mike Brockhurst's Lab, Manchester |
| Plasmid | pLoLa | *E. coli* / *P. aeruginosa* shuttle vector, gentamicin resistance marker | | [10] |
| Plasmid | pLoLa-mCherry | pLoLa + mCherry gene | MANTIS in vivo assays: Fig. 1F, 2A | [10] |
| Plasmid | pLoLa-dCTPase-AleI(1) | pLoLa + dCTP deaminase gene from AU18068 genome with overlapping AZPAE15024 MANTIS and AleI sites | MANTIS in vivo assays | This study |
| Plasmid | pLoLa-AleI(1) | pLoLa with overlapping AZPAE15024 MANTIS and AleI sites | MANTIS in vivo assays | This study |
| Plasmid | pLoLa-AZPAE15024 | pLoLa + AZPAE15024 MANTIS operon | MANTIS in vivo assays: Fig. 1F, 2A | This study |

| Type | Name | Description | Use in this study | Source |
|---|---|---|---|---|
| Plasmid | pLoLa-AZPAE15024- AleI(1) | pLoLa + AZPAE15024 MANTIS operon with overlapping AZPAE15024 MANTIS and AleI sites | MANTIS in vivo assays: Fig. 1E | This study |
| Plasmid | pLoLa-ψ6684 | pLoLa + AZPAE15024 MANTIS operon with MtsC region (aa 444-527) from PSE6684 MANTIS | MANTIS in vivo assays: Fig. 1E, 1F | This study |
| Plasmid | pLoLa-AZPAE15024 ΔmtsA | pLoLa + AZPAE15024 MANTIS operon and ΔmtsA | MANTIS in vivo assays: Fig. 1F | This study |
| Plasmid | pLoLa-AZPAE15024 ΔmtsC | pLoLa + AZPAE15024 MANTIS operon and ΔmtsC | MANTIS in vivo assays: Fig. 1F | This study |
| Plasmid | pLoLa-AZPAE15024- AleI(1) ΔmtsR | pLoLa + AZPAE15024 MANTIS operon with overlapping AZPAE15024 MANTIS and AleI sites and ΔmtsR | MANTIS in vivo assays: Fig. 1E, 1F | This study |
| Plasmid | pLoLa-AZPAE15024- AleI(1) ΔmtsA | pLoLa + AZPAE15024 MANTIS operon with overlapping AZPAE15024 MANTIS and AleI sites and ΔmtsA | MANTIS in vivo assays: Fig. 1E | This study |
| Plasmid | pLoLa-AZPAE15024- AleI(1) ΔmtsB | pLoLa + AZPAE15024 MANTIS operon with overlapping AZPAE15024 MANTIS and AleI sites and ΔmtsB | MANTIS in vivo assays: Fig. 1E, 1F | This study |
| Plasmid | pLoLa-AZPAE15024- AleI(1) ΔmtsC | pLoLa + AZPAE15024 MANTIS operon with overlapping AZPAE15024 MANTIS and AleI sites and ΔmtsC | MANTIS in vivo assays: Fig. 1E | This study |
| Plasmid | pLoLa-AZPAE15024- AleI(1) ΔmtsD | pLoLa + AZPAE15024 MANTIS operon with overlapping AZPAE15024 MANTIS and AleI sites and ΔmtsD | MANTIS in vivo assays: Fig. 1E, 1F | This study |
| Plasmid | pLoLa-AZPAE15024- AleI(1) MtsA K80A | pLoLa + AZPAE15024 MANTIS operon with overlapping AZPAE15024 MANTIS and AleI sites and K80A substitution in MtsA | MANTIS in vivo assays: Fig. 1E, 1F | This study |
| Plasmid | pLoLa-AZPAE15024- AleI(1) MtsA E174A | pLoLa + AZPAE15024 MANTIS operon with overlapping AZPAE15024 MANTIS and AleI sites and E174A substitution in MtsA | MANTIS in vivo assays: Fig. 1E, 1F | This study |
| Plasmid | pLoLa-AZPAE15024- AleI(1) MtsC D621A | pLoLa + AZPAE15024 MANTIS operon with overlapping AZPAE15024 MANTIS and AleI sites and D621A substitution in MtsC | MANTIS in vivo assays: Fig. 1E, 1F | This study |
| Plasmid | pLoLa-AZPAE15024- AleI(1) MtsC Y624A | pLoLa + AZPAE15024 MANTIS operon with overlapping AZPAE15024 MANTIS and AleI sites and Y624A substitution in MtsC | MANTIS in vivo assays: Fig. 1E, 1F | This study |
| Plasmid | pLoLa-AZPAE15024- AleI(1) MtsC D621A MtsC Y624A MtsD H122A | pLoLa + AZPAE15024 MANTIS operon with overlapping AZPAE15024 MANTIS and AleI sites, D621A and Y624A substitutions in MtsC, H122A substitution in MtsD | MANTIS in vivo assays: Fig. 1E, 1F | This study |
| Plasmid | pLoLa-AZPAE15024- AleI(1) MtsD H122A | pLoLa + AZPAE15024 MANTIS operon with overlapping AZPAE15024 MANTIS and AleI sites and H122A substitution in MtsD | MANTIS in vivo assays: Fig. 1E, 1F | This study |
| Plasmid | pLoLa-AZPAE15024- AleI(1) MtsD K260A | pLoLa + AZPAE15024 MANTIS operon with overlapping AZPAE15024 MANTIS | MANTIS in vivo assays: Fig. 1E, 1F | This study |

| Type | Name | Description | Use in this study | Source |
|---|---|---|---|---|
| | | and AleI sites and K260A substitution in MtsD | | |
| Plasmid | pEHisV5TEV | T7 promoter protein expression vector | Protein expression | [49] |
| Plasmid | pEV5HisTev Mts A | pEHisV5TEV expressing WT MtsA | Protein expression | This study |
| Plasmid | pEV5HisTev Mts B | pEHisV5TEV expressing WT MtsB | Protein expression | This study |
| Plasmid | pEV5HisTev Mts C | pEHisV5TEV expressing WT MtsC | Protein expression | This study |
| Plasmid | pEV5HisTev Mts D | pEHisV5TEV expressing WT MtsD | Protein expression | This study |
| Plasmid | pEV5HisTev Mts A (K80A)) | pEHisV5TEV expressing MtsA (K80A) | Protein expression | This study |
| Plasmid | pEV5HisTev Mts A (E174A) | pEHisV5TEV expressing MtsA (K174A) | Protein expression | This study |
| Plasmid | pUC19 | High-copy-number *E. coli* cloning vector, ampicillin-resistance marker | | [48] |
| Plasmid | pSJS1 | pUC19 with MANTIS/AleI site | Fig. 3A-D | This study |
| Plasmid | pLoLa-SJS1 | AZPAE15024 MANTIS 1-site derived by cloning pSJS1 target region into pLoLa | Fig. 2I | This study |
| Plasmid | pLoLa-SJS0 | pLoLa-SJS1 variant with zero AZPAE15024 MANTIS sites | Fig. 2I | This study |
| Plasmid | pLoLa-SJS2a | pLoLa-SJS1 variant with 2 AZPAE15024 MANTIS sites in inverted repeat | Fig. 2I | This study |
| Plasmid | pLoLa-SJS2b | pLoLa-SJS1 variant with 2 AZPAE15024 MANTIS sites in direct repeat | Fig. 2I | This study |
| Plasmid | pLoLa-SJS3a | pLoLa-SJS1 variant with 3 AZPAE15024 MANTIS sites in inverted repeat | Fig. 2I | This study |
| Plasmid | pLoLa-SJS3b | pLoLa-SJS1 variant with 3 AZPAE15024 MANTIS sites in direct repeat | Fig. 2I | This study |
| Plasmid | pLoLa-SJS4a | pLoLa-SJS1 variant with 4 AZPAE15024 MANTIS sites in inverted repeat | Fig. 2I | This study |
| Plasmid | pLoLa-SJS4b | pLoLa-SJS1 variant with 4 AZPAE15024 MANTIS sites in direct repeat | Fig. 2I | This study |
| Plasmid | pLoLa-SJS10a | pLoLa-SJS1 variant with 10 AZPAE15024 MANTIS sites in inverted repeat | Fig. 2I | This study |
| Plasmid | pLoLa-SJS10b | pLoLa-SJS1 variant with 10 AZPAE15024 MANTIS sites in direct repeat | Fig. 2I | This study |
