## Supplementary Figures for "Dual ATPase-activated DNA hemimethylation and cleavage by the MANTIS defence system"

*Communicating authors

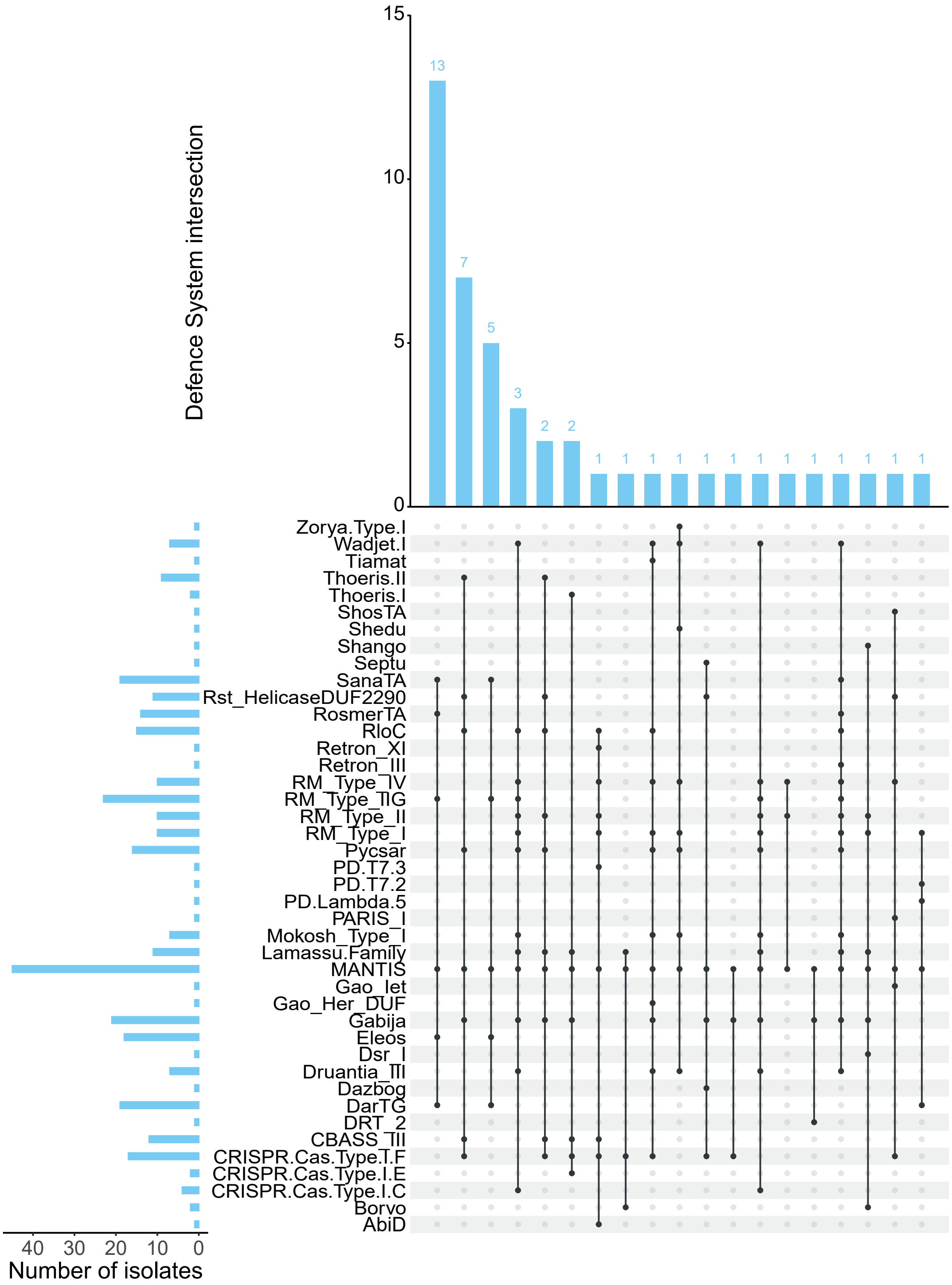


**Supplementary Figure 1 | UpSet plot showing the co-occurrence patterns of defence systems among *P. aeruginosa* isolates encoding MANTIS (n=45).** Each row in the matrix represents a defence system. Filled circles indicate the presence of that system within a given intersection. Connected circles denote defence systems that co-occur within the same set of isolates. The bar chart above the matrix shows the number of MANTIS positive isolates exhibiting each exact combination of defence systems, while the bar chart on the left indicated the total number of MANTIS positive isolates containing each defence system individually. Only defence systems present in at least one MANTIS positive isolate are shown.

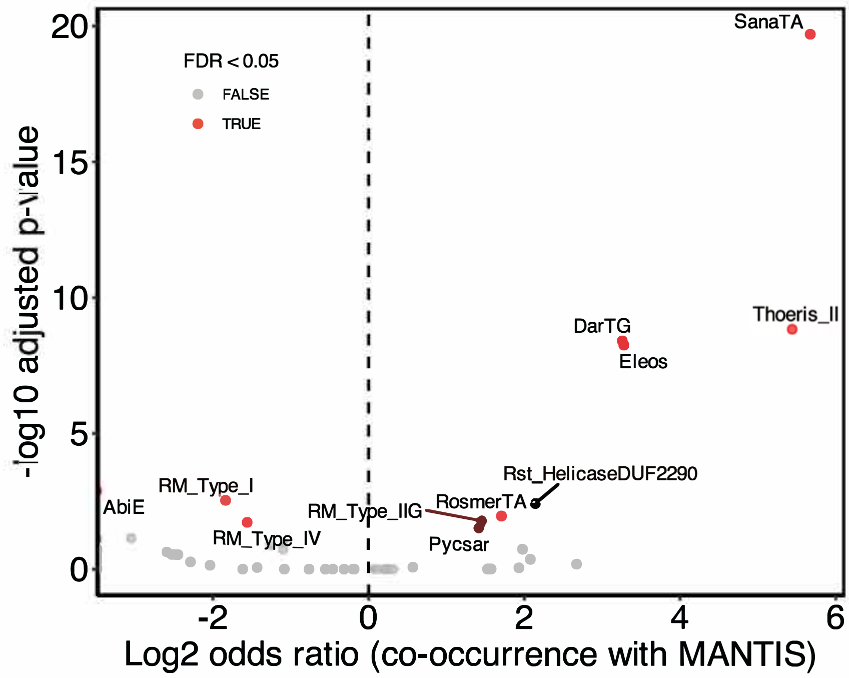


**Supplementary Figure 2 | Volcano plot showing the co-occurrence between bacterial defence systems and the presence of MANTIS across *P. aeruginosa* isolates.** Each point represents one defence system. The *x*-axis shows the $\log_2$-transformed odds ratio from Fisher's exact tests, indicating the strength and direction of association with MANTIS: positive values show enrichment in isolates containing MANTIS; negative values show depletion in isolates containing MANTIS. The *y*-axis shows the $-\log_{10}$ of the Benjamin-Hochberg false discovery rate (FDR)-adjusted p-value. Defence systems significantly associated with MANTIS after multiple testing correction (FDR < 0.05) are highlighted in red and labelled.

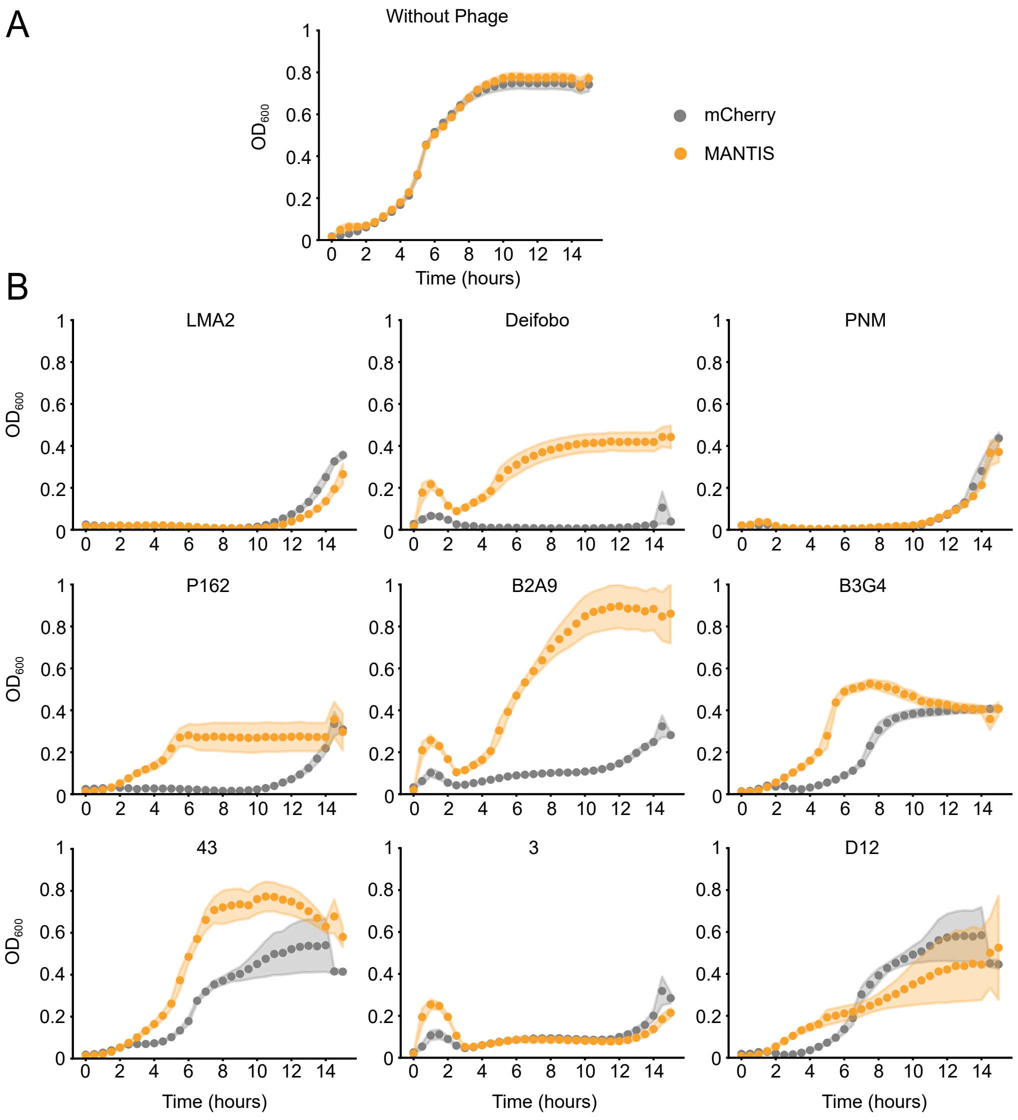


**Supplementary Figure 3 | Example growth curves from liquid culture infection assays.** (**A**) Growth curves of strains carrying mCherry or MANTIS on pLoLa plasmid without phage. (**B**) Growth curves of the same strains during infection with the indicated phages. Lines show mean $OD_{600}$ over time; shaded regions indicate error across biological replicates.

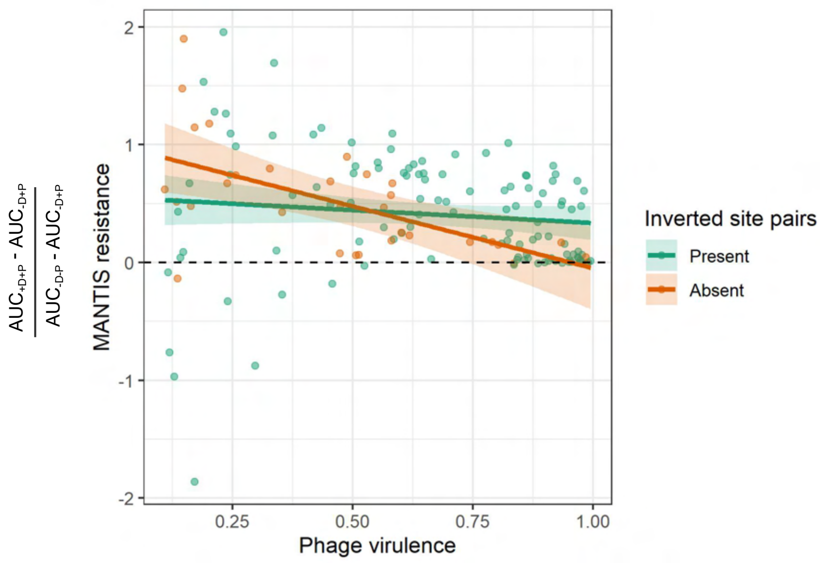


**Supplementary Figure 4 | Correlation between MANTIS resistance and phage virulence.** MANTIS resistance is independent of phage virulence when phages encode inverted site pairs, but for phages lacking them, protection weakens as virulence increases. Points represent phage means for both MANTIS resistance and phage virulence, from 3 biological replicates. Lines denote linear model predictions, and ribbons 95% confidence intervals of the prediction.

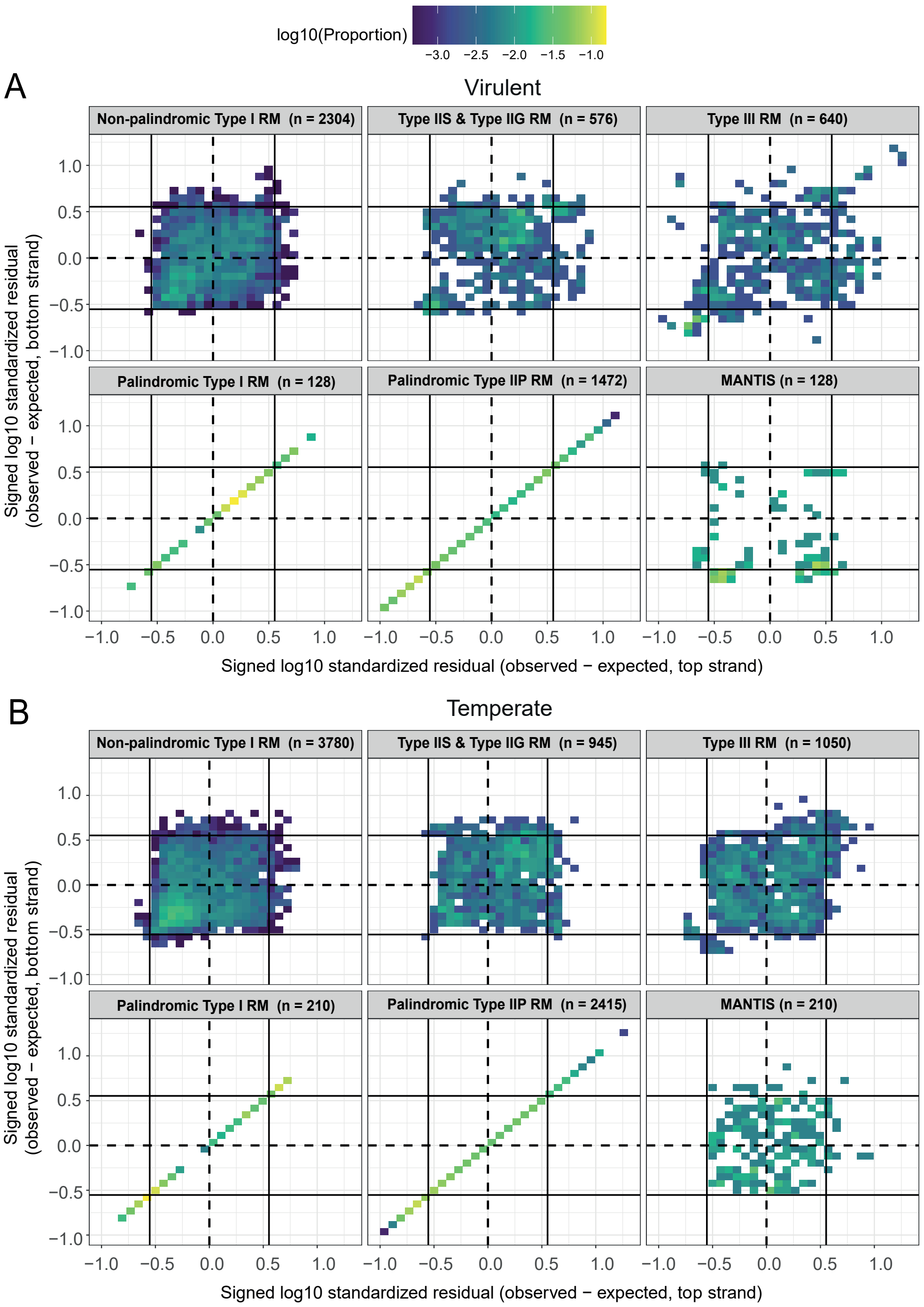


**Supplementary Figure 5 | Strand-specific RM and MANTIS DNA recognition site avoidance in *Pseudomonas* phages.** Density heatmaps indicating the $\log_{10}$-transformed standardised residuals of observed motif counts relative to expectation, calculated independently for forward and reverse strands, for known recognition motifs of *P. aeruginosa* RM systems (separated by system type) and MANTIS, across all virulent (**A**) and temperate phages (**B**). Positive values indicate enrichment of the motif relative to the null expectation, while negative values indicate depletion. Solid horizontal and vertical lines at ±0.553 denote the 99% confidence limits under the null model, residual values exceeding these thresholds are considered significantly different from expectation for the respective strand. Heatmap tile colour indicates the density of values, standardised to the total number of phage-motif pair values in a panel, given with n.

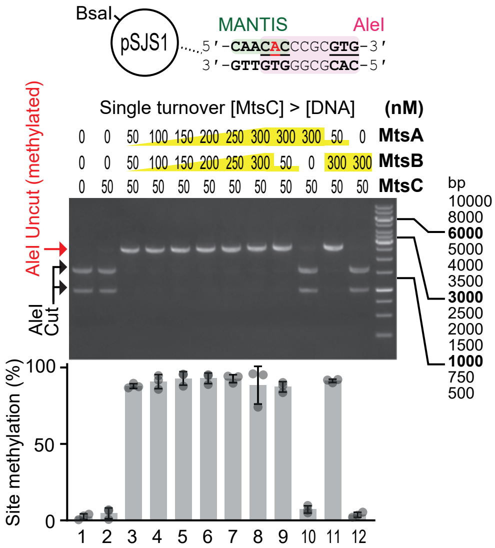


**Supplementary Figure 6 | Dependence on *in vitro* single-turnover DNA methylation of MANTIS subunits.** *In vitro* methylation by MtsC requires both MtsA and MtsB under single turnover conditions. As indicated, 50-300 nM MtsA and/or MtsB were incubated with 50 nM MtsC and 5 nM pSJS1 for 30 minutes with ATP and AdoMet. Following incubation, methylation was quantified by digestion with a combination of AleI and BsaI.

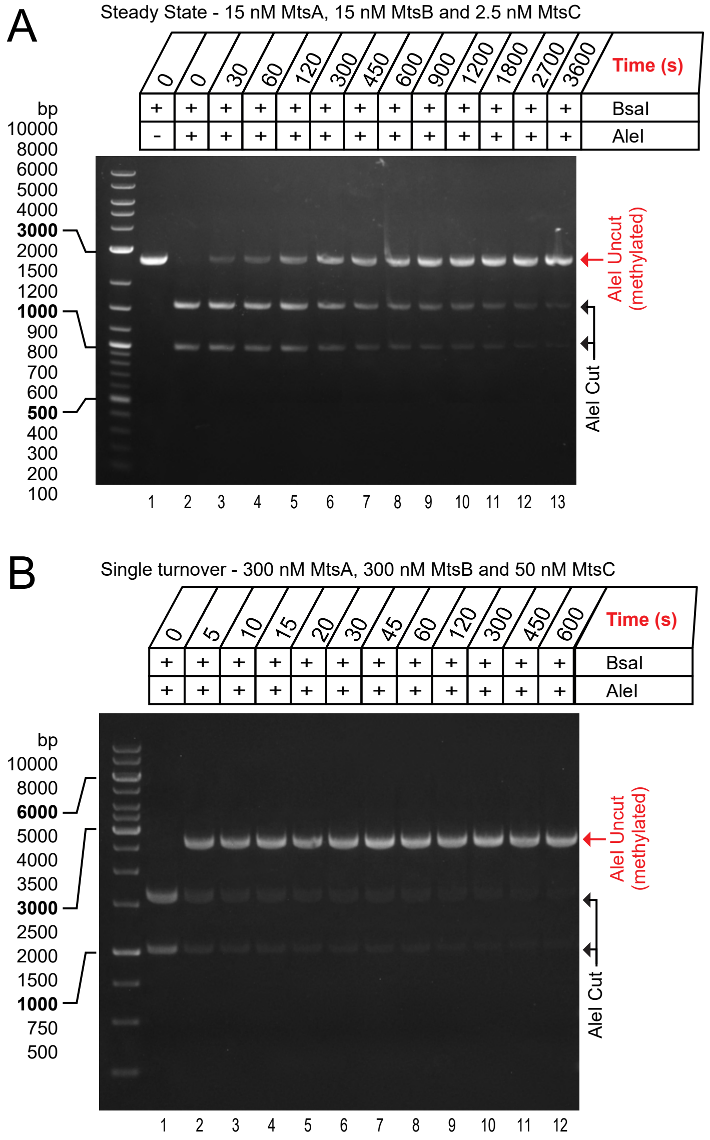


**Supplementary Figure 7 | Measurement of rates of DNA methylation.** *In vitro* time course assays under steady state and single turnover conditions. 5 nM pSJS1 was incubated with MtsABC under steady state (**A**) or single turnover (**B**) conditions, concentrations indicated, with ATP and AdoMet. Samples were taken at time points and quenched with EDTA before purification and quantification by AleI and BsaI digestion.

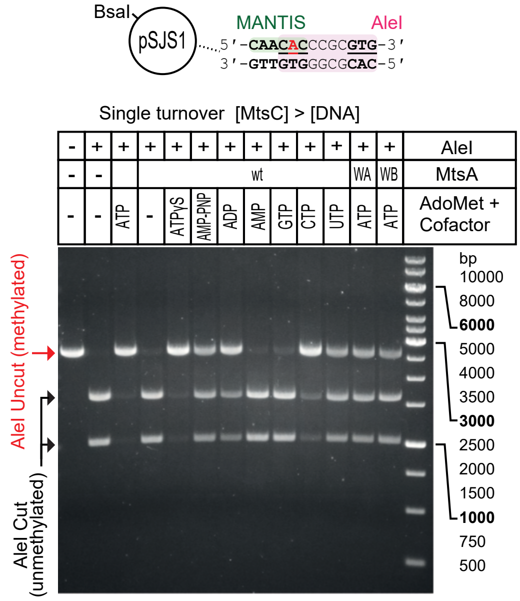


**Supplementary Figure 8 | Measurement of the effects of different nucleotides and MtsA ATPase mutants on single turnover DNA methylation.** Under single turnover conditions, nucleotide requirement for methylation is flexible. Methylation reactions were incubated containing 5 nM pSJS1, 300 nM MtsA (wt, walker A mutant (WA) or walker B mutant (WB) as indicated), 300 nM MtsB, 50 nM MtsC, AdoMet and nucleotide, where indicated. Following a 30 min incubation, methylation was quantified by AleI and BsaI digestion.

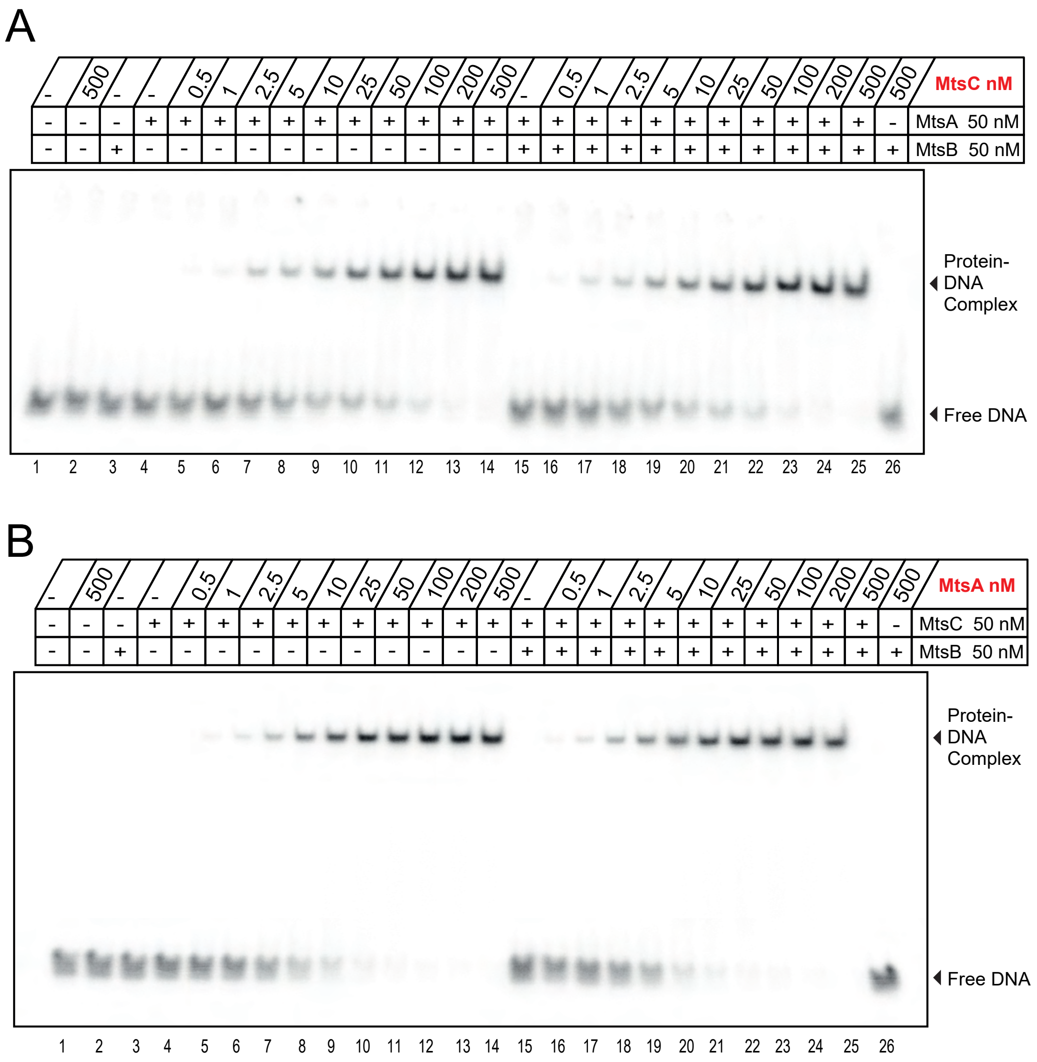


**Supplementary Figure 9 | Measurement of MtsC and MtsA binding affinities by EMSA.** protein-DNA complexes were measured for MtsAC or MtsABC with titrations of MtsC (A) or MtsA (B), respectively. Following a 20 min incubation at room temperature, reactions were separated by EMSA and quantified. (**A**) 0.5 nM $^{32}$P-labelled 50-mer duplex (**Fig. 3E**) was incubated with or without 50 nM MtsA and/or 50 nM MtsB with MtsC titration from 0.5-500 nM, as indicated. (**B**), DNA was incubated with or without 50 nM MtsB and/or 50 nM MtsC with MtsA titration from 0.5-500 nM, as indicated.

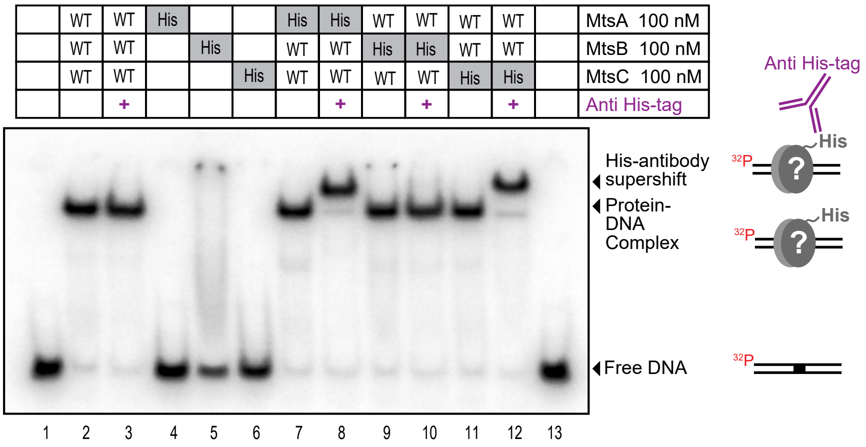


**Supplementary Figure 10 | MtsA and MtsC form a complex on DNA.** Super-shift EMSA experiments with 0.5 nM $^{32}$P-labelled duplex incubated with combinations of 100 nM MtsA, MtsB or MtsC with or without a $(His)_8$-tag, as indicated ("His"). After a 10 min incubation with AdoMet at room temperature, 1 µg Anti His-tag was added, where specified (+), followed by a further 10 min incubation before reactions were separated by EMSA.

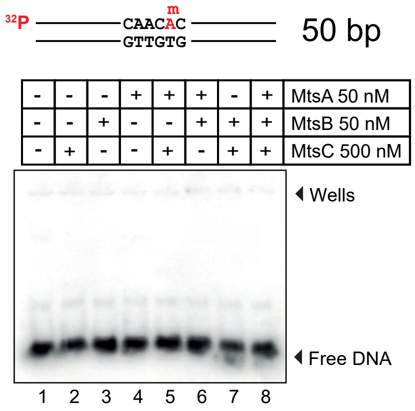


**Supplementary Figure 11 | MtsA, MtsB and MtsC do not form stable interactions with methylated DNA.** EMSA experiments with 0.5 nM $^{32}$P-labelled methylated duplex (equivalent to that in **Fig. 3E**) incubated with 50 nM MtsA/B and 500 nM MtsC, as indicated. Following a 20 min incubation with AdoMet at room temperature, reactions were separated by EMSA.

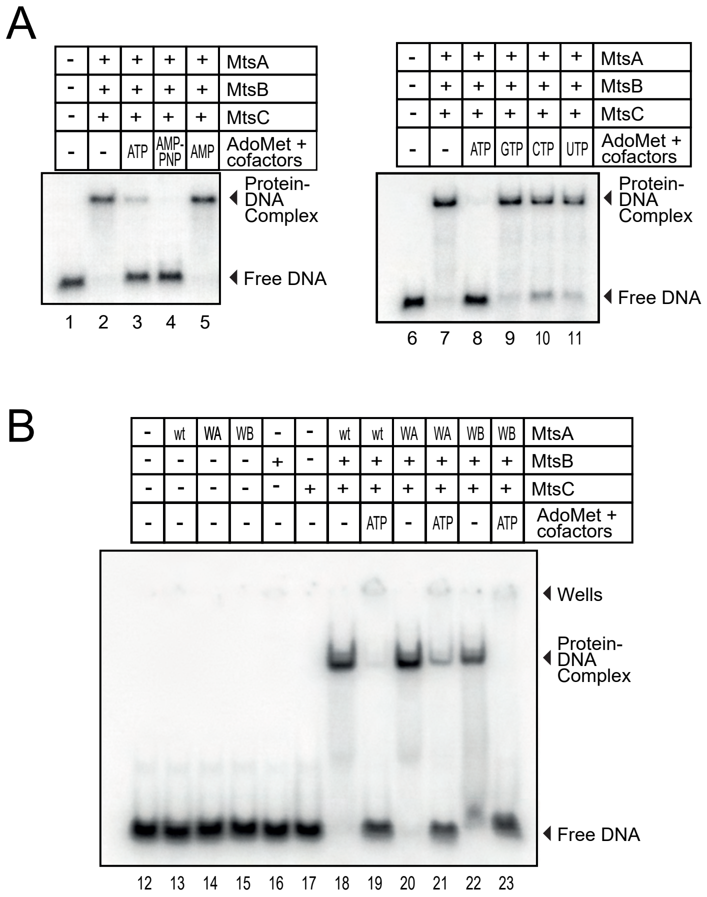


**Supplementary Figure 12 | Measurement of the effects of different nucleotides and MtsA ATPase mutants on DNA binding by EMSA. (A)** Protein-DNA complex displacement is nucleotide dependent. 0.5 nM $^{32}$P-labelled duplex was mixed with 200 nM MtsA, MtsB, MtsC and AdoMet before the addition of cofactors, where indicated. After incubation at room temperature for 20 minutes, reactions were separated by EMSA and quantified (**Fig. 3D**). **(B)** ATP-dependent Protein-DNA complex displacement is observed with MtsA Walker A and B mutants. Combinations of 200 nM MtsA wild-type (wt), Walker A mutant (WA) or Walker B mutant (WB), as indicated, were mixed with 200 nM MtsB and MtsC, and incubated with 0.5 nM $^{32}$P-labelled duplex, AdoMet and ATP, as indicated. Following a 20 min incubation at room temperature, samples were separated by EMSA and quantified (**Fig. 3D**).

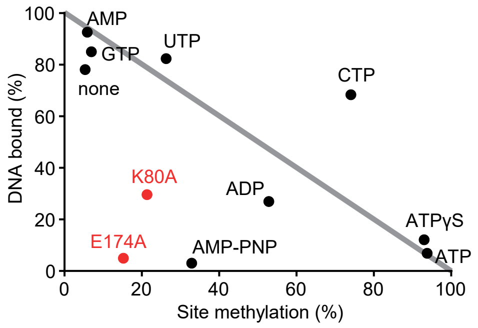


**Supplementary Figure 13 | Correlation between DNA binding and methylation.** Average values (n = 3) taken from **Fig. 3D.** Grey line represents a direct correlation between DNA methylation and the amount of DNA binding.

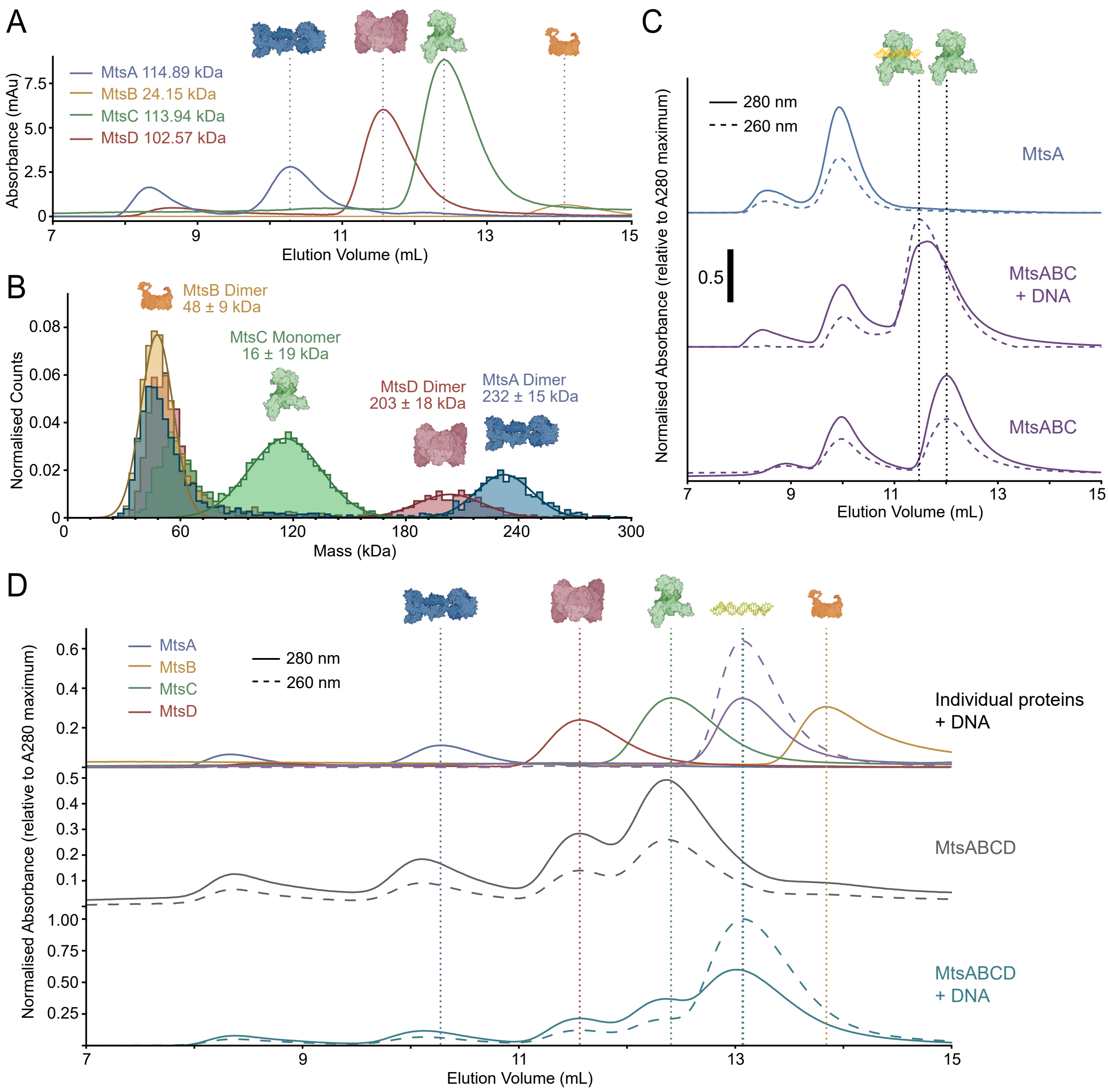


**Supplementary Figure 14 | Measurement of stoichiometries of individual proteins, protein mixtures and protein-DNA complexes.** (**A**) Analytical size exclusion chromatography (SEC) of individual proteins. Theoretical monomer masses indicated. (**B**) Mass photometry analyses of individual proteins fitted to Gaussians with peaks as molecular weight with SD error. The closest stoichiometry is indicated. (**C**) SEC showing loading of MtsC onto a 46 bp oligoduplex by MtsA with MtsB, ATPɣS and sinefungin. Traces are normalised to the maximum absorbance at 280 nM. Bar represents 0.5. (**D**) SEC showing MtsA, MtsB, MtsC and MtsD, individually (top) and in combination (middle and bottom), with and without addition of a 46 bp dsDNA, as indicated. Traces are normalised to the maximum absorbance at 280 nM.

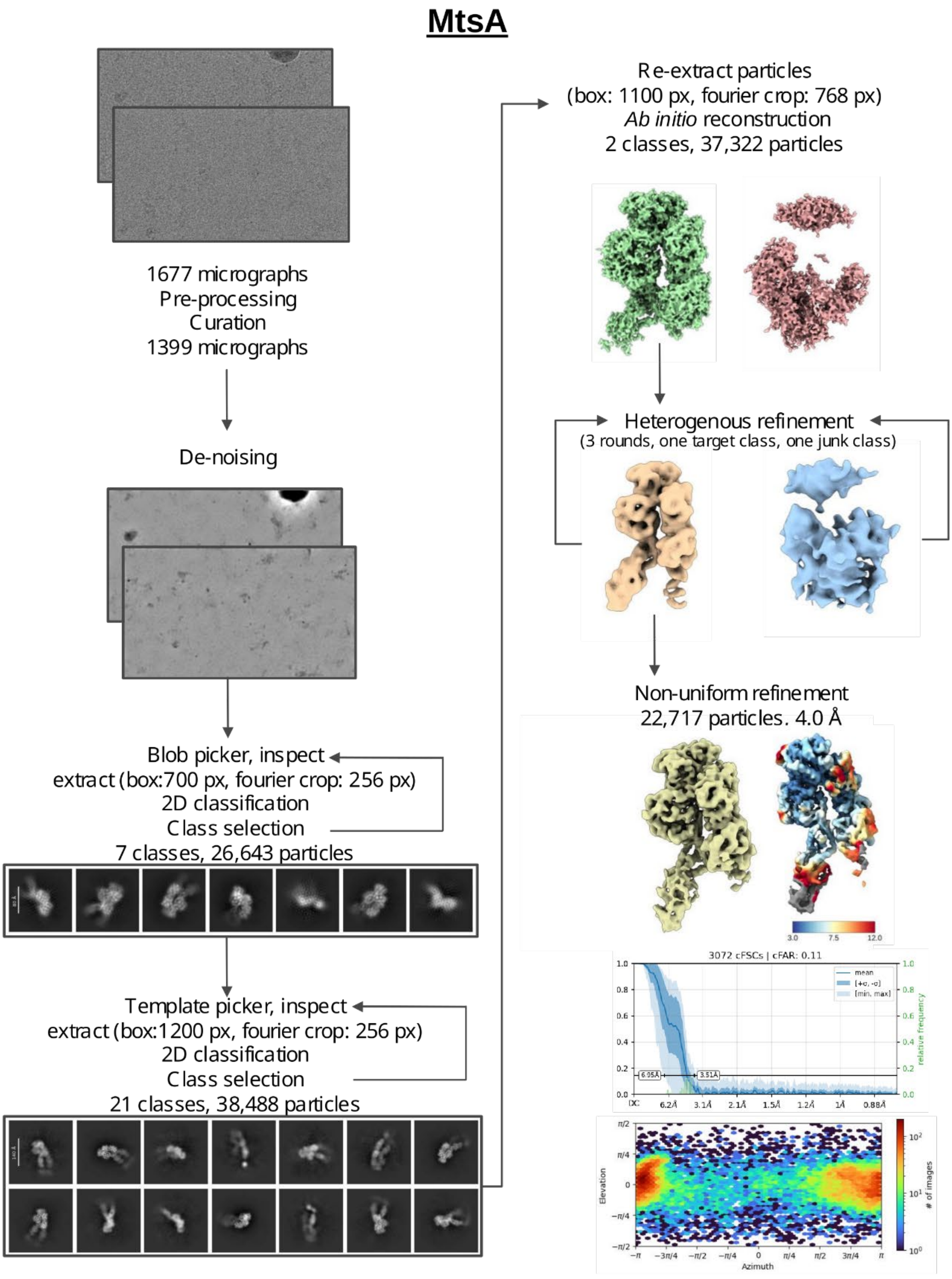


**Supplementary Figure 15 | Cryo-electron microscopy workflow for MtsA.** Data processing flow chart for single particle cryo-EM analysis of dimeric MtsA. Example micrographs and 2D classes are shown along with working and final MtsA maps. Local resolution, GSFCS resolution and orientation analysis are shown for the final map.

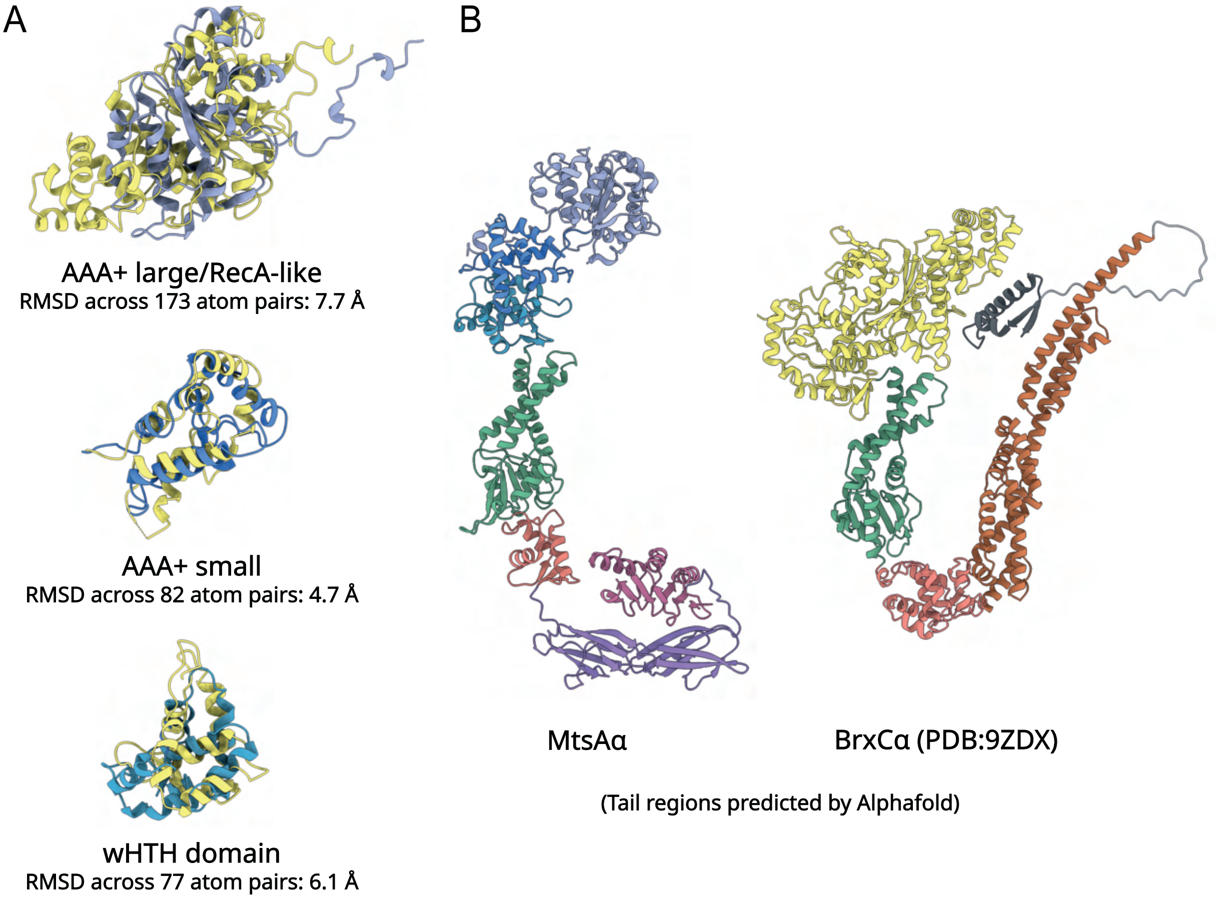


**Supplementary Figure 16 | MtsA shares an overall structural architecture with BrxC but both have distinct C-terminal tail domains.** (**A**) Structural alignments of N-terminal domains of MtsAα and *Acinetobacter* BrxCα (PDB:9ZDX)[15]. (**B**) Comparison of full-length MtsAα and BrxCα (PDB:9ZDX). The N-terminal AAA+ ATPase for MtsA and BrxC are the experimentally derived structures (this study and PDB:9ZDX). Tail regions show a single chain of AlphaFold3 models of full-length dimers. AlphaFold3 models are superimposed onto the respective wHTH domains. Tail domains are coloured to show corresponding/divergent domains in each protomer.

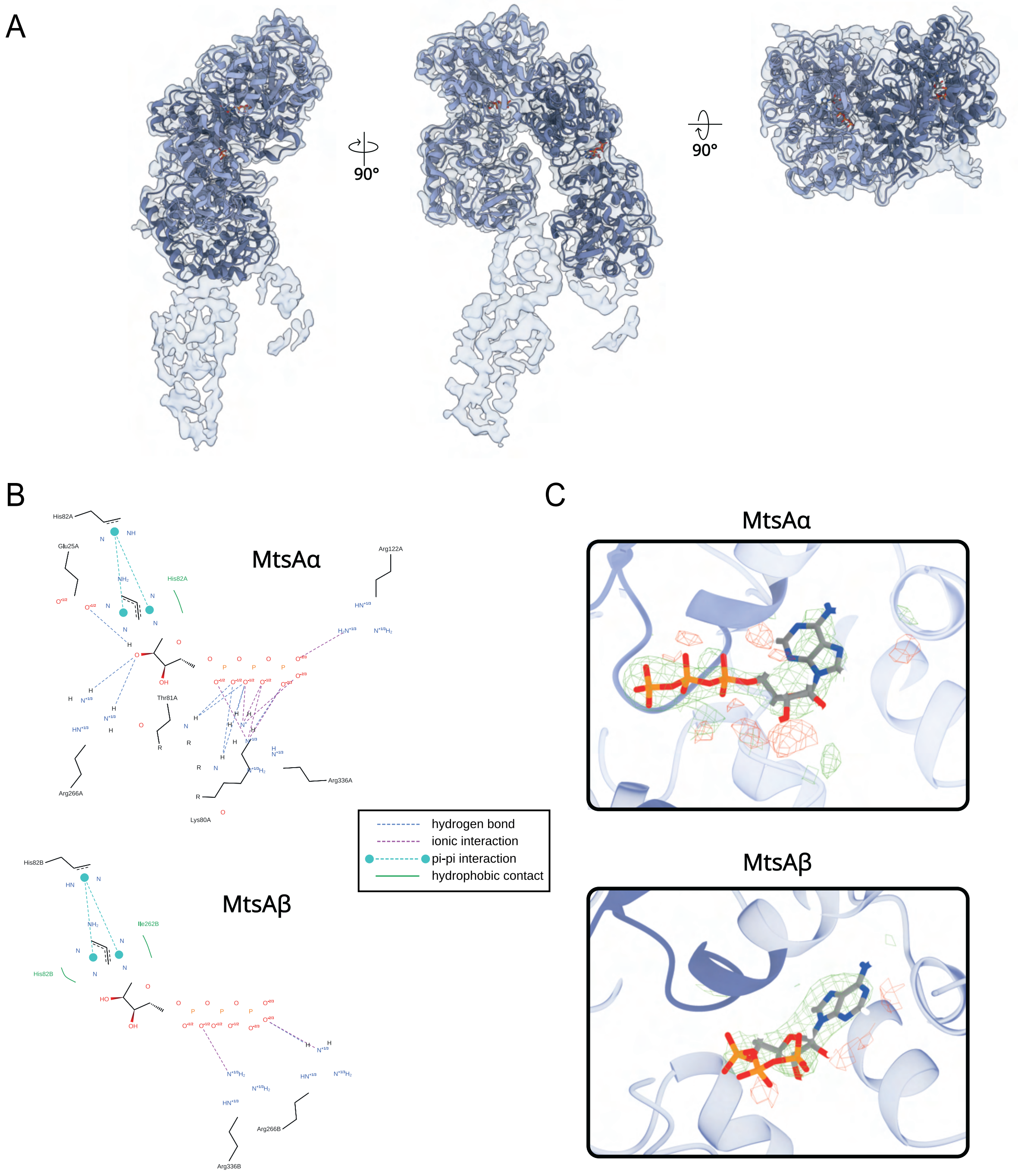


**Supplementary Figure 17 | Analysis of the MtsA structure.** (**A**) MtsA cryo-EM model fitted to experimentally determined density map with rotated views. (**B**) Ligand-residue contacts for ATP binding in MtsAα (left) and MtsAβ (right) subunits, respectively, as calculated by PoseView. (**C**) Polder omit density maps (3 σ) for ATP in MtsAα (top) and MtsBβ (bottom) subunits, respectively.

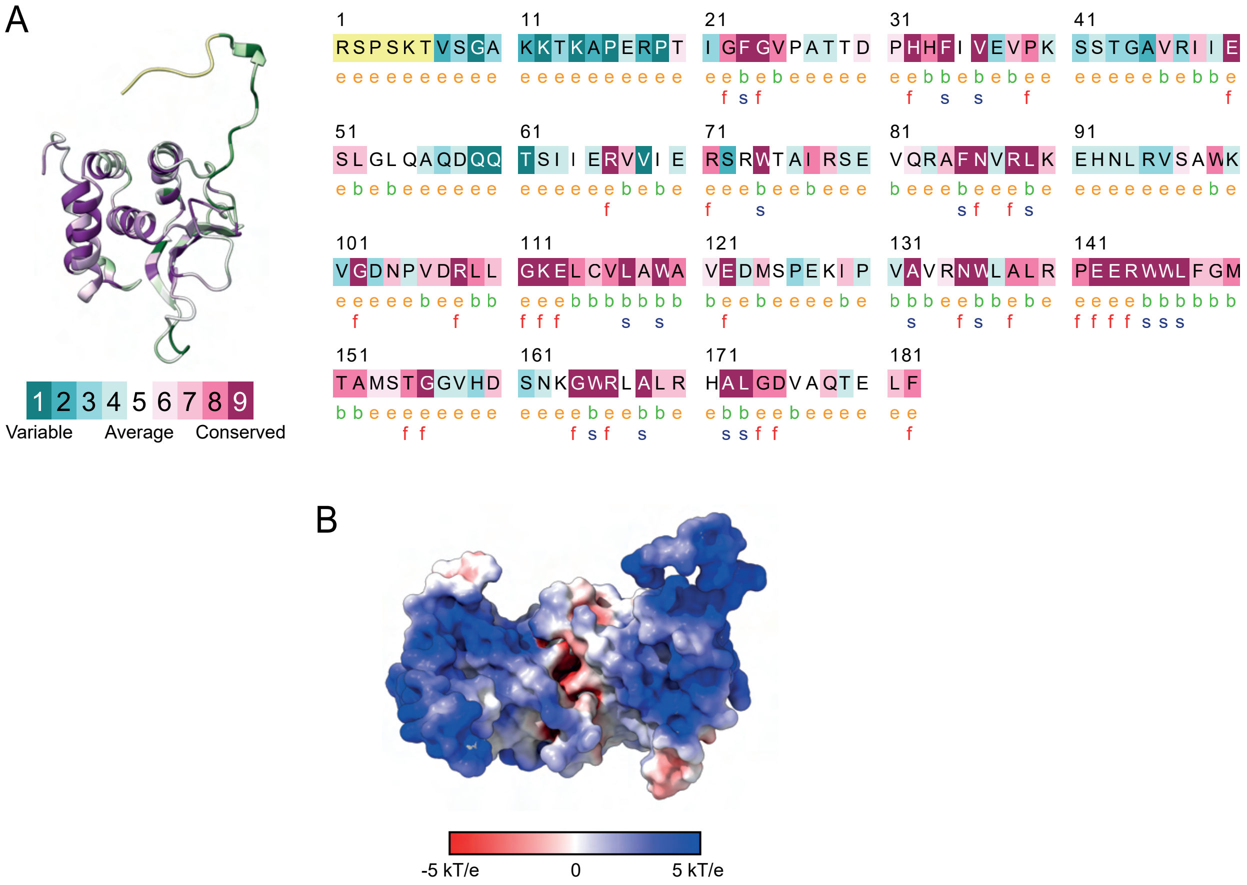


**Supplementary Figure 18 | Analysis of the MtsB structure.** (**A**) Conservation analysis (Consurf) of MtsBα. The dimer interface residues are most highly conserved. (**B**) Surface charge analysis of the MtsB dimer (APBS).

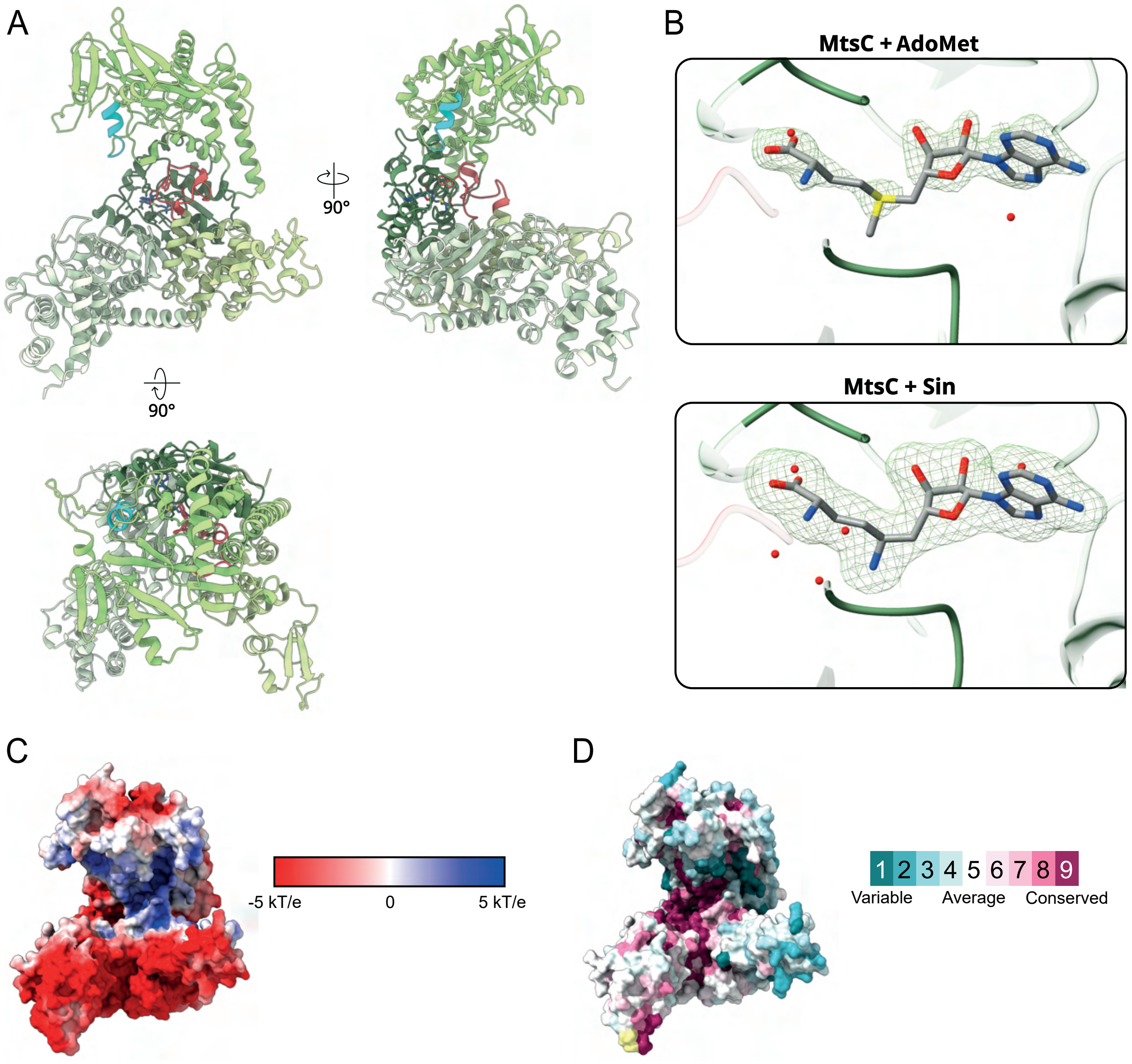


**Supplementary Figure 19 | Analysis of the MtsC structure.** (**A**) Rotated views of the MtsC crystal structure (colouring as in **Fig. 4C**. (**B**) Polder omit density maps (3 σ) for AdoMet and sinefungin ligands in the respective co-crystal structures. (**C**) Surface charge analysis of MtsC (APBS). The DNA binding channel around the conserved methyltransferase core shows an overall positive charge. (**D**) Conservation analysis (Consurf) of MtsC.

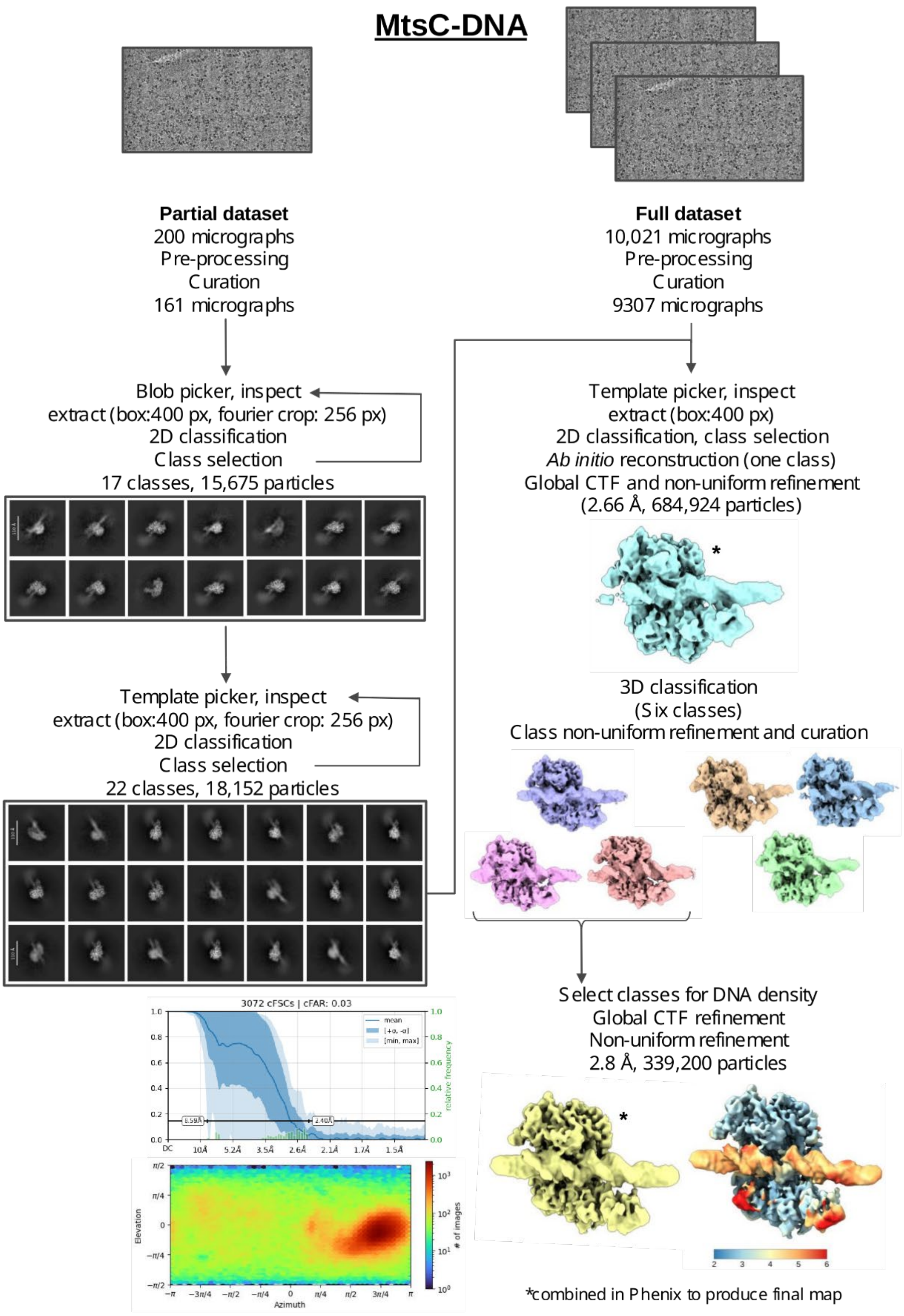


**Supplementary Figure 20 | Cryo-electron microscopy workflow for MtsC-DNA.** Data processing flow chart for single particle cryo-EM analysis of MtsC-DNA. Example micrographs and 2D classes are shown along with working and final MtsC-DNA maps. Local resolution, GSFCS resolution and orientation analysis are shown for the final map.

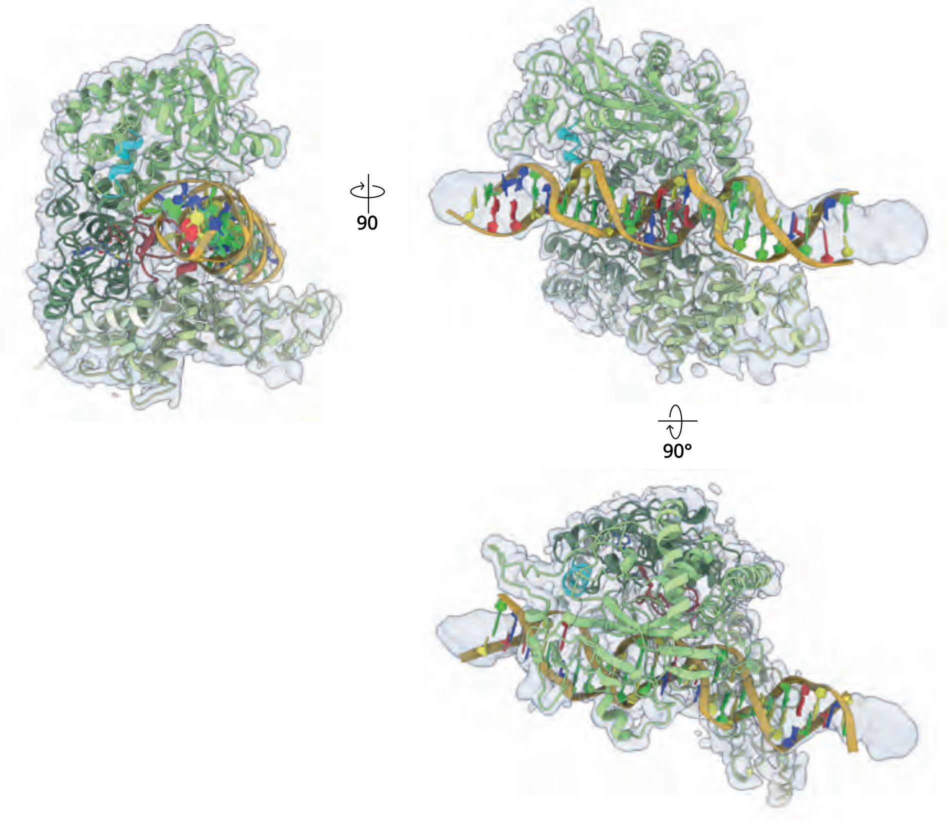


**Supplementary Figure 21 | Analysis of the MtsC-DNA structure.** Rotated views of the MtsC-DNA cryo-EM model fitted to the experimentally derived density map.

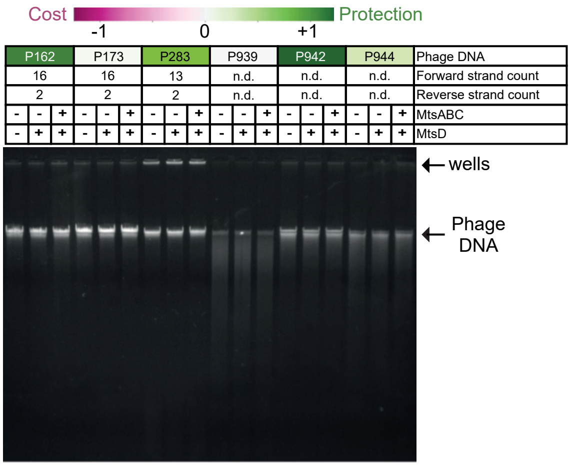


**Supplementary Figure 22 | Phage DNA is not cleaved *in vitro* using MtsABCD.** DNA was purified from various sensitive and resistant phages from our panel (colours indicate activity in the liquid growth curves) and 400 ng was incubated for 1 h at 37 °C with 75 nM MstA-D with ATP, AdoMet and a rCutsmart buffer (NEB), as indicated. Products consistent with DNA cleavage were not observed with any of the phage DNAs

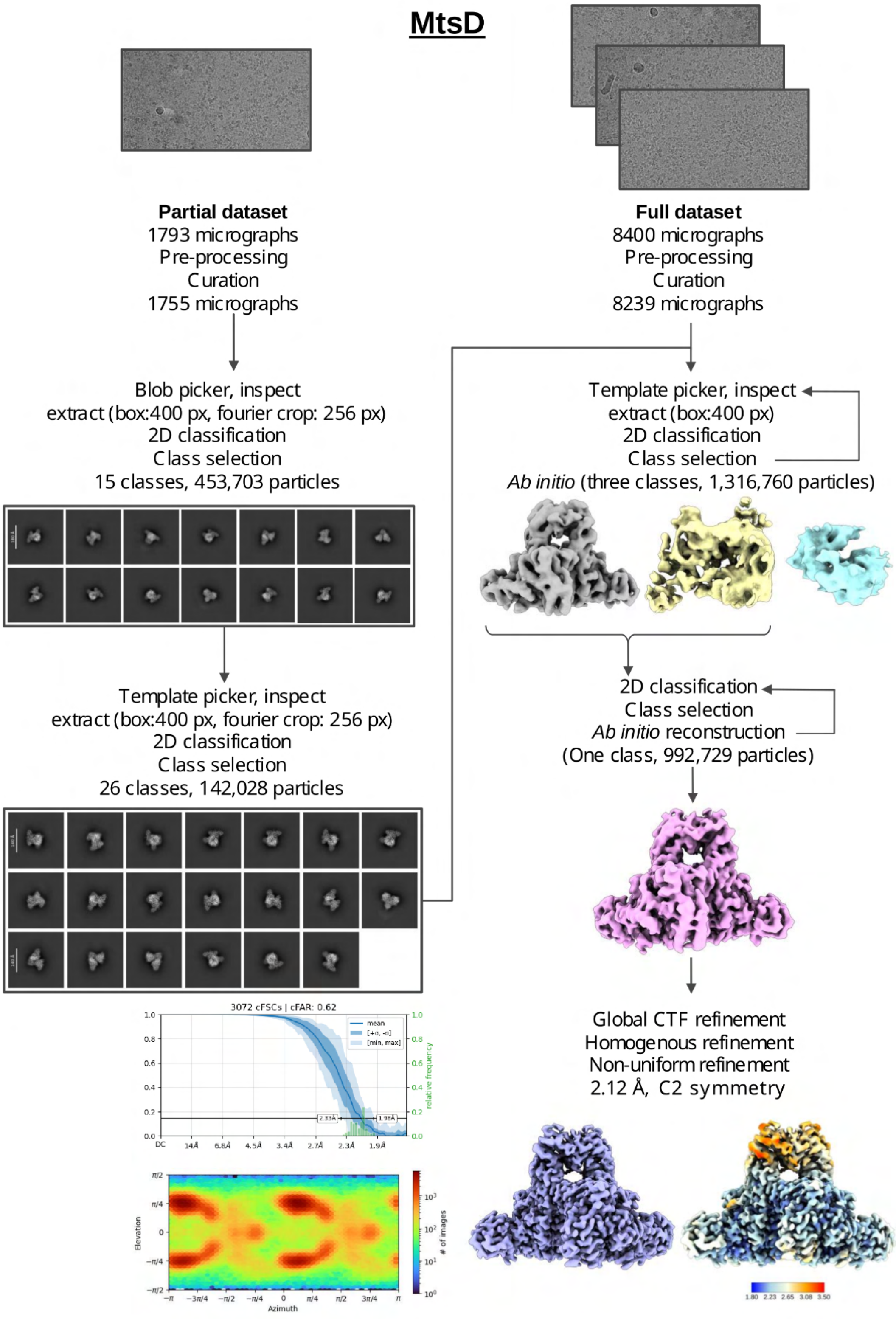


**Supplementary Figure 23 | Cryo-electron microscopy workflow for MtsD.** Data processing flow chart for single particle cryo-EM analysis of dimeric MtsD. Example micrographs and 2D classes are shown along with working and final MtsD maps. Local resolution, GSFCS resolution and orientation analysis are shown for the final map.

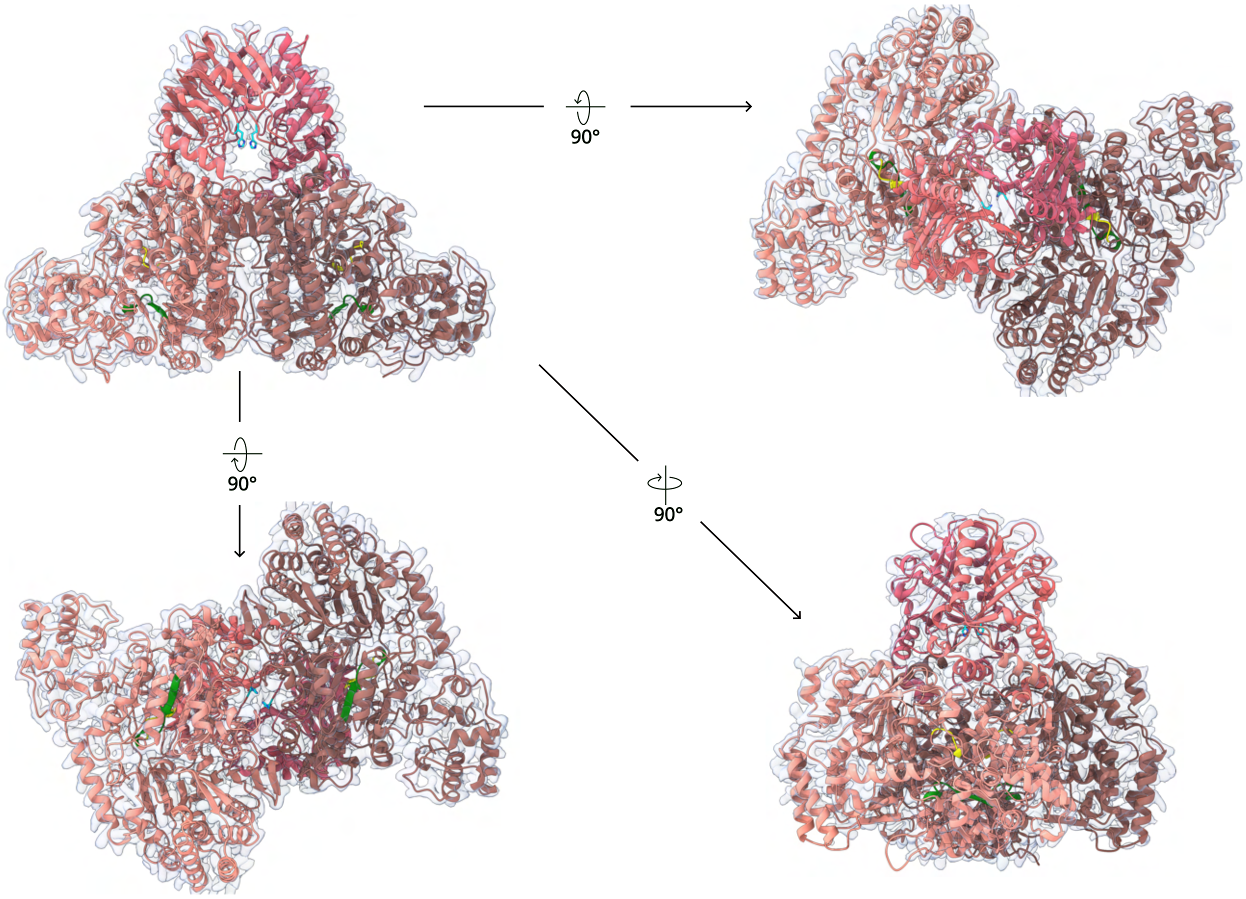


**Supplementary Figure 24 | Analysis of the MtsD structure.** Rotated views of the MtsD cryo-EM structure fitted to the experimentally derived density map. Each protomer is shaded to indicate nuclease and ATPase domains. The catalytic residue H122 is highlighted in cyan. Walker A and Walker B motifs are highlighted in yellow and green, respectively.

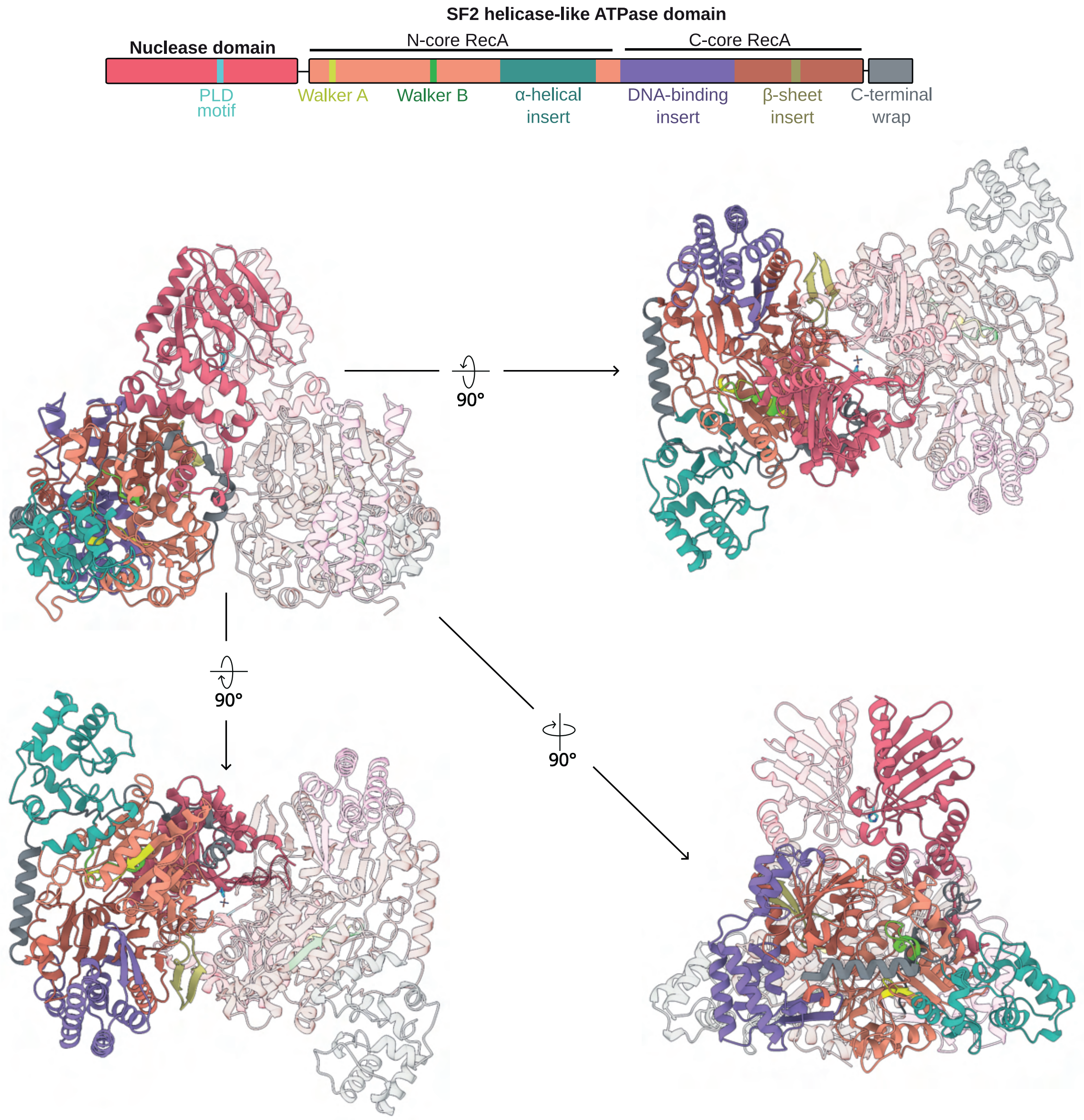


**Supplementary Figure 25 | Relative locations of domains in the MtsD structure.** MtsD protomer coloured by domains and motifs. The domain map above shows the relative positions of domains, motifs and domain insertions
